# Killer-cell dominance dichotomy governs tumor immune networks and stratifies inflamed cancers

**DOI:** 10.64898/2026.06.15.732326

**Authors:** Anlin Li, Zhixin Yu, Wenda Zhang, Hanming Chang, Sirui Feng, Xuan Yang, Kangqiao Xiong, Lina He, Zerui Zhao, Lujun Shen, Zihui Tan, Wei Du, Leqi Zhong, Xu Zhang, Yi Hu, Xiaodong Su, Ruiping Wang, Sha Fu, Li Zhang, Shaodong Hong

## Abstract

Cancer immunotherapy benefits remain limited, even among “hot” tumors with high killer lymphocyte infiltration. Here, we investigated the population-level architecture of killer cells based on nearly 5,000 pan-cancer scRNA-seq samples, together with orthogonal validation by spectral cytometry and spatial transcriptomics. Unlike the prevailing “hot-cold” paradigm, which assumes coordinated infiltration of multiple cytotoxic lineages, we uncovered a conserved framework wherein terminal cytotoxic immunity in individuals or malignancies diverges into states dominated by either exhausted CD8+ T cells (Tex) or CD56^dim^CD16^hi^ NK (NK1) cells. Despite the complexity of the tumor microenvironment, Tex-NK1 divergence governs the primary axis of tumor-intrinsic and tumor-extrinsic variance. Distinct from the conventional view that NK cells positively contribute to immunotherapy efficacy, NK1-skewed tumors, although highly cytolytic, are refractory to current immune checkpoint blockade regimens. This killer divergence defines a foundational axis of cancer immunity and provides a resource for prioritizing next-generation targets for NK-directed immunotherapy.

## Introduction

Cancer immunotherapy, including immune checkpoint blockade (ICB) and adoptive cellular therapy, has transformed the treatment landscape of multiple malignancies, yet durable benefit remains confined to a narrow spectrum of cancer types and patients (1,2). The action of immunotherapy requires cytolytic killer lymphocytes—specifically CD8⁺ T cells and natural killer (NK) cells—to recognize and eliminate malignant cells (3,4). This concept underpins the prevailing “hot-cold” framework, which stratifies tumors along a spectrum from immune-excluded “cold” states to highly cytolytic “hot” phenotypes characterized by coordinated killer infiltration, enhanced interferon signaling, and checkpoint expression (5–13).

Although immune infiltration can positively predict immunotherapy response (5–12), a substantial proportion of patients with inflamed tumors still fail to benefit from current immunotherapies (14). We hypothesized the existence of a more fundamental, yet uncharacterized, layer of immune organization: heterogeneity in killer-cell dominance, which may shape therapeutic responsiveness and reveal new immunotherapeutic opportunities, particularly for tumors refractory to current immunotherapy strategies.

Recent studies have extensively delineated the nuanced subset diversity of killer cells at single-cell resolution (15–20). While invaluable, these cell-centric approaches have not yet led to a population-level stratification system and thus overlook a critical higher-order question: do killer cells exhibit coherent dominance patterns across individuals and cancer types? More specifically, which killer-cell states define the dominant axis of variation across tumors and malignancies, and how might this architecture inform immunotherapy resistance and therapeutic opportunity?

Therefore, distinct from previous cell-atlas descriptions, this study aimed to elucidate the population-level architecture of CD8+ T cells and NK cells based on large-scale pan-cancer scRNA-seq samples. We further validated these findings using spectral cytometry and spatial sequencing and linked them to single-cell immunotherapy efficacy data.

## Results

### Killer landscape in bulk sequencing

We first used bulk RNA-seq data from three large cohorts (TCGA, CPTAC, and UCSF-IPI, Table. S1) to comprehensively characterize the T/NK cells across cancer types. Twenty-five signatures encompassing infiltration, chemotaxis, differentiation, proliferation, activation, and inhibition were curated from the literature and MSigDB (Table. S2), and further filtered for non-overlapping genes. These signatures exhibited significant positive intercorrelations (Fig. 1A), consistent with the concerted T/NK actions in “hot” tumors reported in previous studies (5–13). When tumors were stratified into quartiles based on a global killer score derived from these signatures, the distribution proportions varied by cancer type; nevertheless, nearly all harbored a discernible subset of high-scoring (Q3/Q4) tumors (Fig. 1A), indicating that active immune surveillance can occur across most malignancies.

**Fig. 1.**
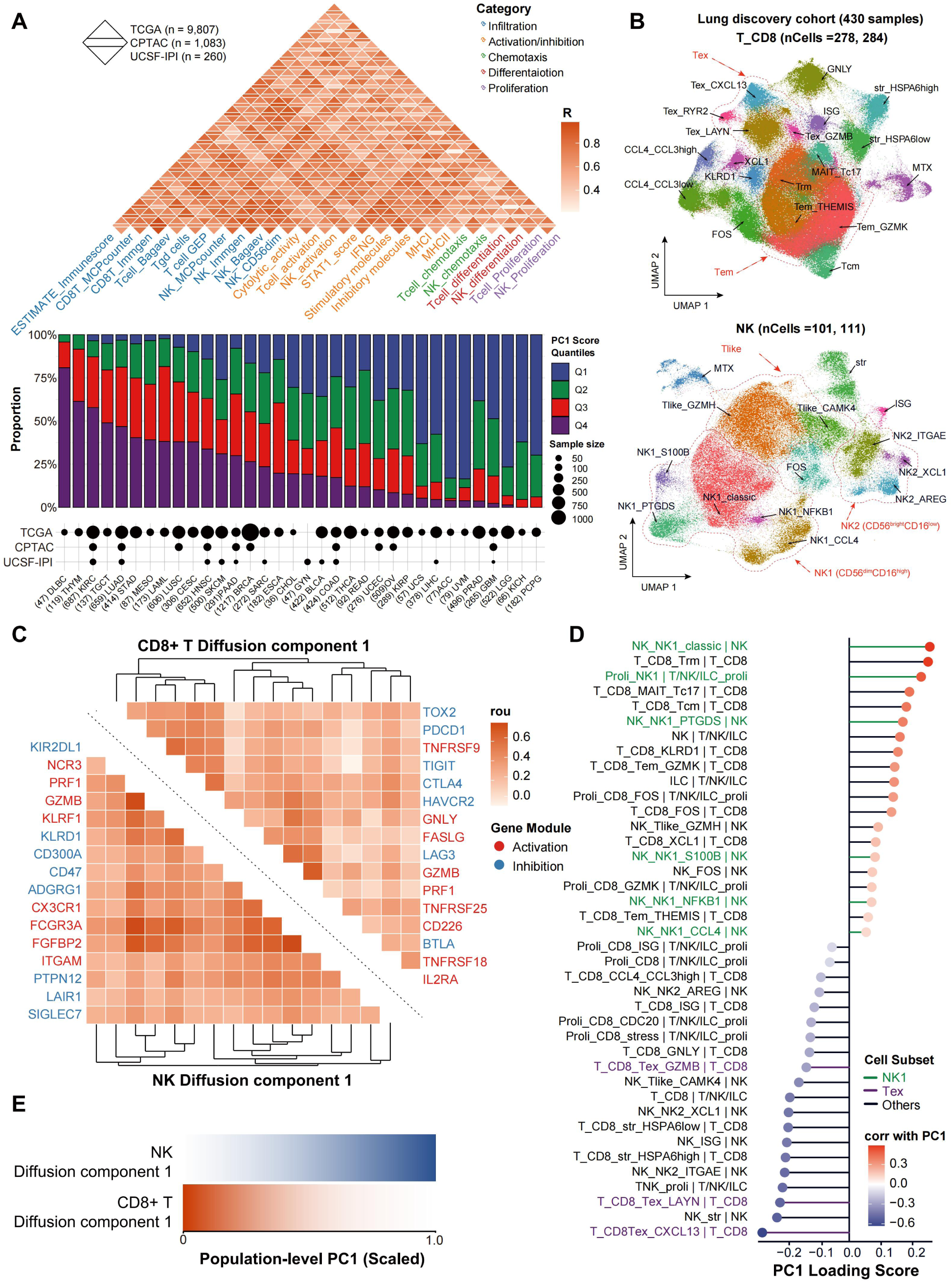
Divergence of cytotoxic lymphocytes across tumors. **(A)** (Top) Inter-correlation matrix of T and NK cell signatures evaluated across TCGA, CPTAC, and UCSF-IPI cohorts. All pairwise comparisons shown are significant. (Bottom) The proportion of patients stratified into quantiles based on a first principal component (PC1) compound score of the killer signatures across 31 cancer types. **(B)** Uniform manifold approximation and projection visualizations of single-cell profiles for CD8^+^ T cells (top) and NK cells (bottom) in lung discovery cohort. **(C)** The first diffusion component (DC1) comprised key activation and inhibition genes for both CD8^+^ T cells (right top) and NK cells (left bottom). **(D)** PC1 loading scores derived from the sample-level principle component analysis using proportions of CD8^+^ T and NK cell subsets. **(E)** Smoothed trend plot showing the fitted trajectory of the NK DC1 and CD8^+^ T DC1 gene module scores across the scaled PC1 continuum.

### Uncover killer dominance dichotomy in lung cancer

We are concerned that bulk sequencing may fail to differentiate between discrete T/NK states due to their substantial transcriptional similarity. Therefore, we sought to explore killer patterns at the single-cell level. Non-small-cell lung cancer (NSCLC) represents an ideal malignancy for initial investigation, as it exhibits a high proportion of “hot” tumors (Fig. 1A) yet manifests variable ICB responses across subtypes (21). By querying public databases and negotiating with investigators to access restricted data, we compiled single-cell RNA-seq (scRNA-seq) data comprising 430 NSCLC tumors from 343 patients, and meticulously curated detailed clinicopathological and genetic annotations (Fig. S1A and Table. S1). We only included treatment-naive samples to circumvent the confounding influence of prior treatment. This large cohort recapitulates the real-world epidemiology of NSCLC; for instance, EGFR mutations were more prevalent in females, Asian, and never-smokers (Fig. S1B).

We obtained 278,284 CD8^+^ T cells and 101,111 NK cells (Fig. 1B, Fig. S2, and Table. S3). Diffusion component analysis revealed that the primary axis of variation (DC1) in both CD8^+^ T and NK cells was driven by the co-expression of activation and inhibition signals (Fig. 1C and S3). Exhausted CD8^+^ T cells (Tex) and CD56^dim^CD16^hi^ NK cells (NK1) were enriched at the end of DC1, whereas memory/stress-associated CD8^+^ T cells and CD56^bright^CD16^low^ (NK2) cells congregated at the least pole (Fig. S3). Clustering of functional markers and signatures (Table. S1) consistently indicated that the primary heterogeneity of CD8^+^ T cells and NK cells subsets were governed by activation and inhibition features (Fig. S4). The two killer lineages also shared other phenotypic states characterized by high expression of stress signals (FOS, HSPA1A), cytokines (XCL1, CXCL4), metallothioneins (MT1X), or interferon-stimulated genes (STAT1, IFIT1) (Fig. S4). However, a key distinction was that tissue residence and interferon response programs were tightly coupled with DC1 terminal differentiation in CD8^+^ T cells but were decoupled in NK cells (Fig. S4). The NK_Tlike subset expressed T cell markers, but its global transcriptomic profile more closely resembled that of NK cells (Fig. 1B). We did not observe canonical NKT or memory/adaptive-like NK cell markers within this subset (Fig. S5). Clustering topologies remained consistent within the proliferating compartments (Fig. S6).

To define the primary sources of killer-cell variation across tumors, the proportions of above subsets in each sample were subjected to principal component analysis (PCA). Surprisingly, in stark contrast to the well-established parallel relationship between CD8^+^ T and NK cells (Fig. 1A), the first principal component (PC1) revealed a mutually antagonistic, “seesaw” relationship between the CD8^+^ T and NK cell immune surveillance programs (Fig. 1D). Cells with the strongest negative weights for PC1 comprised terminally activated Tex and the immature, regulatory NK2 cells. Conversely, the leading positive contributors to PC1 were terminally differentiated, cytotoxic NK1 cells and resident and central memory CD8^+^ T cells, which are generally considered bystander (22–24). Estimating the differentiation trend of CD8^+^ T and NK cells along the PC1 confirm they follow trajectories that are conversely intertwined with each other across samples (Fig. 1E). Consequently, tumors could be dichotomized into Tex-skewed (Ts) or NK1-skewed (Ns) based on the median Tex/NK1 ratio. Sensitivity analyses were performed across clinically relevant subgroups containing at least 15 samples, including age, sex, stage, anatomical site, smoking status, histological subtype, and mutation status, which demonstrated highly conserved killer divergence (Fig. 2A).

**Fig. 2.**
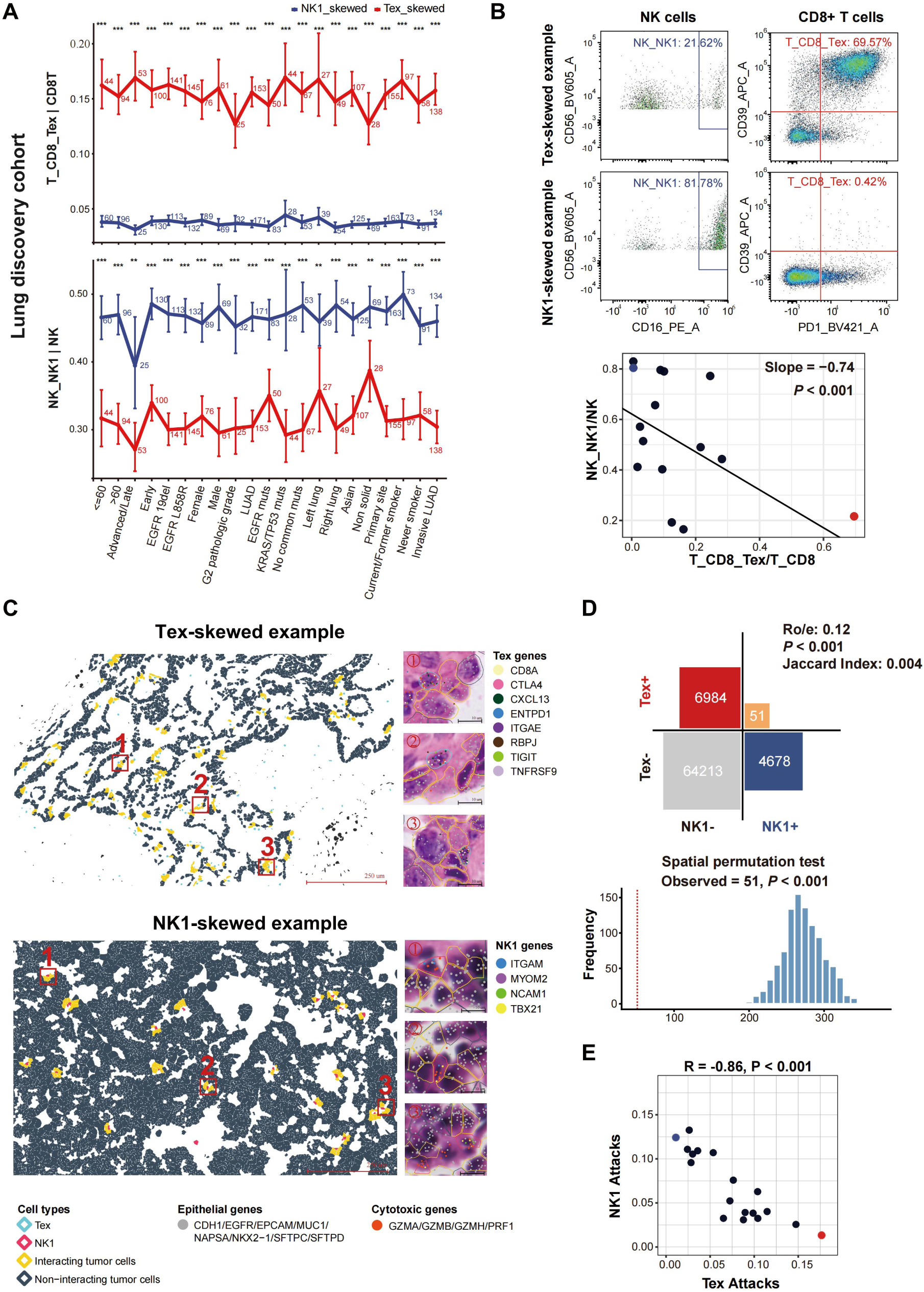
Robustness of Tex-NK1 dichotomy framework. **(A)** Comparison of the proportion of Tex and NK1 cells between Tex-skewed (Ts) and NK1-skewed (Ns) tumors across distinct clinicopathological and molecular subgroups within the lung discovery cohort. The specific sample size for each subgroup is indicated below each point. Statistical significance within each category was evaluated utilizing a Wilcoxon test and is denoted at the top (** *P* < 0.01; *** *P* < 0.001). The 95% confidence intervals are shown as vertical error bars, calculated via bias-corrected bootstrapping. **(B)** Spectral flow cytometry analysis on an independent cohort of 15 lung tumor patients. (Top) Representative gating plots identifying NK1 and Tex in prototypical Ts and Ns patients. (Bottom) Correlation between the proportion of NK1 and Tex cells. The solid black line represents the linear fit derived from a linear mixed-effects model incorporating acquisition date as random effects. The red and blue dots correspond to the two representative patient examples displayed in the top flow plots. **(C)** Xenium analysis mapping the distribution of Tex and NK1 cells in representative Ts (left) and Ns (right) samples. The bottom images (labeled 1-3) visualize individual transcripts at subcellular resolution overlaid on an H&E-stained background, highlighting putative cytotoxic interactions defined by a < 20 μm spatial proximity threshold. **(D)** (Top) Contingency matrix quantifying the spatial dichotomy of tumor cells engaged by Tex versus NK1 cells. (Bottom) Histogram of a spatial permutation test confirming that the observed frequency of dual-engaged tumor cells (*n* = 51, marked by the red dashed line) is significantly lower than the expected random distribution. **(E)** Correlation at the sample level between the frequency of tumor cells interacting with NK1 versus Tex.

Since Tex is a pivotal predictor of ICB response (25), revealing NK1 as a key effector in the opposing direction along the primary killer cell axis defines a distinct population that is likely refractory to current ICB or “ICB plus X” regimens, yet theoretically susceptible to NK-directed immunotherapies. Importantly, our observations do not contradict the established role of NK cells in activating CD8^+^ T cells. The PC1 axis demonstrates the same directionality for NK2 and Tex, and NK2 highly expresses the cytokines XCL1/XCL2 (Fig. 1B, S4 and S6), which have been shown to create an NK-DC niche for bridging innate and adaptive immunity (26).

We next assessed whether this phenomenon remained robust following justified adjustments to our analytical parameters. The Tex-NK1 axis consistently emerged as the primary axis of variation when using a non-imputation strategy prior to PCA (Fig. S7), or retaining clusters with high mitochondrial/ribosomal genes (Fig. S8). Furthermore, a significant negative correlation between Tex and NK1 was maintained in both non-proliferating and proliferating populations (Fig. S9A), when excluding the ambiguous NK_Tlike subset (Fig. S9B), when stratifying the analysis by sequencing platform or cell-sorting method (Fig. S9C), and when recalculating cell proportions using higher-order denominators (Fig. S10).

To orthogonally validate these findings, we performed spectral cytometry on a cohort of 15 patients with lung tumor (Fig. S11 and Table. S1). Again, lower Tex was significantly associated with higher NK1 (Fig. 2B and S12). Furthermore, the spatial relationship between Tex and NK1 was examined using Xenium spatial transcriptomics data from 18 NSCLC samples (Table. S1). We defined killer cells localized within close proximity (< 20 μm) of tumor cells as undergoing putative cytotoxic interactions (Fig. 2C). We observed a striking spatial dichotomy: tumor cells engaged by Tex cells were almost entirely excluded from interacting with NK1 cells, and vice versa, confirmed by R_o/e_ and permutation testing (Fig. 2D). Concordantly, at the sample level, the frequency of tumor cells interacting with Tex cells was strongly and inversely correlated with the frequency of those interacting with NK1 cells (Fig. 2E). This inverse correlation persisted when evaluating killer spatial density (Fig. S13). Altogether, the surveillance pattern of NK cells and CD8^+^ T cells diverge, rather than converge, representing a major source of variation in the anti-tumor immune response across tumors.

### Killer-cell dichotomy in pan-cancer

The reliability of this framework and whether it reflects a general hallmark of cancer are unknown. To this end, we collected 4,066 public and in-house tumor samples to build an extensive pan-cancer, treatment-naive scRNA-seq database (Fig. 3A and Table. S1). We integrated the advantages of reference- and marker-based approaches to develop an automated pipeline to efficiently annotate cells. Sample-level validation was performed after excluding the lung discovery cohort. A strong negative Tex-NK1 correlation was consistently observed across early- and late-stage tumors, encompassing most solid and hematological malignancies, as well as primary and metastatic lesions, and persisted across diverse technical subgroups stratified by sequencing method, cell-sorting strategy, and normalization method (Fig. S14). Notably, this divergence was also evident in precancerous lesions, indicating that the bifurcation of immune surveillance might initiate at the very onset of cancer development (Fig. S14).

**Fig. 3.**
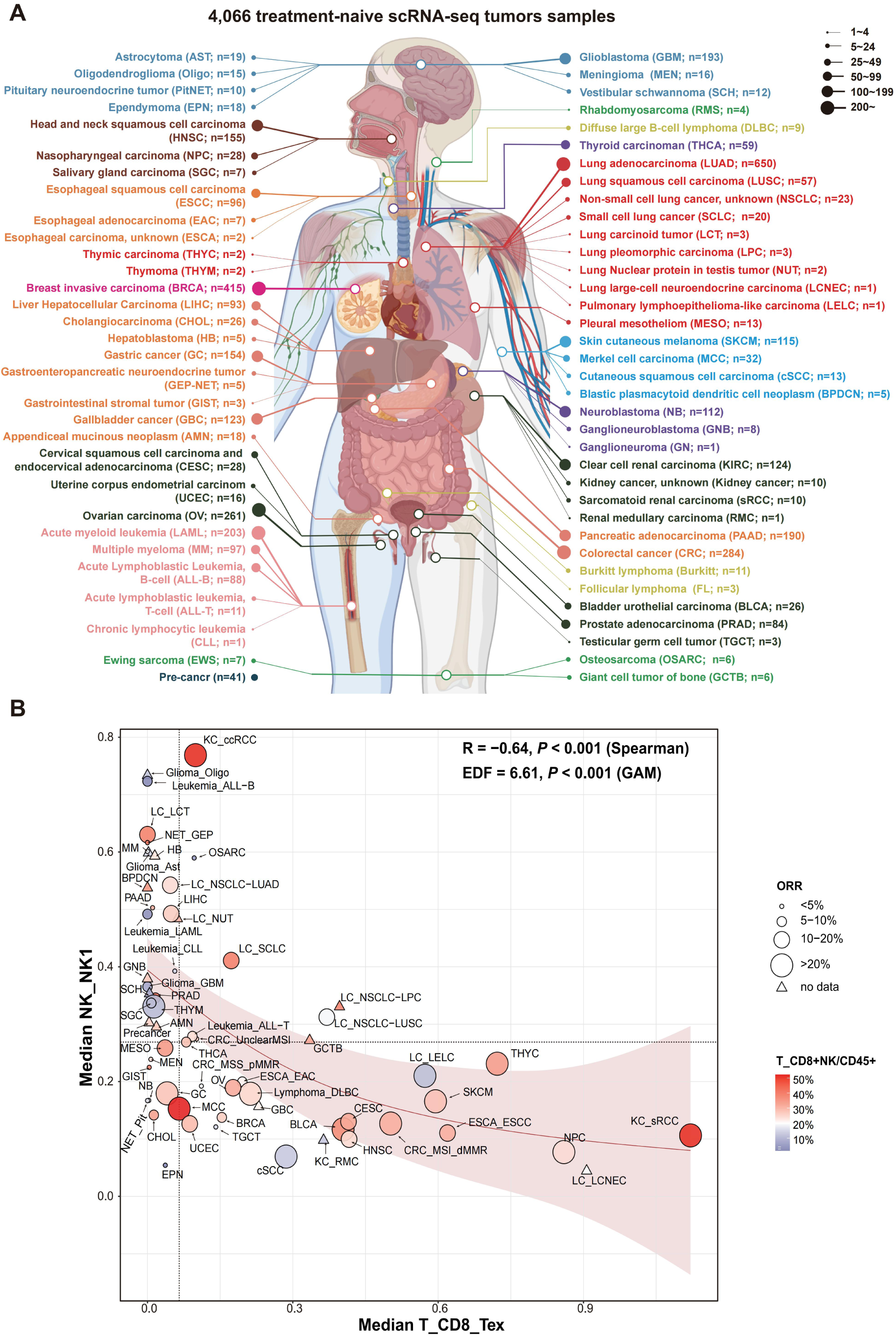
Pan-cancer landscape of killer-cell dichotomy. (**A**) Cancer distribution in the pan-cancer scRNA-seq database. (**B**) Each data point represents the median proportion of Tex versus NK1 cells for a specific malignancy. The inverse, non-linear trajectory of killer cell dominance is fitted utilizing a generalized additive model (solid red line), with the surrounding red-shaded region representing the confidence interval of the smooth term. Dashed vertical and horizontal lines denote the global median proportions across all evaluated cancer types.

At the cancer-type level, Tex and NK1 were highly mutually exclusive, such that very few malignancies showed simultaneously high levels of both killers when stratified by median values (Fig. 3B). To correlate this immune phenotyping with pan-cancer ICB efficacy, we pooled the objective response rate (ORR) data for ICB monotherapy for each cancer type (Table. S4). Cancer types skewed toward Tex were enriched for malignancies responsive better to ICB (e.g., High microsatellite instability colorectal cancer [MSI-H CRC], melanoma, esophageal squamous cell carcinoma, thymic carcinoma, and head and neck cancer [HNSC]). Conversely, Ns tumors primarily comprised indications that typically derive minimal benefit from ICB—such as pancreatic cancer, prostate cancer (PRAD), glioma, osteosarcoma, and acute myeloid leukemia (LAML). Exceptions existed, such as lung adenocarcinoma (LUAD) and clear cell renal carcinoma, where high ORRs might be driven by an elevated overall “hot” phenotype, reflected by high abundance of NK and CD8+ T cells. Importantly, the Tex-NK1 axis did not correlate with the conventional “hot-cold” status at cancer level (Fig. S15). The proportions of cancer types distributed across the four quadrants (defined by the hot/cold and Tex/NK1 axes) were comparable, indicating that these two axes represent complementary classification systems (Fig. S15). Proportion of ICB-beneficial cancer types was highest in the Tex-Hot quadrant (Fig. S16). Tex-Cold quadrant exhibited a higher proportion of ICB-beneficial cancer types than NK1-Hot quadrant, demonstrating that the Tex-NK1 divergence may be a more critical determinant of ICB efficacy than the hot-cold axis. The poorest outcomes were observed in the NK1-Cold quadrant (Fig. S16). Our framework can generate therapeutic hypotheses for rare but distinctly skewed malignancies in which immunotherapy trials have not yet been conducted, such as hepatoblastoma (Ns) and lung large-cell neuroendocrine carcinoma (Ts) (Fig. 3B).

We next tested whether killer divergence could be further sculpted by clinical factors even within a malignancy (Fig. S17 and S18). LUAD at advanced/late stages preferentially accumulated Tex compared to early-stage disease, whereas HNSC and gallbladder cancer were progressively enriched for NK1. However, these variations largely belonged to the same Ts or Ns category on a broader pan-cancer scale, and stage had no significant effect in most cancer types. Similarly, stratification by anatomic site revealed that variations within most tumors were category-restricted. Nevertheless, compared with primary tumors, liver metastases exhibited higher NK1 in breast cancer (BRCA) and gastric cancer (GC). Brain metastases drove distinctly high Tex infiltration in melanoma but had no effect in LUAD. Cavity metastases, such as pleural effusions in LUAD and ascites in GC and ovarian cancer (OV), tended to be “cold”, exhibiting lower levels of both Tex and NK1.

Profiling molecular subtypes within cancers (Fig. S19) revealed that, early-stage LUAD was predominantly NK1-driven regardless of mutation type, particularly in tumors harboring EGFR_L858R, SMARCA4, ATM, or ARID1A/B mutations. Conversely, advanced-stage LUAD exhibited a more dispersed landscape spanning the Ns to Ts spectrum. EGFR mutations favored NK1, whereas tumors lacking common driver mutations and those with TP53/KRAS co-mutations favored Tex. STK11- and/or KEAP1-mutant tumors exhibited low levels of both Tex and NK1. This observation mirrors the clinical efficacy of ICB in advanced LUAD, which is effective in TP53/KRAS-mutant or driver-negative tumors, but markedly less effective in EGFR-or STK11/KEAP1-mutant settings (21). Notably, IDH-WT glioblastoma and virus-negative hepatocellular carcinoma exhibited stronger NK1 dominance than their respective overall populations. Evaluation of BRCA and OV molecular subtypes revealed limited variation.

When using a multivariate linear model to predict Tex or NK1 levels, cancer type accounted for the highest proportion of the explained variance (20∼30%), whereas stage, anatomic site, and molecular subgroup contributed minimally (1∼2%). Notably, approximately 70% of the total variance remained unexplained, suggesting that killer divergence are shaped by more complex, unmeasured factors (Fig. S20).

### Killer-cell dichotomy dominates global tumor-immune variation

Although many aspects of intratumor heterogeneity (ITH) have been reported, not all are equally important. Isolated investigations lacking a broader perspective can lead to a suboptimal understanding of the major axes of tumor-intrinsic and tumor microenvironment (TME) variation, which are critical for identifying patient populations where precision medicine can deliver the greatest value.

We first investigated whether killer-cell divergence reflects immune surveillance over distinct tumor cell hallmarks. The “missing-self” theory prompted us to test whether tumors with low epithelial MHC-I scores might be enriched for NK1 rather than Tex to facilitate tumor eradication. We did observe a positive correlation between Tex and MHC-I in a limited number of cancer types (LUAD, BRCA, OV, MSI-H CRC, and thyroid carcinoma), as well as a negative correlation between NK1 and MHC-I in others (PRAD, BRCA, and HNSC); however, this pattern was not observed for other majority of cancer types (Fig. S21).

To gain an unbiased view of the global malignant network, we detected functionally non-redundant co-expression meta-programs (MPs) of tumor cells that recurrently occur across tumors in the lung discovery cohort (Fig. 4A, S22, and Table. S5). Most MPs could be mapped to a pan-cancer MP resource (27), including Cell cycle, Respiration, Stress, and lung tissue-related (Cilia and Alveolar) (Table. S5). However, an MP enriched for ribosomal protein genes, which emerged as the most recurrent MP in our cohort, was excluded from the previous resource (27) due to suspicions of reflecting low-quality data. Nevertheless, we believe this MP conveys biologically meaningful information because it could represent highly active protein synthesis and other studies have retained this as a functional MP (28). We thus preserved this MP and designated it as “Translation”. Additionally, we identified a novel MP, ranking second in overall recurrence, whose top enriched pathways were associated with responses to trabectedin and ultraviolet irradiation (Table. S5), both of which trigger the DNA damage response (DDR). This MP may indicate a adaptive upregulation of core networks to survive to genotoxic stress. It also includes stress-responsive lncRNAs (MALAT1, NEAT1, XIST) and DNA repair factors (SPIDR, CUX1). Distinctly, another Stress MP is highly concordant with previous definition (27), driven by AP-1 transcription factors (FOS, FOSB, JUN), anti-proliferative genes (CDKN1A, GADD45B, BTG2), anti-apoptotic factors (MCL1), and heat shock chaperones (HSPA1A, HSPA1B). Therefore, we designated these two MPs as the “Stress-DDR” and “Stress-classic”, respectively. Clustering of MPs across tumors revealed two distinct MP families: 1) one comprising the Cell cycle, Translation, Respiration, and Cilia MPs, and 2) another grouping the two Stress MPs with the Alveolar MP (Fig. S23).

**Fig. 4.**
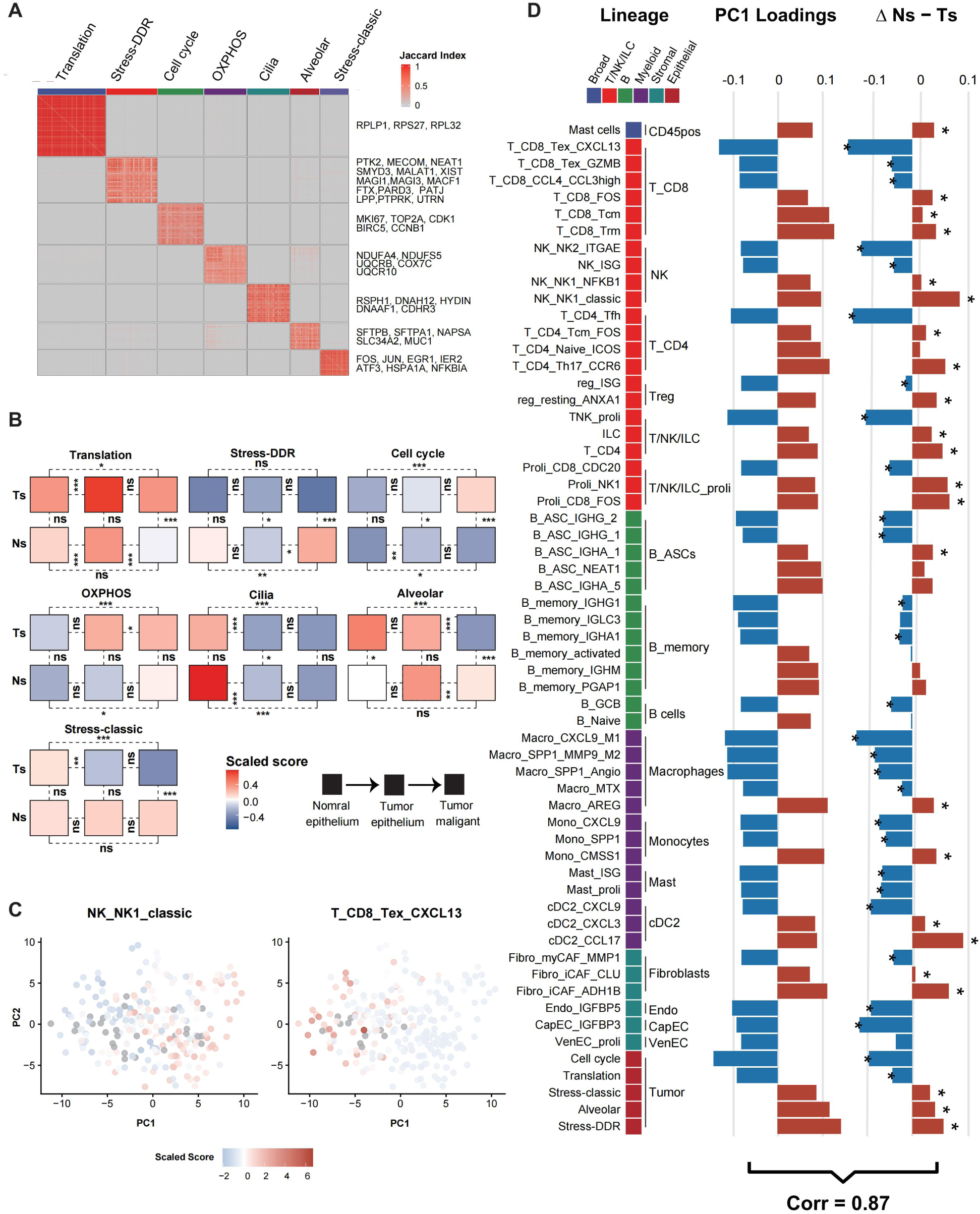
Killer-cell dichotomy orchestrates global tumor-intrinsic programs and the tumor microenvironment (TME). (**A**) Jaccard similarity index matrix delineating seven functionally non-redundant, recurrent co-expression meta-programs (MPs) across malignant cells in the lung discovery cohort. (**B**) Median scaled activity scores of each MP across a continuum of normal epithelium, intratumoral non-malignant epithelium, and malignant tumor cells. Patient samples are stratified into Tex-skewed (Ts) and NK1-skewed (Ns) groups. Statistical significance was assessed utilizing the Wilcoxon rank-sum test for cross-sectional Ts versus Ns comparisons, and linear mixed-effects models, with patient identity included as a random effect, to evaluate epithelial progression (* *P* < 0.05; ** *P* < 0.01; *** *P* < 0.001; ns, not significant). (**C**) Two-dimensional PC1 and PC2 plots visualizing the infiltration levels of NK1 (top) and Tex (bottom), derived from the principal component analysis of 298 multi-lineage TME cellular composition metrics across tumors. (**D**) Paired horizontal bar plots linking the primary axis of global TME variance to the Tex-NK1 divergence. (Left) PC1 loading scores for the top ±30 contributing multi-lineage TME features. (Right) The corresponding differential abundance of these specific subsets between Ns and Ts tumors. Individual bars are color-coded by their respective overarching cellular lineages. The significance of differential abundance between groups is denoted by asterisks (*FDR < 0.05).

Both the number of detected MPs and copy number variation clone were comparable between the Ts and Ns groups (Fig. S24). However, they differed drastically in their MP profiles. Ts tumors exhibited higher activity in MP family 1 (Cell cycle and Translation), whereas Ns tumors were enriched for MP family 2 (Fig. 4B). The MP differences were absent when comparing matched adjacent normal tissue epithelium or intratumoral non-malignant epithelial cells, indicating that the Ts and Ns tumor states diverge during tumor evolution (Fig. 4B). We found that Ts tumors possessed a higher proportion of undifferentiated tumor cells than Ns tumors based on CytoTrace2 (29) (Fig. S25). These observations remained consistent when restricting the analysis to tumor cells defined by the overlap of inferCNV (30) and CopyKat (31) predictions (Fig. S26). Although tumors eventually undergo immune escape, these findings suggested that CD8^+^ T cells preferentially survey proliferating, translationally active tumor states, which may ensure a continuous supply of synthesized proteins for rapid antigen presentation (32). In contrast, NK cells are more likely to govern tumors characterized by quiescence, stress-induced growth arrest, and injury-repair mimicry, which might be driven by the induction of NK-activating ligands (33). Intriguingly, the Stress-DDR MP also showed a weak but significant enrichment for previously described Glioma and Pancreatic MPs (27) (Table. S5), two malignancies highly skewed toward NK1 (Fig. 3).

To further explore whether killer divergence reflects broader TME complexity, we performed deep clustering across all TME cell lineages and obtained 298 cellular composition metrics for each case in the lung discovery cohort (Fig. S2, Tables S3 and S6). Strikingly, PCA based on these metrics and the tumor MP scores showed that NK1 and Tex remained the primary drivers of PC1 variation (Fig. 4C). Many of the other top-loading features along PC1 were significantly differentially abundant between the Ts and Ns groups, as demonstrated by a strong correlation (Fig. 4D). Cell subsets enriched in Ts tumors included key components of tertiary lymphoid structures (34) and T-myeloid networks (35), such as CD4^+^ T follicular helper (Tfh) cells, IgG^+^ plasma cells, and CXCL9-high subsets of macrophages and monocytes. The coupling of proliferating T/NK cells and tumor MPs in Ts tumors was recently noted in a spatial transcriptomics study of breast cancer, which found that their interaction is a key factor in ICB response (36). Of note, Ts tumors also exhibited higher Macro_SPP1 and Mono_SPP1 infiltration. While this appears contradictory to a previous study demonstrating the CXCL9/SPP1 axis as an important TME feature (37), this discrepancy might arise from a lack of unbiased sample-level analysis in prior work, or because this axis represents a sub-branch within the broader Tex-skewed direction. Regarding stromal components, Ts tumors accumulated myofibroblasts and various endothelial cell subsets. Ns tumors were characterized by components such as naive CD4+ T cells, innate lymphoid cells, resting regulatory T cells, Macro_AREG, several subsets of type 2 conventional dendritic cells (cDC2), IgA+ plasma cells, inflammatory fibroblasts, and stress-related tumor MPs. This network was largely undescribed until a recent study (38) elucidated that the cDC2-CD4⁺ T cell niche was indispensable for NK-dependent tumor control. Some of these cell subsets are further discussed in the following section on target prioritization. Repeated PCA using lower-granularity cell clustering yielded consistent trends (Fig. S27 and Table. S6). Altogether, the Tex-NK1 axis dominates the cellular variability of the TME ecosystem, standing out as the primary branch upon which immense complexity converges.

### High NK1 defines population refractory to T cell-based immunotherapy

Cytotoxic NK1 cells are generally considered a positive predictor of ICB response^5^. A study published last year also support that higher NK1 positively associated with anti-PD1 response in NSCLC (39). However, because this study utilized exclusively post-treatment samples, the findings might reflect an immunological state conflated by the treatment. Our results presented above suggest that tumors enriched for NK1 represent an evolutionarily independent population that does not benefit from current T-cell-based immunotherapies. To explicitly demonstrate this point, we compiled a comprehensive database comprising 628 scRNA-seq samples and 7,346 bulk RNA-seq samples with available ICB efficacy data from both in-house and public cohorts (Fig. 5A and Table. S1). All bulk samples were collected prior to ICB treatment, whereas the scRNA-seq cohort included site-matched post-treatment samples to evaluate dynamic shifts.

**Fig. 5.**
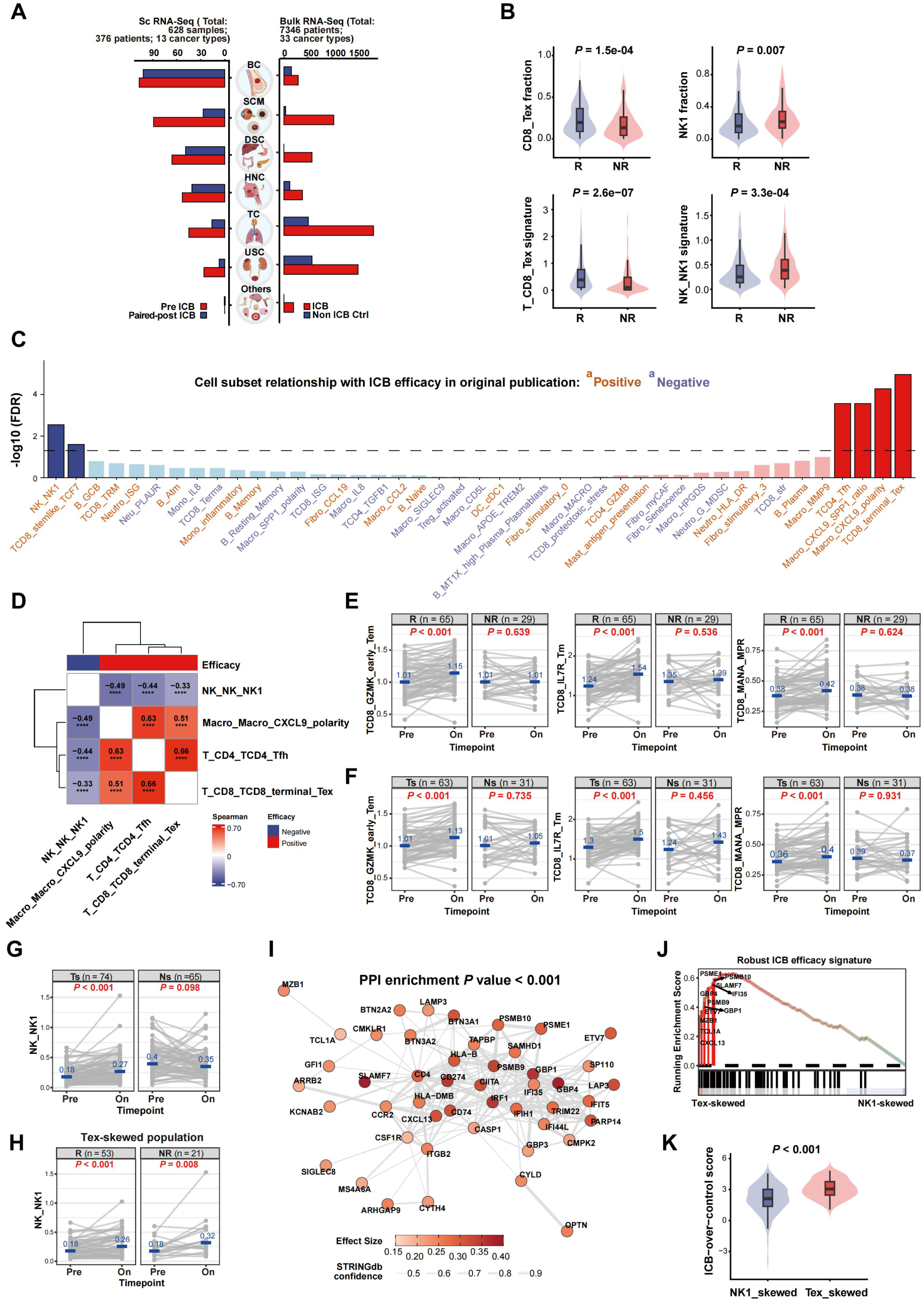
Killer-cell dichotomy dictates sensitivity to immune checkpoint blockade. (**A**) Schematic summarizing the large-scale immunotherapy transcriptomic database. BC, breast cancer; SCM, skin cancers and melanoma; DSC, digestive system cancers; HNC, head and neck cancers; TC, thoracic cancers; USC: urinary system cancers. (**B**) Pre-treatment fractions and continuous signature scores of Tex (left) and NK1 (right) compared between ICB responders (R) and non-responders (NR) in the scRNA-seq cohort. Internal box plots display the medians and interquartile ranges. (**C**) Benchmarking the predictive performance of curated single-cell subsets for ICB response. Subsets are color-coded by their relationship with ICB efficacy in the original publications (red, positive; blue, negative). The horizontal dashed line denote a FDR < 0.05. (**D**) Pairwise Spearman’s rank correlations among the ICB-responsive cell subsets and NK1 (* *P* < 0.05; *** *P* < 0.001; **** *P* < 0.0001). (**E**) Dynamic changes in memory CD8⁺ T cell signatures between matched pre-treatment and on-treatment samples. Data are stratified by clinical response, with *P* values reflecting paired Wilcoxon signed-rank tests. (**F**) Dynamic changes in the identical CD8⁺ T cell signatures from (E), but re-stratified according to the patients’ baseline killer profiles. (**G**) Dynamic shifts of NK1 cells compared between baseline Tex-skewed (Ts) and NK1-skewed (Ns) groups (Left), (**H**) and further stratified by clinical response specifically within the Ts population (Right). (**I**) Association of pseudobulk transcriptomic profiles with a robust ICB efficacy signature derived from bulk RNA-seq cohorts. STRING protein-protein interaction network of 60 positive ICB efficacy genes. Node color intensity and size reflect the interaction effect size. **(J)** Gene set enrichment analysis running enrichment score plot evaluating the ICB efficacy signature between the Ts and Ns populations. (**K**) Comparison of the ICB-over-control score between Ts and Ns groups. Internal box plots delineate the median.

The mutually exclusive correlation between Tex and NK1 was recapitulated in the scRNA-seq cohort (Fig. S28). Pooled analysis of pre-treatment samples revealed that ICB responders had significantly higher Tex and lower NK1 cell proportions compared to non-responders, a finding that remained consistent when evaluating continuous signature scores (Fig. 5B). To avoid confounding effects from combined treatments, we restricted our analysis to samples treated with ICB monotherapy and observed consistent findings (Fig. S29).

To benchmark the predictive performance of the Tex-NK1 axis against cell subsets reported as biomarkers in single-cell studies (Table. S2), we evaluated the association of each biomarker signature with ICB response in the scRNA-seq cohort (Fig. 5C). Surprisingly, most published signatures yielded non-predictive or even contradictory results compared to the directionality reported in the original studies. Tex and NK1 emerged as the top-ranking positive and negative predictors, respectively (Fig. 5C). Tfh and Macro_CXCL9 also remained significant after adjusting for the false discovery rate (FDR). The predictiveness of the proposed CXCL9/SPP1 ratio (37) in macrophages appears to be largely driven by the CXCL9 subset. TCF7^+^ CD8^+^ T cells exhibited a negative predictive value, aligning with evidence that TCF7 expression is higher in non-tumor-reactive bystander clonotypes (22–24). ICB-responsive cells passing FDR correction clustered into a hub, from which NK1 cells were distinctly excluded (Fig. 5D), consistent with the PC1 axis above (Fig. 4D).

Mechanistically, the action of ICB is driven by the targeting of inhibitory checkpoints and the expansion of progenitor memory CD8^+^ T cells through reinvigoration or clonal replacement^3^. Thus, we reasoned that these features could serve as biological endpoints to further evaluate differential ICB sensitivity between Ts and Ns tumors. The scRNA-seq and cytometry jointly suggested that NK1 cells exhibited a distinct checkpoint spectrum and expressed low levels of conventional T cell-related checkpoints such as PD-1 (Fig. S30). To systematically assess T cell dynamics following ICB, we curated several relevant signatures (Table. S2) and analyzed site-matched pre- and on-treatment tumor samples to determine whether their shifts correlated with clinical response and baseline killer profiles. Among these, GZMK^+^ early Tem, IL7R^+^ memory cells, and an experimentally validated signature associated with a major pathologic response in post-treatment mutation-associated neoantigen (MANA)-specific CD8^+^ T cells (40) were significantly increased in on-treatment samples from responders, but not from non-responders (Fig. 5E and Fig. S31A). Crucially, these post-treatment expansions occurred exclusively in patients with a baseline Ts profile and were absent in the Ns group (Fig. 5F and Fig. S31B). This highlights that baseline Ts tumors phenocopy the responsive CD8^+^ T cell dynamics necessary for ICB efficacy. Because cytotoxic or clonal revival can occur at early time points, such as 2 to 4 weeks (3), we next assessed whether the time gap between sampling affected our findings. By classifying samples into early-on and late-on treatment groups using a 4-week cutoff, we observed that the dynamic expansion of the three significant CD8^+^ phenotypes was retained in both early- and late-on responder samples, and remained restricted to the Ts group (Fig. S32 and S33). These results highlight an early immune mobilization that is limited to the Ts population.

A recent study (41) suggests that NK1 cells increase in non-responders after treatment with ICB. Interestingly, however, we found that NK1 cells increased in the Ts group, whereas they remained unchanged in the Ns group (Fig. 5G). This expansion in the Ts group occurred regardless of response, which may reflect a byproduct effect irrelevant to ICB efficacy.

To assess whether the Tex-NK1 divergence underlies variable responses to other T cell-directed immunotherapies, we evaluated 18 pre-treatment samples from melanoma and glioblastoma patients treated with adoptive T cell therapy (Table. S1). We consistently observed a negative correlation between Tex and NK1, with clinical responders being enriched within the Ts population, although statistical significance was not reached, likely due to the limited sample size (Fig. S34).

scRNAseq is often limited by small sample sizes and a lack of long-term survival data. To address this, we assessed whether the pseudobulk transcriptomic difference between the killer-skewed groups were associated with ICB efficacy signatures derived from massive bulk RNA-seq ICB datasets (Fig. S35 and Table. S1). We performed a meta-analysis across more than 100 cohorts to define a robust set of genes that consistently and significantly predict ICB efficacy across three clinical outcome categories: response, disease control endpoints such as progression-free survival, and overall survival. We restricted these ICB efficacy-related genes to those exhibiting high gene-wise concordance between the pseudobulk profiles of our lung discovery cohort and the actual bulk expression levels in the TCGA-NSCLC cohort (Fig. S35). This filtering yielded 60 valid positive genes, whereas no negative genes passed these thresholds. These efficacy-associated genes were significantly enriched for protein–protein interactions and mapped primarily to active immune responses (Fig. 5H and Table. S7). Gene set enrichment analysis showed that this signature was significantly more enriched in the Ts group compared to the Ns group, indicating a pan-transcriptome shift toward ICB benefit in Ts patients (Fig. 5H). Repeating our analysis in ICB monotherapy studies showed consistent findings (Fig. S36 and Table. S7).

Furthermore, through a meta-analysis of treatment-by-gene interaction tests across eight randomized controlled trials, including two in-house trials, ORIENT-11 (42) and RATIONALE-309 (43) (Fig. S35 and Table. S1), we derived a score that predicts the benefit of ICB over conventional therapies across three types of endpoints. This metric was also significantly higher in the Ts group than in the Ns group (Fig. 5I).

### Prioritize targets for NK1

Given that the Ns population occupies the opposite extreme of the ICB sensitivity spectrum, this framework provides a unique evolutionary lens for prioritizing targets for next-generation immunotherapies aimed at activating NK1 (Fig. 6A). We first searched for receptors upregulated on NK1 cells and concurrently upregulated ligands within the TME of the Ns population compared with the Ts population in the lung discovery cohort. Interactions with NK1 were primarily driven by malignant and myeloid cells (Fig. 6B). We identified CD47 as a top-ranking candidate, whose ligand, thrombospondin-1 (THBS1), has been shown to suppress NK function (44) and exhibited the strongest signal here. CD47 expression is also markedly higher in NK1 than in Tex (Fig. S30). The identified CX3CR1-CX3CL1, CXCR2-CXCL2/3, and ITGAM-ICAM1 interactions may indicate active NK cell recruitment and immune synapse formation (45,46). Importantly, aligned with the NK1-related TME network, multiple ligands correlate with Alveolar/Stress tumor MP scores, and high expression levels of THBS1, CXCL2, and CXCL3 are detected in Macro_AREG and cDC2_CXCL3, which are critical PC1-positive features (Fig. S38 and Fig. 4D).

**Fig. 6.**
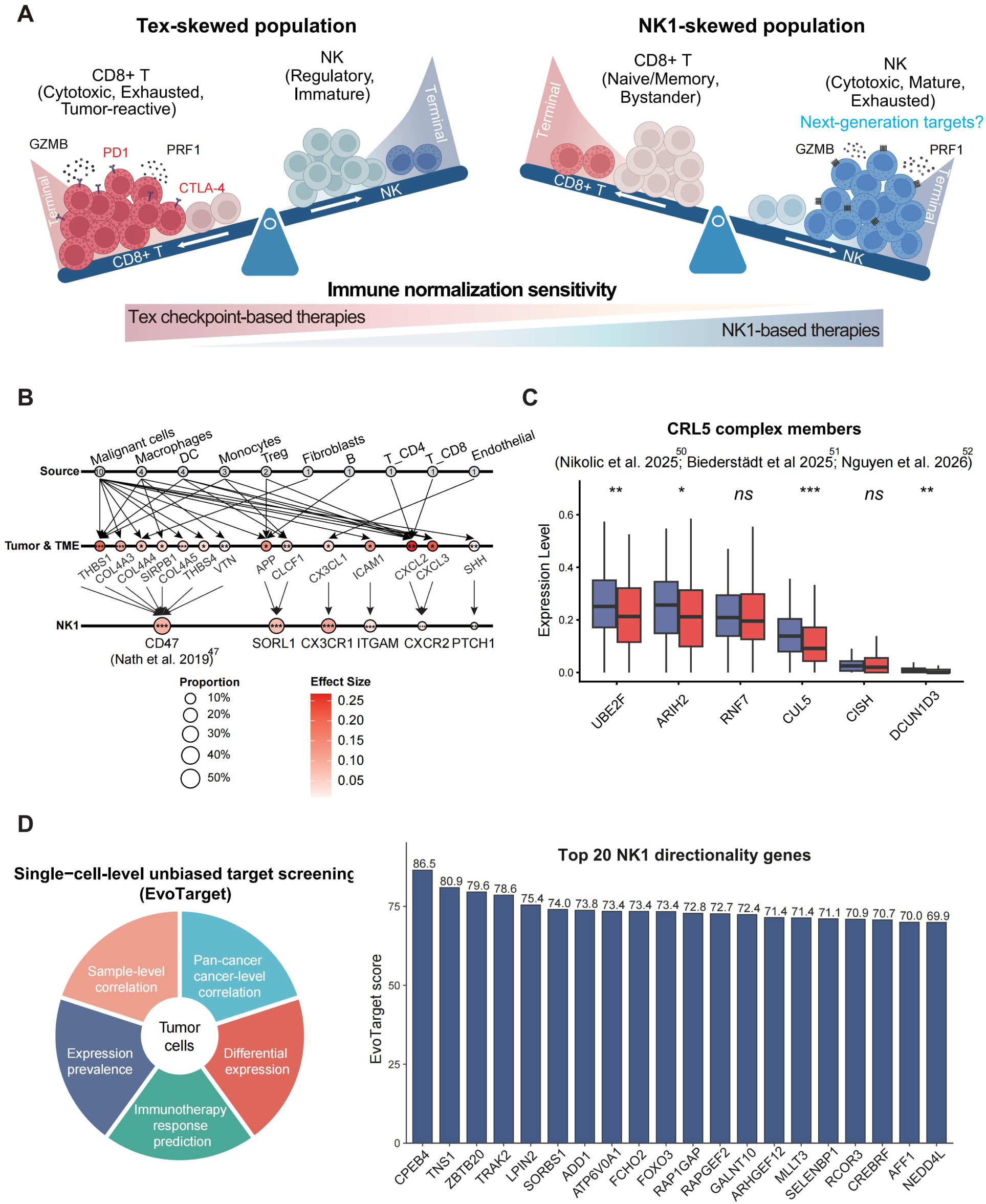
Prioritize targets in NK1-skewed tumors. (**A**) Conceptual seesaw model illustrating the mutually antagonistic balance between CD8⁺ T cell and NK cell surveillance programs. (**B**) Ligand-receptor interaction network displaying the significant candidates. The top layer displays the specific sender cell lineages, the middle layer lists the ligands, and the bottom layer delineates the corresponding receptors on NK1 cells. Asterisks inside the nodes denote the statistical significance of differential expression between the Ns and Ts populations. (**C**) Expression of experimentally validated Cullin-5 RING E3 ligase complex (CRL5) members within the NK1 cell subsets between the two groups. (**D**) EvoTarget screening evidence layer diagram (left) and the top 20 NK1-directional genes (right). * *P* < 0.05; ** *P* < 0.01; *** *P* < 0.001; ns, not significant.

Irrespective of the experimental setup, three independent groups performed genome-wide CRISPR screens in primary human natural killer cells and identified the Cullin-5 RING E3 ligase (CRL5) complex as a key intracellular negative regulator, the blockade of which enhances IL-2/IL-15 signaling and killer cytotoxicity (47–49). Upon projecting their findings onto our dataset, we found several CRL5 genes (UBE2F, ARIH2, CUL5, DCUN1D3) were significantly upregulated in NK1 cells of the Ns group compared to the Ts group (Fig. 6C). Although the expression of IL-2 and IL-15 cytokines did not differ between the killer groups (Fig. S39A), we observed that NK1 cells within Ns tumors exhibited reduced cytokine activity based on CytoSig (50) (Fig. S39B).

To more unbiasedly identify tumor-cell factors that may activate or inhibit NK1 cells in the Ns population beyond known mechanisms, we developed a screening strategy termed EvoTarget to prioritize tumor-cell genes skewed toward the NK1 axis by integrating five independent single-cell-derived evidence layers (Fig. 6D): 1) high tumor-cell expression prevalence in Ns; 2) tumor-cell differential expression in Ns over Ts; 3) cross-cohort/subgroup sample-level positive and negative correlations with NK1 and Tex states, respectively; 4) concordant positive and negative cancer-level correlations with NK1 and Tex across tumor types; and 5) negative prediction of ICB response. Most of the top NK1-directional genes remain unexplored in the context of NK1 function (Fig. 6D and Table S8), providing a rich resource for future mechanistic studies.

## Discussion

Here, we put forward a highly generalizable framework complementary to the established “hot-cold” paradigm, demonstrating that Tex and NK1 represent a dominant axis of ITH across the pan-cancer landscape. Our findings provide a rationale for why current ICB regimens targeting Tex have been the most successful strategy in recent decades, while simultaneously establishing that targeting NK1 represents an independent, promising therapeutic avenue.

The fact that blocking PD-1 alone is sufficient to profoundly reset the immune response in a subset of patients indicates that knowing which immune defect acts as the dominant suppressive master switch for tumor escape is critical (51). This success has driven extensive efforts aiming to overcome immunotherapy resistance by targeting alternative checkpoints, yet they typically yield only modest gains (52). Determining which factor is most important among all immune defects identified within the TME remains largely empirical due to the lack of established population-level, TME-wide systematic evolutionary patterns. The mutual exclusivity of the overall TME and tumor-intrinsic programs between the Ts and Ns populations, combined with their distinct checkpoint spectra, suggests that the three phases of cancer immunoediting (elimination, equilibrium, and escape) manifest via distinct mechanisms within these respective groups. Within the Ns population, Darwinian selection drives tumors to develop evasion mechanisms primarily to escape the pressure exerted by NK cells rather than CD8⁺ T cells. Unlike T cells, NK cells lack well-characterized checkpoints. The conserved Tex-NK1 axis here would aid in finding a master “brake” driving NK dysfunction.

NK immunotherapy has garnered rapidly growing interest in recent years, driven by its broad antigen/MHC-independent recognition capacity, constitutive expression of cytotoxic granules for rapid responses, high tolerance toward normal tissues, and favorable safety profile (5,45). As of manuscript submission, more than 1,600 clinical trials evaluating NK-based therapies in cancer are registered on ClinicalTrials.gov, encompassing checkpoint inhibitors, cytokine agonists, chimeric antigen receptor (CAR)-NK, and NK cell engager. Although most NK trial data are not yet available, anecdotal evidence connects our landscape findings with emerging clinical observations. For instance, CAR-NK therapies have recently demonstrated encouraging efficacy and a manageable safety profile in difficult-to-treat solid tumor glioblastoma (NCT06061809), where the median overall survival has not yet been reached. Consistently, this malignancy was identified as distinctly NK1-skewed based on our pan-cancer Tex-NK1 mapping. Implementing killer profile enrichment will increase the likelihood of detecting robust clinical signals while minimizing immune-related adverse events.

Other findings indicate that there may be broad, inherently susceptible patient populations for deploying NK-based therapies. We observed an increase in NK1 among patients with baseline Ts tumors following ICB regardless of clinical response, suggesting that targeting this expanded compartment may be well-suited for sequential treatment strategies. The killer cell divergence rule also applies to precancerous tissues, which are predominantly skewed toward NK1, highlighting the potential of such interventions at the earliest stages of tumorigenesis for effective disease interception.

A major limitation of this study is that its findings are primarily correlative rather than causal. The mechanisms driving the formation of killer-cell divergence, as well as the key dysfunctional molecules underlying each state, remain unknown. However, these questions are beyond the scope of the current landscape study. What we provide here is a robust framework for future killer-profile-specific patient stratification and target prioritization, moving beyond the conventional goal of simply converting cold tumors into hot tumors. Additionally, although our proposed killer divergence framework significantly reduces the dimensionality of profound ITH, we acknowledge that it does not fully resolve all sources of complexity. Nevertheless, we believe that ITH is not inherently fragmented, but rather operates within a systematic “branch-and-leaf” framework. For example, we observed that well-defined Tex-driving cell subtypes, such as SPP1^+^ macrophages (53) and myofibroblasts (54), along the same PC1 direction with Tex. This suggests that patients skewed toward the same branch direction may harbor distinct, context-specific exhaustion mechanisms. Future studies should further extend our framework to map the “leaves” of TME ITH by elucidating the roles of coexisting tumor cells and TME contexts across different patients, ultimately guiding tailored combination regimens to more thoroughly reverse killer exhaustion. This is particularly crucial for the Tex population, where the main “branch” treatment is already established.

Immunotherapy remains the most promising strategy for achieving a cure for cancer. The Tex-NK1 dichotomy as a foundational paradigm will extend the clinical promise of immunotherapy to a broader demographic, shifting the field from an exclusively T-centric focus toward entirely new therapeutic directions. The Ns population must be considered a key subgroup when exploring novel immune normalization targets and designing clinical trials.

## Methods

### Spectral cytometry data generation

Fresh lung tumor surgical specimens (Table. S1) were placed in MACS Tissue Storage Solution (Miltenyi Biotec) on ice for preservation and transport until the preparation of single-cell suspensions. The samples were dissociated into single-cell suspensions using the Tumor Dissociation Kit human (Miltenyi Biotec, 130-095-929) at 37°C with 100 rpm agitation for 40 min, with the enzyme cocktail proportionally adjusted based on the weight of each individual sample according to the manufacturer’s protocol. To optimally preserve surface epitopes for T and NK cell analysis, the protocol was modified by reducing Enzyme R to 20% of the standard recommendation (10 µL for <0.2 g tissue; 20 µL for 0.2-1.0 g tissue). Following digestion, suspensions were passed through a 70-µm mesh. Erythrocytes were lysed with 1X RBC Lysis Buffer (BioLegend, 420301) for 5 min at room temperature. Cells were subsequently stained for viability using the Zombie NIR™ Fixable Viability Kit (1:200 dilution; BioLegend, 423105) for 15 min at room temperature in the dark, and the reaction was stopped with 1 mL of Cell Staining Buffer. To block non-specific binding, cells were incubated with Human TruStain FcX™ (5 µL per 10⁶ cells; BioLegend, 422301) for 5 min at room temperature. Without washing, cells were immediately stained with a surface antibody cocktail for 30 min at 4°C in the dark. The panel designed to detect exhausted CD8⁺ T cells (Tex) and CD56^dim^CD16^hi^ NK cells (NK1) consisted of the following BioLegend antibodies: Alexa Fluor® 488 anti-human CD45 (304017), Spark Red™ 718 anti-human CD3 (300340), Brilliant Violet 785™ anti-human CD8 (344739), Brilliant Violet 421™ anti-human PD-1 (367422), APC anti-human CD39 (328209), Brilliant Violet 605™ anti-human CD56 (318334), PE anti-human CD16 (360704), PerCP/Cyanine5.5 anti-human CD19 (302230), and PerCP/Cyanine5.5 anti-human CD14 (325622). These latter two antibodies, conjugated to the same fluorochrome, were utilized as a dump channel to exclude B cells and myeloid cells. To thoroughly remove unbound antibodies and minimize non-specific background fluorescence, cells were washed two times with 1 mL of Cell Staining Buffer and finally resuspended in 300 µL of Cell Staining Buffer.

Samples were acquired on a four-laser Sony SA3800 spectral flow cytometer using a standardized template across all samples (Fluorescence PMT voltage 68.1%, FSC gain 17, SSC voltage 30.6%). A stopping gate was set to record 70,000 CD45^+^ cells. Samples with high cellularity were acquired across multiple tubes and subsequently concatenated.Spectral unmixing was performed utilizing reference controls exhibiting the highest spectral index for each fluorochrome within the library. Due to the continuous expression profile of PD-1, fluorescence minus one controls and positive controls (stimulated PBMC CD8⁺ T cells) were jointly utilized to establish precise gating thresholds. Background fluorescence was corrected using the multiple autofluorescent populations mode. Data variance was further minimized via Flow Point Correction, which calculates unmixing based on cell position information. Manual adjustments were applied via the spectral reference adjuster to correct any imperfect auto-unmixing results.

### Single-cell RNA-seq (scRNA-seq) data generation

#### 1) In-house cohort

Leveraging our group’s clinical specialty in thoracic and head and neck oncology, and to address the relative scarcity of certain cancer types or subtypes in current literature, we collected a cohort of treatment-naive specimens comprising 23 driver-mutant non-small-cell lung cancer (NSCLC) (predominantly advanced/late-stage EGFR-mutant), five thymic tumors, four nasopharyngeal carcinomas (NPC), and one lung lymphoepithelioma-like carcinoma (Table. S1). One tumor located in the thymic region was later pathologically confirmed as a B-cell lymphoma. The study was approved by the Research and Ethics Committee of the Sun Yat-sen University Cancer Center [B2024-334-01, B2024-335-01, B2025-502-01, G2025-018-01, B2025-899-01, and B2020-402-Y02]. Fully informed written consent was obtained from all participants prior to sample collection.

Fresh tumor tissue beyond standard diagnostic requirements was obtained via surgical resection, CT/ultrasound-guided biopsy, or endoscopy. During sampling, non-target tissues (such as surrounding normal tissue, excess fat, and blood clots) were removed. The samples were placed in MACS Tissue Storage Solution (Miltenyi Biotec) on ice for preservation and transport until the preparation of single-cell suspensions. Tissue was washed with DPBS (PBS without calcium and magnesium) and chopped into small pieces. Enzymatic digestion was performed with collagenase I (2 mg/mL), collagenase II (2 mg/mL), collagenase IV (2 mg/mL), and dispase (0.2 mg/mL) at 37℃ for 15-25 minutes. The digested suspension was filtered through a 70-μm nylon mesh (Greiner Bio-One GmbH, Germany). The filtrate was immediately centrifuged at 500 g for 5 min, and the supernatant was slowly discarded. The cell pellet was resuspended and lysed with RBC Lysis Buffer on ice for 5 min to remove red blood cells, followed by washing twice with DPBS at 400 g for 5 min. Cell concentration and viability were determined using a Cellometer Auto 2000 (Nexcelom, USA) following AO/PI staining.

Single cells were processed using the Chromium Next GEM Single Cell 5’ Kit v2 (10x Genomics) following the manufacturer’s protocol with modifications. Approximately 20,000 cells were loaded into each channel, resulting in the capture of approximately 8,000 cells per sample. The scRNA-seq libraries were synthesized according to standard 10x Genomics protocols and sequenced on the NovaSeq X Plus platform (Illumina) to a depth of approximately 330 million reads per library, utilizing the following read configuration: Read 1: 28 cycles; Read 2: 96 cycles; Index 1: 8 cycles.

FASTQ files were aligned and quantified using Cell Ranger (v7.1.0) against the GRCh38 reference genome. To maximize transcriptional sensitivity, we considered both exonic and intronic reads (--include-introns=true). To prevent the premature loss of biologically significant cell populations characterized by intrinsically low counts, the capture threshold was forced to 25,000 cells per sample (--force-cells=25000). Although this permissive threshold retains a fraction of low-quality barcodes or ambient RNA profiles during initial matrix generation, these artifacts will be rigorously identified and excluded during downstream quality control (see section “Shared scRNA-seq data processing methodologies”).

#### 2) Lung discovery cohort

A systematic search of PubMed and GEO was performed to identify relevant datasets published from inception to June 2024. The search strategy utilized keywords and Medical Subject Headings (MeSH) terms encompassing three primary domains (e.g., sequencing modalities such as “single-cell” or “scRNA” and anatomical and disease terminology such as “lung”, “NSCLC”, “adenocarcinoma”, or “metastasis”. Notably, approximately half of the data included in our lung discovery cohort are controlled-access and not publicly available. To address this, we negotiated with the respective laboratories to acquire the raw sequencing data for our analysis. The inclusion criteria were as follows: treatment-naive patients diagnosed with NSCLC who had not received prior systemic therapy or radiotherapy, with available scRNA-seq or snRNA-seq data. Samples that were redundantly utilized across different studies were strictly excluded.

We invested significant efforts in curating comprehensive clinical annotations, where available, including age and sex, pathological type and histological grade, tumor stage, race/region, smoking history, tumor location, radiological classification, and the status of driver mutations. In instances of co-mutations, the prioritization hierarchy was based on the current clinical practice landscape of NSCLC. Priority was first given to clinically relevant actionable genomic alterations (EGFR/KRAS/ALK/HER2/MET/RET/BRAF) and key tumor suppressor genes (STK11/KEAP1/SMARCA4/TP53), followed by other research-relevant mutations (including MUC16, KMT2, ARID1A/B, LRP1B, ATM, and PTPRD). The remaining patients were classified as lacking common mutations. Furthermore, the EGFR and KRAS mutant groups were sub-stratified based on the presence of concomitant TP53 mutations, and the EGFR group was additionally sub-stratified by specific mutant alleles. Only pathogenic or likely pathogenic mutations were considered. In cases where the treatment-naive status of the samples was ambiguous, or when clinical annotations could not be matched with the corresponding expression data, these discrepancies were reconciled and confirmed through correspondence with the original authors.

For studies that provided whole-exome sequencing FASTQ files without accompanying mutation profiles, we performed somatic variant calling according to GATK best practice (55). Briefly, pre-trimmed, paired-end sequencing reads from matched tumor and normal/PBMC samples were aligned to the GRCh38 reference genome using the BWA-MEM algorithm. PCR and optical duplicates were identified and removed from the BAM files utilizing the MarkDuplicates function within the Picard toolkit. Empirical base quality scores were recalibrated against known variant sites from dbSNP (build 138) using the BaseRecalibrator and ApplyBQSR modules to correct for systematic sequencing errors. Somatic single nucleotide variants and short insertions/deletions were identified using Mutect2, leveraging a gnomAD-derived germline resource (allele-frequency only) and a 1000 Genomes Project-derived Panel of Normals. The VCF file was processed through FilterMutectCalls, which integrated contamination tables and orientation bias models to compute the final probability of each variant being a true somatic mutation. High-confidence somatic calls were subsequently annotated utilizing ANNOVAR.

Ultimately, the cohort comprised 343 patients and 430 tumor samples, including 152 matched adjacent normal tissue samples (Table. S1 and Fig. S1). The majority of samples were derived from lung adenocarcinomas (90.5%) and primary tumor sites (89.8%). Approximately half of the samples were obtained from patients aged over 60 years (53.5%), female patients (47.2%), and current or former smokers (48.6%). All samples were accompanied by genetic profiling data. The majority of samples were sequenced using the 10x Genomics platform (79.8%) and were not subjected to prior flow cytometry sorting (85.6%).

To generate gene expression matrices, we processed raw FASTQ files whenever provided by the authors; otherwise, we directly utilized the expression matrices. Data processing for the 10x Genomics platform was consistent with our in-house cohort pipeline (see section “In-house cohort”). Other sequencing platforms requiring upstream analysis included BD Rhapsody, SMART-seq2, and Drop-seq. BD Rhapsody data were analyzed using the BD Rhapsody Sequence Analysis Pipeline (version 2.0), with alignment and quantification performed against the Human Whole Transcriptome Assay reference archive. Both SMART-seq2 and Drop-seq data were aligned to the GRCh38 reference genome and quantified utilizing the STARsolo algorithm (56) (--soloType SmartSeq and --soloType CB_UMI_Simple, respectively). Two core principles were maintained across all platforms: (1) the inclusion of intronic transcripts for quantification (enforced via Exclude_Intronic_Reads: false in the BD Rhapsody pipeline and --soloFeatures GeneFull_Ex50pAS in the STARsolo pipeline), and (2) the implementation of a permissive upstream cell-calling strategy (Exact_Cell_Count: 25,000 for BD Rhapsody, --soloCellFilter None for SMART-seq2, and --soloCellFilter TopCells 1000 for Drop-seq). Because SMART-seq2 libraries do not incorporate unique molecular identifier (UMI), deduplication was disabled (--soloUMIdedup Exact NoDedup), and read quantification was performed in an unstranded manner (--soloStrand Unstranded). For Drop-seq data, PCR duplicates were resolved utilizing a 1-mismatch collapse strategy analogous to the Cell Ranger algorithm (--soloUMIdedup 1MM_CR).

#### 3) Pan-cancer cohort

Although some pan-cancer scRNA-seq databases have been constructed, they frequently mix pre- and post-treatment samples and lack detailed clinical annotation. Pre-treatment samples reflects the actual immunological state of the tumor, which is shaped by long-term growth and intrinsic immune pressures, unaffected by therapeutic interventions. Therefore, we established a pan-cancer database strictly comprising treatment-naive tumor samples, featuring a larger sample size and a more comprehensive range of cancer types than previous atlases.

A systematic search of PubMed and GEO was performed to identify relevant datasets published up to October 2025. The search strategy was similar to that of the lung discovery cohort but expanded to encompass a broader array of organ and cancer terminology (e.g., “neoplasm”, “carcinoma”, “sarcoma”, “glioblastoma”, or “melanoma”). Furthermore, we manually searched and evaluated previously published single-cell atlas, such as the Curated Cancer Cell Atlas (3CA; https://www.weizmann.ac.il/sites/3CA/), CZ CELLxGENE Discover (https://cellxgene.cziscience.com/datasets), Tumor Immune Single-cell Hub (TISCH; http://tisch.comp-genomics.org/), and the Human Tumor Atlas Network (HTAN; https://data.humantumoratlas.org/explore). Datasets already included in the lung discovery cohort were excluded.

The inclusion criteria required treatment-naive cancer patients with available scRNA-seq or snRNA-seq data for tumor tissues. Precancerous tissues were also considered eligible if reported within the included cancer studies. However, normal adjacent tissues or blood samples were strictly excluded. Additionally, the samples were required to be either unsorted (containing all viable cells) or at least contain the CD45^+^ leukocyte fraction. To improve efficiency, only processed expression matrix data were included. Five investigators independently and in duplicate screened, collected, and verified the pan-cancer clinical information. If an inconsistency arose, a consensus was reached through discussion among all investigators. The collected data included cancer type, clinical stage, and anatomical site (primary or metastasis). Specific subgroup information was recorded where available, including: molecular subtypes for lung cancer; microsatellite instability status for colorectal cancer; pathological subtypes for breast cancer; homologous recombination status for ovarian cancer; viral infection status for liver hepatocellular carcinoma and head and neck squamous cell carcinoma; and isocitrate dehydrogenase (IDH) mutation status for glioblastoma. Ultimately, 3,603 samples were included; when combined with our in-house cohort and the lung discovery cohort, the final pan-cancer database reached 4,066 samples and 2,748 patients. Detailed cancer type information can be found in Fig. 3A and Table. S1.

To correlate this killer divergence with pan-cancer immune checkpoint inhibitor (ICB) efficacy at the cancer-type level, we collected objective response rates (ORRs) for each included malignancy. Mao et al. (1) and Yarchoan et al. (57) previously curated and calculated the average ORR for several cancer types. For cancer types reported by both prior studies, we utilized the mean of their reported ORRs as the final average ORR; if reported by only one study, that specific ORR was utilized. A substantial number of cancer types included in our pan-cancer cohort were not covered by these prior works. For these remaining malignancies, we conducted a systematic search utilizing inclusion criteria analogous to their works (1,57). Briefly, we excluded studies investigating combination therapies or selecting patients based on immune-related biomarkers such as PD-L1 expression. Generally, studies enrolling fewer than 10 participants were excluded; however, in scenarios where only trials with <10 participants were available for a specific cancer type, those data were retained and noted in Table. S4. We calculated the average ORR for each cancer type by dividing the total number of responders by the total number of patients receiving ICB monotherapy. Finally, cancer types were stratified into four ORR categories (<5%, 5–10%, 10–20%, and >20%).

#### 4) Immunotherapy cohort

Based on our comprehensive review of the current literature, most studies evaluating biomarkers for immunotherapy at the single-cell level suffer from at least one of the following drawbacks: (1) small sample sizes in individual scRNA-seq datasets; (2) a reliance on bulk RNA-seq data to establish efficacy associations via signature scoring or deconvolution; (3) the frequent employment of cell-level signature scores rather than sample-level analyses to evaluate biomarker predictiveness, a methodological flaw that introduces statistical pseudo-replication; and (4) the mixing of pre- and post-treatment samples, or an exclusive reliance on post-ICB tissues, where predictiveness may be conflated by treatment effects. To address these limitations, we aimed to compile a large-scale ICB scRNA-seq cohort, carefully stratifying samples by their acquisition timepoints.

A systematic search of PubMed and GEO was performed to identify relevant datasets published from inception to Nov 2025. The search strategy utilized keywords and MeSH terms encompassing three primary domains (e.g., sequencing modalities such as “Transcriptome” or “RNA-Seq”; immunotherapeutic targets/drugs such as “Immune Checkpoint Inhibitors”, “PD-1”, “Pembrolizumab”, “adoptive cell therapy”, or “CAR-T”; and cancer terminology such as “Neoplasms” or “carcinoma”).

We included tumor samples from patients treated with ICB monotherapy or combination strategies with available scRNA-seq or snRNA-seq data. We only considered baseline pre-treatment samples or strictly site-paired longitudinal samples. Datasets were required to report pathologic or radiologic response data or other surrogate endpoints such as T cell clonal expansion. For all included samples, we recorded the cancer type, precise sampling timepoints, treatment regimens, and response. This database contain 628 samples from 376 patients across 13 cancer types (Table. S1 and Fig. 5A). We also included 18 pre-treatment samples from melanoma and GBM patients treated with adoptive T cell therapy (Table. S1).

### Bulk RNA-seq data generation

#### 1) In-house clinical trials

This study includes three in-house trials of ICB (Table. S1): two randomized controlled trials (RCTs) (ORIENT-11 [NCT03607539] and RATIONALE-309 [NCT03924986]) and one non-randomized trial [NCT02516527/NCT02721589]. Detailed clinical settings and RNA-seq methodologies have been described in previous publications (42,43,58). The protocols were approved by the local ethics committee for each trial, and all participants provided written informed consent. These trials were conducted in accordance with the Declaration of Helsinki.

In brief, ORIENT-11 was a double-blind, phase 3 study that enrolled patients from 47 centers in China who had previously untreated locally advanced/metastatic nonsquamous NSCLC without sensitizing EGFR mutation or ALK translocation. Patients were randomized 2:1 to receive the anti–PD-1 antibody sintilimab or placebo plus pemetrexed and platinum every 3 weeks (Q3W) for four cycles, followed by sintilimab or placebo plus pemetrexed as maintenance therapy. RATIONALE-309 was a phase 3, multicenter, randomized, double-blind, placebo-controlled clinical trial conducted at 42 sites in Asia. Patients with recurrent or metastatic NPC were randomly allocated (1:1) to either tislelizumab 200 mg or matching placebo Q3W, plus the chemotherapy regimen gemcitabine and cisplatin. Gemcitabine 1 g/m2 was given on Day 1 and Day 8, and cisplatin 80 mg/m2 on Day 1. The chemotherapy regimen was administered Q3W for 4 to 6 cycles, at the investigators’ discretion. The non-randomized trial assessed sequential anti-CTLA-4 and anti-PD-1 therapies in patients with advanced/metastatic NSCLC and NPC who had failed at least two lines of systemic treatment. Patients received 3 or 10 mg/kg ipilimumab Q3W for up to 4 cycles then entered a maintenance phase every 12 weeks starting at week 24, followed by camrelizumab treatment. The washout period was 4 weeks or longer.

These three trials included a total of 432 patients assessable for both efficacy and RNA-seq data, comprising 171 from ORIENT-11 (sintilimab plus chemotherapy, n = 113; chemotherapy, n = 58), 247 from RATIONALE-309 (tislelizumab plus chemotherapy, n = 124; chemotherapy, n = 123), and 14 from the non-randomized trial (7 NSCLC and 7 NPC).

#### 2) Public immunotherapy cohorts

The search strategy for these datasets was similar to that for the scRNA-seq immunotherapy cohorts. The inclusion criteria were as follows: 1) assessment of ICB in human cancer, with a minimum of six patients receiving ICB per cohort; 2) availability of clinical outcome data for at least one of three categories: response, disease control endpoints (DCE, such as progression-free survival, event-free survival, or disease-free survival), or overall survival. For response evaluation, cohorts were required to have at least three responders and three non-responders; and 3) reporting of bulk RNA-seq or microarray expression data. For all included datasets, we extracted cancer type, efficacy results across the aforementioned three endpoints, and treatment information. Ultimately, 6,914 public samples were collected; when combined with our in-house data, this yielded a comprehensive cohort of 7,346 patients across 33 cancer types (Table. S1 and Fig. 5A).

If processed expression matrices were provided by the authors, we directly utilized them. We imposed no restrictions on the specific normalization methods utilized in the original papers, because our meta-analysis strategy evaluated each cohort independently (see section “Immunotherapy efficacy analysis”), thereby circumventing cross-cohort batch effects. However, when applicable, we transformed all data to the log2 transcripts per million format. For raw RNA-seq data, reads were subjected to adapter trimming and quality filtering using Trim Galore (59). Low-quality base calls were removed based on a Phred33 quality score threshold of < 25. Sequence reads were discarded if they were reduced to fewer than 20 base pairs. Quality-filtered reads were aligned to the GRCh38 reference genome utilizing the HISAT2 algorithm (60). Read counting and gene-level expression quantification were performed using featureCounts (61). For raw microarray CEL data, probe intensities were processed using the Robust Multi-array Average algorithm to generate a normalized probe-level expression matrix. Probe identifiers were mapped to gene symbols utilizing the hgu219.db and hgu219cdf R packages.

#### 3) Public non-immunotherapy cohorts

Normalized pan-cancer transcriptomic datasets were obtained from The Cancer Genome Atlas (TCGA; n = 9,807 samples) (62) (https://toil-xena-hub.s3.us-east-1.amazonaws.com/download/tcga_RSEM_gene_tpm.gz), the Clinical Proteomic Tumor Analysis Consortium (CPTAC; n = 1,083 samples) (63) (https://pdc.cancer.gov/pdc/cptac-pancancer; file: RNA_WashU_v1.zip), and the University of California, San Francisco (UCSF) Immunoprofiler Initiative (UCSF-IPI; n = 260 samples) (64) (GEO dataset GSE184398; file: GSE184398_pancan_all_pc_genes_Live_TPM_Aug_3_20.tsv.gz). Unlike the TCGA and CPTAC datasets, the UCSF-IPI dataset exclusively sequenced viable cells isolated following enzymatic tissue digestion. Cancer type names were standardized across all three cohorts (Table. S1).

### Spatial sequencing data generation

Our analytical goal was to validate the spatial and sample-level relationship between Tex and NK1 in situ. However, based on our comprehensive review of the current literature regarding the application of spatial transcriptomics in tumor samples, existing spatial technologies excel at characterizing malignant and stromal cells (e.g., fibroblasts and endothelial cells) but face significant challenges in capturing T/NK cell subsets, particularly for NK cells.

A recent benchmarking study (65) suggested that image-based platforms outperform sequencing-based platforms in detecting lymphocytes. Xenium achieved optimal performance regarding marker capture sensitivity, a lower proportion of negative control signals, and better annotation robustness compared to other platforms. Importantly, Xenium yielded the most distinct and biologically coherent expression patterns for T/NK cells.

Using a search strategy similar to that of the lung discovery scRNA-seq cohort but focused on spatial transcriptomics rather than scRNA-seq, we identified several available platforms in current public NSCLC datasets, including image-based Xenium, CosMx, and MERSCOPE, as well as sequencing-based Visium. We found no published spatial proteomics data (i.e., IMC, CODEX, MIBI) containing suitable markers to concurrently define both NK1 and Tex.

Our subsequent evaluations of different platforms demonstrated that Xenium rather than MERSCOPE, CosMx, and Visiumcan can robustly capture a substantial number of Tex and NK1 cells using a customized strategy (see section “Xenium data analysis”) (data not shown for other platforms). Additionally, the effective tissue capture area of Xenium is generally larger than other platforms, thereby providing a more accurate representation of sample-level heterogeneity. Consequently, we selected the Xenium platform for our analysis. The utilized data were derived from Takano et al. (66) (302-gene panel) and from the official 10x Genomics dataset resource (https://s3-us-west-2.amazonaws.com/10x.files/samples/xenium/3.0.0/Xenium_Prime_Human_Lung_Cancer_FFPE /Xenium_Prime_Human_Lung_Cancer_FFPE_outs.zip; 5k-gene panel), resulting in a total of 18 samples (Table. S1).

### Shared scRNA-seq data processing methodologies

All analyses were performed utilizing the Seurat (67) (version 5.4.0) or scanpy (68) (version 1.9.8) framework.

#### 1) Quality control (QC)

For cohorts utilizing multiplexed configurations, raw matrices were filtered to retain only cell barcodes containing detectable counts within the Hashtag Oligonucleotide (HTO) feature space. HTO count matrices were normalized utilizing a Centered Log-Ratio transformation. Sample demultiplexing was executed employing HTODemux command based on a threshold of 0.99. Any barcodes classified as “Negative” (insufficient hashtagging) or “Doublet” (discordant multi-sample tagging) were excluded from the transcriptomic matrices. The remaining singlets were partitioned by their experimental origins.

QC was performed independently on a per-sample basis. QC metrics can inherently vary across distinct physiological cell states—for instance, cells exhibiting a relatively high fraction of mitochondrial counts might be involved in essential respiratory processes, whereas those with aberrant overall transcript counts could represent naturally quiescent populations or larger cells. Therefore, our QC strategy integrated hard thresholds with cluster-guided dynamic filtration. Furthermore, while cells exhibiting high expression of ribosomal, dissociation-related, or heat-shock protein genes may represent low-quality captures, cumulative evidence suggests that they can form biologically meaningful cell populations. Therefore, we did not perform QC filtration based on these metrics. Such populations are annotated as isolated clusters during downstream unsupervised clustering and do not compromise the analytical integrity of the primary cell types of interest.

Cells were retained if they contained > 200 detected unique genes and an mitochondrial transcript proportion of < 20%. To implement the cluster-specific dynamic QC, the surviving cells underwent preliminary clustering. Raw counts were normalized utilizing SCTransform using the top 2,000 highly variable genes (HVGs). The number of principal components (PCs) and the resolution for clustering were dynamically assigned on a per-sample basis to account for variable cell yields. Within each cluster, the median and median absolute deviation (MAD) were calculated for three critical covariates: unique gene counts, total transcript counts, and mitochondrial transcript percentages. Any cell deviating by more than 3 MADs from its respective cluster-specific median for any of these parameters was classified as low-quality and discarded. Concurrently, to eliminate erythrocyte contamination, any cell exhibiting a proportion of hemoglobin-related transcripts exceeding the 99th percentile globally across the sample was removed. Following cluster-specific filtration, each sample was re-normalized and re-clustered for subjecting to DoubletFinder (69) to remove doublets. The expected doublet rate was scaled to the cell recovery rate and adjusted to account for the proportion of homotypic doublets within the refined clusters.

#### 2) Dimensionality reduction, batch integration, and clustering

We performed unsupervised clustering and re-clustering hierarchically, progressing from broad cell lineages to granular cell subsets. Samples containing fewer than 20 cells for a specific cell type were excluded from the downstream analysis of that cell type. Gene symbols exhibiting poor sample coverage—defined as having zero expression across more than 25% of the total samples—were removed from the dataset. Because proliferating cells from distinct lineages frequently cluster together, clustering and annotation for proliferating populations were conducted independently from the non-proliferating cellular compartments.

Cell-by-gene matrices were normalized by SCTransform using the top 3,000 HVGs; however, this threshold can be adjusted based on cell yield and dataset complexity. Initial dimensionality reduction was performed via Principal Component Analysis (PCA). Typically, 30 PCs were calculated, but this parameter was similarly tailored to the specific cellular yield and complexity of each sample. In the first round of B cell clustering (B_memory, B_naive, ASC, GCB), to prevent transcriptomic heterogeneity from being predominantly driven by the high expression of immunoglobulin transcripts, all Ig genes were excluded from the HVG list. To correct for batch effects, the Harmony integration algorithm was applied to the PCA embeddings (70). A shared nearest-neighbor (SNN) graph was constructed utilizing the Harmony-corrected embeddings, followed by unsupervised clustering (FindClusters) and Uniform Manifold Approximation and Projection (UMAP) for visualization. To determine the optimal cluster granularity, clustering outputs across multiple resolutions were visualized utilizing the clustree R package.

When clustering massive cellular compartments (e.g., T/NK or myeloid lineages), which incur heavy computational memory costs and extended processing times, three high-efficiency strategies were deployed: 1) Rather than integrating all cell types across all cohorts simultaneously, each cohort was initially analyzed independently to annotate broad cell lineages. Specific broad cell types were subsequently isolated, merged across cohorts, and re-clustered. 2) The BPCells R package was utilized to transition cell matrices into on-disk storage. 3) Individual cells were aggregated into metacells utilizing the SEACells algorithm (71). Dimensionality reduction, batch integration, clustering, and low-granularity annotation were performed at the metacell level, and the resulting annotations were subsequently mapped back from the metacell space to the original single-cell resolution for further clustering and analysis.

#### 3) Cell type annotation

Broad cellular compartments were defined based on the expression of canonical marker genes as follows: epithelial cells (EPCAM, KRT19, SFTPC), T and natural killer (NK) cells (CD3D, CD8A, CD4, GNLY, PRF1), B and plasma cells (CD19, CD79A, JCHAIN, MS4A1, MZB1), myeloid cells including macrophages, monocytes, dendritic cells, and neutrophils (CD68, CD14, LYZ, C1QB), mast cells (TPSAB1, TPSB2), mesenchymal cells (COL1A1, COL1A2, DCN, FAP), and endothelial cells (VWF, PECAM1) (Fig. S2).

Granular cell subsets were annotated utilizing a integrative approach incorporating the following criteria: (1) Because scRNA-seq data are inherently sparse due to dropout events, differentially expressed genes (DEGs) defining each sub-cluster were identified based on sample-level pseudobulk derived from AggregateExpression function. Significant DEGs were calculated from these pseudobulk matrices via the FindAllMarkers function using a Wilcoxon rank-sum test. (2) The top 50 highest-ranking DEGs by their average log_2_ fold-change, were submitted to the ACT web server to guide objective cell assignments (72). (3) Functional gene signatures and markers established in previous pan-cancer single-cell atlas were visualized using FeaturePlot and Dotplot. (4) The spatial proximity of clusters within the UMAP embedding was evaluated to indicate relationships: Functionally related cellular states cluster contiguously in reduced space, whereas spatially distant clusters represent distinct phenotypes.

Multi-layered nomenclature was retained for cells exhibiting phenotypic hierarchies (e.g., broad annotation as Tex, accompanied by granular state annotations such as T_CD8_Tex_CXCL13 or T_CD8_Tex_GZMB). Cells with ambiguous identities—characterized by elevated mitochondrial or ribosomal fractions, or uniquely low transcript counts—were specifically flagged. Despite execution of DoubletFinder (69), clusters exhibiting co-expression of highly divergent lineage markers were further manually classified as doublets and removed. For annotation in pan-cancer scRNA-seq cohort, please see section “Automatic cell type annotation pipeline”.

#### 4) Sample-level comparative analysis

To mitigate the statistical bias of pseudoreplication inherent in single-cell datasets when performing sample-level comparative analyses, scRNA-seq profiles were transformed into pseudobulk utilizing the normalized data slot via the aggregateAcrossCells function (73). Signature scores were quantified by calculating the mean expression of the respective gene lists for each sample. By default, the analysis of the signature was restricted to datasets generated using the 10x Genomics sequencing platform with UMI counts; however, samples generated using other methods were treated as independent validation subgroups.

### Landscape of cytotoxic lymphocytes (killers)

#### 1) Bulk RNA-seq analysis

Twenty-five killer cell functional signatures were curated from published literature and the Molecular Signatures Database (MSigDB) (Table. S2). Although multiple signatures have been developed for NK and CD8⁺ T cells in the literature, we utilized the ones explicitly defined as cell-type-restricted by original authors to more accurately reflect different cell types in bulk RNA-seq data. Prior to scoring, the gene sets were deduplicated to eliminate gene overlap. Specifically, any gene present in multiple signatures was assigned exclusively to the signature comprising the fewest genes.

For the majority of the signatures, scores were calculated as the mean of the normalized expression. However, the MCPcounter signature was calculated using the MCPcounter package (74), and global tumor immune infiltration (ESTIMATE ImmuneScore) was quantified by ESTIMATE algorithm (75). Pairwise Spearman correlations were calculated across the signatures independently within the TCGA, CPTAC, and IPI cohorts. Given the substantial positive correlation observed among the signatures, a unified metric representing the global killer state was derived using the first principal component (PC1) score from a PCA.

#### 2) scRNA-seq analysis

We sought to understand the primary dimensions driving CD8⁺ and NK cell phenotypes and differentiation from both continuous and discrete perspectives. To reconstruct the developmental continuum, diffusion maps was executed utilizing the sc.tl.diffmap function in scanpy. To understand the differences among cell subsets, we performed unsupervised clustering based on curated markers and signatures (Table. S2) and visualized the results using a heatmap. Distinct from most studies that directly compare subsets based on single-cell scoring, we generated sample-level pseudobulk matrices for each cell subset. Expression values were subsequently averaged across all samples to generate a representative profile for each subset. This approach takes advantage of our large sample size to mitigate bias introduced by individual sample variance.

Notably, for NK cells, we found that unsupervised clustering inevitably resulted in a cluster (NK_Tlike) expressing T cell markers such as CD3D and CD8A, even after combining DoubletFinder (69) and manual strategy excluding doublets. There are two primary approaches for identifying NK cells in the current literature: the first, and most widely utilized approach—which we employed in our study—involves an unsupervised clustering strategy followed by annotation based on canonical NK markers such as GNLY. The second strategy, utilized in a NK landscape study (20), employs an in-silico gating strategy using NCAM1 and KLRF1 to select NK cells while explicitly excluding T cell-related markers. We ultimately selected the first strategy based on the following rationale: (1) Although the conventional definition of NK cells requires them to be CD3-negative, more recent evidence highlights the existence of naive-like NK cells, adaptive memory-like NK cells (also termed NK3), NKT cells or other cell populations with ambiguous identities, which could all validly express T cell markers. The gating strategy might erroneously exclude these biologically relevant proportions. (2) The gating strategy risks overestimating the proportion of CD56^dim^CD16^hi^ NK1 cells while underestimating CD56^bright^CD16^low^ NK2 cells at the sample level. This bias arises because NCAM1 (CD56) expression, which is higher in NK2 cells, is often artificially low due to scRNA-seq dropout events, whereas KLRF1 expression, which is elevated in NK1 cells, is more effectively captured. (3) By including the NK_Tlike cluster, we retain the analytical flexibility to either keep or remove it during specific analyses, achieving results equivalent to the gating strategy when necessary. Taken together, while the in-silico gating method can ensure the purity of classical NK cells, it may distort the internal NK infiltration proportions at the sample level. Because the primary objective of our study is to illustrate killer pattern across tumors, we selected the unsupervised clustering approach.

### Uncover killer divergence in NSCLC

#### 1) scRNA-seq analysis

We evaluated the killer pattern across samples via PCA using the granular-layer cell proportions of CD8⁺ T cells and NK cells (Table. S6). PCA identifies uncorrelated PCs that are linear combinations of the original variables, with the first PC capturing the maximum variance across the dataset. The proportion of each cell subset was transformed into non-parametric ranks utilizing an average tie-breaking method, and were centered and scaled to unit variance. PCA was executed using the prcomp function. Cell proportion were calculated by default relative to each corresponding parent cellular compartment (e.g., the denominator for Tex was total CD8⁺ T cells, and for NK1, total NK cells). The advantage of this metric is that it isolates the internal cellular programming and differentiation without being confounded by unrelated cell types.

To evaluate the differentiation relationships of CD8⁺ T cells and NK cells across tumors, the primary cell-level diffusion component (DC1) was linked to the sample-level PC1. First, Spearman’s correlation coefficients were calculated between the normalized expression of each gene and the DC1 scores, performed independently for the NK and CD8⁺ T cell populations. Genes demonstrating a significant positive correlation (FDR < 0.05 and corr > 0.4) were extracted as key genes defining the DC1 developmental axis. These identified genes were utilized to quantify DC1 scores at the pseudobulk level and were subsequently modeled along the scaled PC1 axis utilizing a generalized additive model (GAM) with cubic regression splines, interpolated across a 500-point grid.

Robustness and sensitivity analyses of our findings were confirmed across clinical subgroups and through justified adjustments to analytical parameters. We evaluated the stability of the PCA results when utilizing an imputation versus a non-imputation strategy for missing cellular fractions; this involved comparing a primary strategy where missing values were imputed with the cell proportion mean against a strict complete-case validation where samples containing any missing cell type information were excluded. Furthermore, sensitivity was assessed by evaluating the retention versus exclusion of clusters exhibiting high mitochondrial or ribosomal gene expression, testing the inclusion versus exclusion of the ambiguous NK_Tlike subset, and stratifying the cohorts by sequencing platform or cell-sorting methodology.

Although utilizing the respective NK and CD8⁺ T cell compartments as denominators effectively prevent compositional effects, this approach does not reflect the overall infiltration level of Tex and NK1 within the entire tumor sample. Therefore, to determine if the observed divergence persists at the global tissue level, we re-evaluated our analyses by changing the denominator to either total CD45⁺ immune cells or all captured cells. For this global-denominator analysis, we excluded samples exhibiting systematic cell-type biases, such as those processed via targeted flow cytometry sorting or those with an explicit absence of major cellular lineages.

#### 2) Spectral cytometry analysis

Following spectral unmixing, a hierarchical gating strategy was applied (Fig. S11). Lymphocytes were enriched based on forward and side scatter (FSC/SSC) properties. Viable leukocytes were subsequently identified by gating for CD45 positivity and the absence of a viability dye. B cells and myeloid lineages were excluded using a “dump” channel (CD14/CD19 negative). The remaining viable lymphocytes were bifurcated into NK cells (CD3^-^, CD56^+^) and T cells (CD3^+^, CD56^-^). Within the T cell compartment, CD8^+^ T cells were further gated. Tex were defined as the double-positive population expressing both PD-1 and CD39 within the CD8+ T cell gate. NK1 cells were defined by CD16 positivity within the NK cell compartment. Event counts for all target populations and reference denominators were summed across technical replicates. The frequencies of the Tex and NK1 subsets were calculated across three distinct denominators: (1) direct parent lineages (CD8^+^ T cells or NK cells); (2) total T/NK cells; and (3) total CD45^+^ cells.

To evaluate the correlation between the frequencies of Tex and NK1 cells across the cohort while controlling for technical and biological confounders, linear mixed models (LMMs) were utilized. Correlation assessments were performed at both the sample level and the patient level. At the sample level, both patient identity and acquisition date were modeled as random intercepts. For patient-level analysis, the acquisition date was retained as a random intercept. For all LMMs, the fixed-effect slope was extracted to determine the direction and magnitude of the correlation, with significance derived via standard Satterthwaite approximations.

#### 3) Xenium data analysis

All analyses were performed utilizing the Seurat (version 5.4.0) framework (67). QC of the Xenium data was performed independently for each sample. Four primary metrics were evaluated per cell: total transcript counts, unique features detected, cell area (µm²), and transcript count density (defined as the total transcript counts divided by the cell area). All four metrics were initially log_2_-transformed. Cells were excluded from downstream analysis if any of their metrics fell below 4 MADs or exceeded 5 MADs from the sample-specific median. In addition to these dynamic thresholds, a minimum threshold of 20 unique features was required for samples analyzed with the 5k gene panel, whereas a minimum of 5 unique features was applied for the 302-gene panel. To control for non-specific probe binding and optical decoding anomalies, a ratio-based noise evaluation strategy was implemented. For every cell, total noise events were aggregated by summing the counts of all detected blank codewords, control codewords, and control probes. A noise ratio was then calculated by dividing this total noise sum by the overall count. Cells exhibiting a noise ratio exceeding 5% were discarded. All discarded cells were mapped back onto the field of view to confirm a random distribution pattern across the tissue or aggregation strictly at borders, as a biologically organized spatial pattern might indicate the overly strict exclusion of valid cell populations.

Identifying malignant cells based on the restricted scale of gene panels in Xenium data is challenging, which precludes the robust inference of DNA copy number variations (CNVs). Additionally, our evaluation utilizing SingleR with a standard lung malignant cell signature demonstrated a propensity for false-negative classifications upon histological review. Consequently, a dual-modality approach integrating histopathological examination with unsupervised clustering was developed to ensure high-confidence tumor cell identification. Two experienced pathologists performed detailed histopathological examinations of H&E-stained sections for each sample to identify tumor regions based on cellular atypia. In parallel, for each sample, spatial count matrices were independently normalized by SCTransform. PCA was executed utilizing the entire gene panel as input features. A SNN graph and UMAP was subsequently constructed. We observed that malignant cells consistently formed a distinct cluster separated from normal epithelial cells in the UMAP embedding. Compared with normal epithelial cells (e.g., AT2), tumor cells often exhibited unique expression profiles—such as elevated expression of cancer-related genes (e.g., VEGFA and CCND1) or driver mutation-related genes (e.g., EGFR)—while showing low expression of lineage-specific markers such as SFTPC, SFTPD, and ETV5.

When attempting to annotate Tex and NK1 cells, we found that the primary PCs were disproportionately driven by non-immune variance, likely attributable to the restricted gene panels (particularly the 302-gene panel) and the overwhelming dominance of malignant and stromal populations within the slice. This phenomenon frequently masked the subtler signals required to resolve CD8^+^ T and NK cell subsets. To maximize the capture of biological variance while excluding confounding lineages, we first utilized in silico gating to enrich for the targeted Tex and NK1 cells. The remaining cells were subjected to repeated iterations of dimensionality reduction, sub-clustering, and annotation based on the coordinated expression of Tex or NK1 genes. We employed two complementary in silico screening processes: a “broad marker” strategy and a “subtype marker” strategy. Under the broad marker logic, a cell was retained if it expressed at least one target broad-lineage transcript (i.e., CD8^+^ T cell or NK cell markers). Under the subtype marker logic, a fractional voting approach was applied to accommodate dropout and biological noise, wherein a cell was retained if it expressed a defined minimal proportion of curated subtype-specific functional markers (i.e., Tex or NK1 markers). Both methods enforced mutual exclusivity between T and NK cells utilizing a list of negative markers. The final Tex and NK1 populations were defined as the collective union of cells surviving either screening pipeline.

Positive marker lists were curated based on spatial panel availability intersecting with DEGs established in our lung discovery cohort. Screening parameters were tailored to the spatial gene panels.

For samples with the 5k gene panel:

- **NK1 Broad Screen:** Positive for NCAM1 or KLRF1; negative for CD8A, CD8B, CD3E, CD3D, and CD3G.
- **NK1 Subtype Screen:** Required expression of at least 1/6 of 24 curated markers (including GZMB, GZMH, PRF1, CST7, CD247, TBX21, S1PR5, RUNX3, GZMA, NCR3, NCR1, PTGDR, DIP2A, ZAP70, KLRG1, KLRC1, SLAMF6, KLRD1, CARD11, CLIC3, CD300A, TXK, SPN, CEMIP2); negative for CD8A, CD8B, CD3E, CD3D, and CD3G.
- **Tex Broad Screen:** Positive for CD8A, CD8B, CD3E, CD3D, or CD3G; negative for NCAM1.
- **Tex Subtype Screen:** Required expression of at least 1/6 of 25 curated markers (including TNFRSF9, BATF, TIGIT, GZMB, CXCL13, ENTPD1, ITGAE, LAG3, TNFSF4, CRTAM, CTLA4, ETV1, TOX2, TOX, RBPJ, ICOS, IL2RG, SIRPG, FASLG, APOBEC3C, CD96, CD200, HAVCR2, APOBEC3G, PDCD1); negative for NCAM1.

For samples with the 302-gene panel:

- **NK1 Broad Screen:** Positive for NCAM1, LCK, or PTPRC; negative for CD8A and CD3E.
- **NK1 Subtype Screen:** Required expression of at least 1/3 of 7 markers (PRF1, TBX21, MYOM2, GZMA, GZMB, SLC14A1, ITGAM); negative for CD8A and CD3E.
- **Tex Broad Screen:** Positive for CD8A, CD3E, LCK, or PTPRC; negative for NCAM1.
- **Tex Subtype Screen:** Required expression of at least 1/4 of 11 markers (CXCL13, PDCD1, LAG3, CTLA4, TIGIT, MKI67, CXCR6, ITGAE, GZMB, GZMA, PRF1); negative for NCAM1.

To reflect the putative ongoing killing process of killer cells against tumor cells, and based on evidence that sequential killing is typically directed at adjacent cells, we mapped Tex and NK1 localized within a 20μm radius of each tumor cell, evaluating up to 10 nearest neighbors utilizing the RANN R package. The exclusivity of such contacts was quantified using a contingency matrix, and statistical significance was confirmed based on the observed-to-expected (Ro/e) interaction frequency ratio alongside permutation testing. The observed frequency of co-targeted tumor cells was compared against an sample-restricted null distribution generated via 1,000 permutations to calculate a single-tailed empirical P value. Spatial contacts were visualized using Xenium Explorer software. Sample-level correlations between the Tex and NK1 were confirmed across two distinct spatial dimensions: the absolute frequency of interactions with tumor cells and the infiltration density. For both space-level and sample-level exclusivity analyses, to normalize against variable tissue slice size and to mitigate topological bias, the metric for each sample was divided by the number of malignant cells localized within a 50μm radius of any identified killer cell, evaluating up to 50 nearest neighbors. This ensured that the analysis was restricted to area where immune contact was physically possible.

### Killer divergence in pan-cancer

#### 1) Automatic cell type annotation pipeline

Manual annotation of scRNA-seq data across nearly 4,000 samples is computationally prohibitive and memory-intensive, necessitating a efficient automated strategy. Our primary objective was to annotate the CD8⁺ T and NK cells to validate the killer divergence framework. However, upon evaluating several automated tools (e.g., SingleR, Seurat label transfer, and SciBet) on selected large cohorts, we observed frequent misclassification between CD8⁺ T and NK cells (data not shown). This ambiguity is likely attributable to the substantial global overlap in their effector transcriptional profiles. To address this, we developed a in-house computational pipeline to accurately annotate these populations by integrating the strengths of both reference-based and marker-based methodologies. This pipeline comprised a three-layer hierarchical strategy executed on a per-cohort basis. Cohorts containing fewer than five samples were merged into a single analytical block if they originated from the same cancer type. Datasets utilizing TPM normalization were processed independently from raw UMI count data.

The first layer was designed to categorize cells into broad lineages. A reference-based strategy was selected because the transcriptomic distinctions between broad cell types are driven by the coordinated expression of many genes. A large-scale reference based on the lung discovery cohort was constructed to include eight major compartments: T/NK cells, B cells, myeloid cells (encompassing dendritic cells, macrophages, monocytes, and neutrophils), mast cells, mesenchymal cells, endothelial cells, and epithelial cells. Annotation was executed utilizing the SciBet R package (76).

For the Layer 2 annotation, we aimed to isolate killer cells as a singular entity (i.e., CD8⁺ T and NK cells) from conventional CD4⁺ T cells and Tregs. At this stage, we did not directly differentiate CD8⁺ T and NK cells because we found that reference-based approaches frequently misclassified them, likely due to their similarly strong upregulation of effector gene network compared to CD4⁺ T cells (i.e., often cluster together in UMAP embeddings). To amplify lineage signals, a metacell reference was established; T/NK cells from the lung discovery cohort were aggregated into pseudobulk metacells utilizing the AggregateExpression function, grouped by sample, cell type (CD8⁺ T cells, NK cells, conventional CD4⁺ T cells, and Tregs), and cohort origin. The top 3,000 HVGs were calculated. To ensure that dimensionality reduction was driven by lineage identities rather than shared, transient cellular states, a curated “black list” of state-associated transcripts was deployed. Genes associated with mitochondrial and ribosomal activity, stress responses, proliferation, interferon stimulation, and specific shared chemokines (e.g., XCL1, XCL2, CXCL13) were explicitly subtracted from the HVG pool. The filtered genes were then utilized to compute PCA embeddings, followed by RunHarmony (70) for batch correction. Query T/NK cells were subjected to the identical “black list” protocol utilized in the reference model prior to PCA and were aggregated into metacells. To identify an optimal metacell partitioning resolution, an iterative SNN clustering loop was deployed. The resolution was adjusted in dynamic increments until the total number of valid clusters fell within a predefined target range proportional to the overall T/NK cell count. Query data were transformed into metacells by aggregating the counts across each unique sample-and-cluster pairing. Metacells were annotated by projecting from the Layer 2 reference using the FindTransferAnchors and TransferData algorithms. A cluster was defined as T_CD8/NK if over 70% of its constituent metacells were annotated as CD8⁺ T cells or NK cells, indicating that the majority of samples within that cluster exhibited a cohesive phenotype.

Layer 3 annotation aimed to differentiate between NK and CD8⁺ T cells. At the resolution of finer cellular subtypes, transcriptional differences become subtle. Consequently, employing a reference-based strategy that relies on broad gene networks becomes susceptible to confounding factors. A simple yet effective method to confirm cell identity relies on targeted, discriminatory genes. GNLY was selected for identifying NK cells due to its robust sensitivity and specificity in previous landscape studies (27). We deployed a rank-based scoring strategy based on the AUCell algorithm (77). Genes were ranked within each cell based on their expression levels. The enrichment of the GNLY was then quantified by calculating the Area Under the Curve (AUC) for each cell. To ensure the analysis was driven primarily by highly expressed, biologically relevant transcripts, the AUC calculation was bounded to consider only the top 2% of the expressed gene rankings within each respective cell (aucMaxRank = 0.02). Cells were annotated by modeling the distribution of the AUC scores across the dataset to establish an automated, objective positive threshold.

However, when attempting to further auto-annotate the Tex and NK1, we found that neither reference-based nor marker-based strategies proved reliable. Therefore, we concluded the automated annotation pipeline at Layer 3 and subsequently utilized gene signatures (Table. S2) to estimate Tex and NK1 levels. To rigorously evaluate the automated pipeline, each cohort outputted a DimPlot, DotPlot, and FeaturePlot for the visual verification of canonical markers across each cell type. In rare instances, manual annotation was required for certain cohorts if the automated process resulted in imperfect classifications.

#### 2) Analysis

The relationship between Tex and NK1 was analyzed at both the sample-level and cancer-level, using both Spearman’s rank-order correlation and Kendall’s tau rank correlation. The overall Tex and NK1 values for a given cancer type were defined by their respective median values. GAM was deployed to account for observed non-linear trends. Subgroup and sensitivity analyses were performed based on clinical variables (cancer type, stage, organ system, and anatomical site) as well as diverse technical subgroups (sequencing platform, cell-sorting strategy, and normalization method). Cancer types were classified into four groups based on killer divergence and hot-cold status, which was defined as the combined CD8^+^ T and NK cell fraction relative to total CD45^+^ cells. These classifications were associated with cancer-level ORR. A multivariate linear model was constructed to predict Tex or NK1 levels based on cancer type, stage, site, and key molecular subgroups for NSCLC, breast invasive carcinoma, head and neck squamous cell carcinoma, hepatocellular carcinoma, glioblastoma, and ovarian cancer.

### Malignant cell analysis

#### 1) Identification of malignant cells

To distinguish malignant populations from normal epithelial cells, CNVs were inferred on a per-sample basis utilizing the inferCNV (30) and CopyKAT (31) algorithms. For the analysis of meta-programs (MPs) and CytoTRACE2 (29), we tested two different sets of tumor cells: 1) only those identified by inferCNV, and 2) the overlapping consensus of both inferCNV (30) and CopyKAT (31).

The inferCNV was executed utilizing a sliding window length of 101 genes. A minimum expression cutoff was tailored to the specific sequencing technology, utilizing a threshold of 1.0 for Smart-seq2 datasets and 0.1 for droplet-based technology such as 10x Genomics. Technical noise was denoised with an SD amplifier of 1.5. Subcluster-level HMM predictions were activated (HMM = TRUE, analysis_mode = “subclusters”, per_chr_hmm_subclusters = TRUE), utilizing a Bayesian latent mixture model to compute the posterior probabilities of alteration (BayesMaxPNormal = 0.5). The tumor subcluster partition algorithm was set to “leiden” and, as recommended by the developers, the initial resolution was lowered as the cellular yield increased. To ensure optimal topological granularity before executing the resource-intensive HMM, preliminary runs were initially restricted to step 15 (up_to_step = 15); the resultant subclustering heatmaps (infercnv_subclusters.png) were visually inspected, and the Leiden resolutions were empirically fine-tuned where necessary.

The final inferCNV-defined malignant cells were established by the intersection of the two methodologies: categorical HMMi6 state distributions and continuous CNV scoring. The former can determine the proportion of chromosomal regions assigned to the neutral, unamplified diploid state (State 3). Subclusters having < 30% of their regions in the State 3 were flagged as malignant. Additionally, a quantitative CNV score was calculated independently for every cell. A neutral baseline was established utilizing the reference cell population, defined as the mean expression bounded by ± 2 standard deviations (SD). Absolute deviations outside this neutral interval were converted into a point-based penalty system: complete single-copy alterations (loss or addition) were assigned 1 point, while multi-copy amplifications or complete homozygous deletions were assigned 2 points. These values were summed to generate a total CNV score per cell. Subclusters with an average CNV score exceeding the normal reference threshold (defined as the reference mean + 1 SD) were classified as malignant.

For CopyKAT, the minimum number of genes required per chromosome segment was set to 4 (ngene.chr = 4), and the Kolmogorov-Smirnov test P value cutoff for segmentation was set to 0.05 (KS.cut = 0.05). The analysis was restricted to genes expressed in a specified proportion of cells, defined by a lower limit of 3% and an upper limit of 10% (LOW.DR = 0.03, UP.DR = 0.1). Genomic segmentation was computed across a window size of 25 genes (win.size = 25). Cells classified by as “aneuploid” were designated as malignant.

#### 2) Meta-program analysis

Hotspot algorithm (78) was performed to detect functional co-expression modules of tumor cells independently for each sample. The optimal number of PCs was determined iteratively to select the minimal number of PCs required to cumulatively explain 75% of the total variance. The Hotspot object was modeled via a depth-adjusted negative binomial distribution. A K-nearest neighbors (KNN) graph was constructed to map the local neighborhoods of the tumor cells. To balance the detection of robust global correlations against fine-grained local patterns, the KNN neighborhood size was scaled to the tumor cell yield. Genes demonstrating significant autocorrelated expression across the KNN graph were retained. Samples failing to yield a minimum of 30 significantly autocorrelated genes were excluded from downstream module detection. Pairwise local correlations were calculated to quantify the degree to which informative genes shared identical local expression topologies. Module branching was controlled using an FDR threshold of 0.05.

For each sample, unassigned genes were discarded, and the remaining genes within each module were ranked in descending order according to their Z-scores. Each module was capped at a maximum of 50 top-ranked genes. To transition from sample-level modules to intermediate MPs (iMPs) that recurrently occur across tumors, every module was compared against all other modules across the cohort. An iMP was successfully nucleated only if a core module shared a minimum overlap of 10 genes with at least 10 other independent sample modules. Within these qualifying iMPs, the constituent genes were aggregated and ranked—first by their total cross-sample appearance frequency, and subsequently by their median Z-score—to distill each iMP network down to its 50 most definitive genes. To derive a set of non-redundant MPs, a pairwise Jaccard similarity matrix was computed across all iMPs. To objectively determine the optimal number of distinct MPs (k), the dendrogram was iteratively partitioned across a testing range of 5 to 20 clusters. For each k, the maximum gene overlap between any two resultant clusters was quantified. The optimal clustering number was defined as the highest k that maintained independence, ensuring that no two MPs shared more than 20% of their top genes. This identified k = 9 as the ideal resolution (Fig. S23). To establish the signature for each final MP, all genes from its constituent iMPs were ranked first by their total appearance frequency within the cluster, and subsequently by their median Z-score.

To assign biological functions to MPs, gene set enrichment analysis was performed utilizing the compareCluster algorithm within the clusterProfiler R package (79) based on a previously established meta-program resource (27) and MSigDB signatures (https://www.gsea-msigdb.org/gsea/msigdb). To ensure that only highly cohesive and reproducible meta-programs were retained for final downstream analysis, each MP cluster was evaluated utilizing two metrics: (1) The average Jaccard index measuring internal core cohesion. (2) Silhouette coefficient evaluating how tightly grouped the modules were within their assigned MP relative to their separation from other MPs. Any MP demonstrating an average intra-cluster Jaccard similarity < 0.3 or a mean silhouette score < 0.3 was indicative of a diffuse gene network and was excluded. This process removed one MP, and a suspected doublet MP was subsequently excluded due to high expression of T cell lineage genes, yielding a final set of seven MPs.

We utilized the approach described by Tyler et al. (27) to quantify the enrichment of the MPs on the pseudobulk matrices. First, all genes within the dataset were ranked and stratified into 200 expression bins based on their global average expression. For every target gene within a specific MP signature, 100 background control genes were randomly sampled from its exact corresponding expression bin. The background-adjusted expression value for each target gene was calculated by subtracting the mean expression of its 100 matched control genes from its expression value. The final MP score for each biological sample was defined as the mean of these background-adjusted values across all genes within the respective MP. MP scores were compared across three patient-matched epithelial states using a LMM: (1) normal epithelial cells derived from adjacent non-malignant tissue; (2) intratumoral normal epithelial cells; and (3) intratumoral malignant cells. Pairwise statistical contrasts between the three epithelial states were computed using estimated marginal means. Comparisons between Tex-skewed (Ts) and NK1-skewed (Ns) tumors within identical tissue compartments were evaluated utilizing the Wilcoxon rank-sum test.

#### 3) Other analyses

We used CytoTRACE 2 (29) to evaluate differentiation potential, and the undifferentiated proportion was calculated. To test whether tumors with low epithelial MHC-I expression might be enriched for NK1 rather than Tex, we calculated an MHC score, defined by the median expression of classical HLA-A, HLA-B, and HLA-C genes, and assessed the correlation of MHC-I score with Tex or NK1.

### Killer divergence dominates global variation

The methodology for understanding the major variation axis across the entire TME was analogous to the PCA-based approach used to evaluate variation within the killer compartment (see section “Uncover killer divergence in lung cancer”). However, the input features were expanded to broader cellular composition metrics from all TME cell lineages alongside the tumor MP scores (Tables. S5 and S6). Tissue-specific cell types, including oligodendrocytes and microglia, that do not exhibit broad cross-sample conservation were excluded from the PCA. Differential cellular proportion analyses were conducted between the Ts and Ns groups. The differential effect size for each TME feature was quantified as the difference between the 75th percentile values of the Ns and Ts groups. To ascertain whether the Tex-NK1 divergence anchors the primary variance of the global TME, the top 30 features driving the highest absolute loadings on PC1 were extracted. Spearman correlation was calculated between the PC1 loading weights of these features and their corresponding differential effect sizes. This analytical pipeline was repeated using cell subsets across different layers of granularity (Table. S6).

### Immunotherapy efficacy analysis

#### 1) scRNA-seq cohorts

The predictive value of Tex and NK1 was validated utilizing both cellular fractions and signature scores. Fractions were calculated relative to their respective parent lineages: Tex divided by total CD8⁺ T cells, and NK1 divided by total NK cells. To benchmark the predictive performance of the Tex-NK1 axis against other cell subsets, we curated signatures for several cell subsets previously established as biomarkers for ICB efficacy in high-quality single-cell studies across diverse lineages (Table. S2). The gene list for each cell subset was established by jointly considering the top-ranking marker genes provided in the original publications and the DEGs identified in our cohort. For cell subsets reported across multiple independent single-cell studies, a robust consensus gene signature was derived by extracting the intersection of the published gene sets. We recorded the predictive directionality (i.e., a positive or negative association with ICB efficacy) as demonstrated in the original publications. If a specific cell subset had been evaluated using both bulk RNA-seq (e.g., via deconvolution or signature scoring) and scRNA-seq, the predictive directionality was based on the latter. We evaluated the association of each biomarker with ICB response utilizing a Wilcoxon rank-sum test.

The action of ICB is driven by the expansion of progenitor memory CD8⁺ T cells. Consequently, assessing the dynamic shifts of these specific T cell subsets following ICB administration serves as a biological endpoint to evaluate differential treatment sensitivity between Ts and Ns patients. We curated several relevant signatures (Table. S2) and analyzed matched pre- and on-treatment tumor samples to determine whether their dynamic shifts correlated with clinical response and baseline killer profiles. We performed this analysis specifically within tumor-reactive CD8⁺ T cells because the expression profiles of tumor-reactive populations have been shown to predict ICB outcomes better than analyses based on total CD8⁺ T cells in previous studies. Additionally, key memory genes (such as GZMK, IL7R, and TCF7) also mark bystander CD8⁺ T cells beyond the treatment-related populations. Evaluating these shifts within the total CD8⁺ T cell compartment would therefore conflate these distinct cellular sources. Given the lack of TCR data for a substantial proportion of samples, we designed a transcriptomics-based method to differentiate tumor-reactive CD8⁺ T cells in silico. Based on a previous meta-analysis (25), the single gene CXCL13 performs best in identifying tumor-reactive T cells in both pre- and post-treatment samples, outperforming the exhaustion signatures. Therefore, tumor-reactive subpopulations were isolated utilizing the AUCell algorithm (77) based on the CXCL13 gene. Subsequently, pseudobulk expression matrices were established for scoring. This analysis was repeated stratified by early and late on-treatment timepoints based on a cutoff of four weeks post-initiation of ICB therapy. Significance of these shifts were evaluated using paired Wilcoxon signed-rank tests.

#### 2) Bulk RNA-seq cohorts

We assessed whether the pseudobulk profiles of the two killer-skewed groups were associated with ICB efficacy signatures derived from large-scale bulk RNA-seq ICB datasets (Table. S1). To ensure the pseudobulk mirrored bulk tissue architecture, any samples subjected to flow sorting were excluded. Additionally, only samples simultaneously exhibiting detectable proportions (fractional abundance > 0 and < 0.95) across all major TME compartments—including epithelial cells, T/NK cells, myeloid populations, B cells, and stromal elements—were deemed valid and retained for pseudobulk aggregation.

To define a robust set of genes that predict ICB efficacy, each gene was analyzed individually within the ICB arm of each study, after which the resulting effect sizes and standard errors were pooled utilizing a random-effects meta-analysis model via the MetaIntegrator R package (80). Only genes available in at least 30 cohorts were considered for analysis. Gene expression was evaluated against three clinical endpoints: response, DCE, and OS. For response, differential expression effect sizes were calculated between responders and non-responders within each independent study. For time-to-event endpoints, patient were stratified into the upper quartile (> 75th percentile) versus the lower quartile (< 25th percentile). Genes exhibiting significance across all three endpoints were used to build the ICB efficacy signature. To ensure that the signature demonstrated high cross-modality expression concordance, the median gene expression profiles of the pseudobulk lung discovery cohort were mapped against the actual bulk RNA-seq expression levels of the TCGA-NSCLC cohort. The overall correlation between the pseudobulk and real-bulk datasets was high (Spearman’s rho = 0.81, Fig. S36). Genes falling within the bottom 10% of global expression in either the pseudobulk or real-bulk datasets were discarded. The absolute difference in median expression between the two modalities was calculated. The signature was restricted to genes falling within the lowest quartile (bottom 25%) of this absolute difference. These efficacy-associated genes were inputted into the STRING database (https://string-db.org/) for protein–protein interaction testing and enrichment analysis. Gene set enrichment analysis was performed to assess the differential enrichment of this signature between the two killer-skewed groups using the clusterProfiler R package (79).

We also assessed whether the relative efficacy of ICB over conventional therapy differed between the Ts and Ns groups. To derive a predictive signature capable of distinguishing true ICB-specific benefit from general prognostic advantages, a meta-analysis of treatment-by-gene interaction tests was conducted. For each RCT, the predictive capacity of every gene was evaluated independently across three types of clinical endpoints as above. For time-to-event endpoints, a Cox Proportional Hazards regression model was utilized. The interaction Z-score was extracted from the model summary. To align the mathematical directionality with clinical benefit, the sign of the survival Z-score was inverted so that a positive value indicated a lower hazard ratio (improved survival) in the ICB arm relative to the control arm. For binary response data, a logistic regression model was employed. The interaction Z-score was extracted, where a positive value intrinsically indicated higher odds of clinical response under ICB relative to conventional therapy. To determine if a gene provided a consistent ICB-specific benefit across eight RCTs, the vector of pooled Z-scores for each gene was evaluated utilizing a Wilcoxon signed-rank test against a theoretical median of zero. Genes were considered significant if they met the following endpoint-specific thresholds, established to account for the varying number of RCTs available for each endpoint: P < 0.05 for response, P < 0.05 for DCE, and P < 0.1 for OS. Genes that achieved significance concurrently across all three types of endpoints were intersected to form a preliminary signature, which was also filtered against the cross-modality high-concordance gene list, ultimately identifying GBP1 and FCRLA. An ICB-over-control score was computed for each sample based on the mean expression of the two genes, and these scores were compared between the Ts and Ns groups utilizing a Wilcoxon rank-sum test.

Of note, we did not employ deconvolution strategies such as BayesPrism here, because Tex and NK1 subsets are characterized by a fine resolution, low fractional abundances, and highly similar transcriptional profiles. Under these conditions, deconvolution frequently yields low-confidence estimates due to insufficient distinct signals within the bulk mixture. This uncertainty is quantitatively reflected by elevated theta.cv values in BayesPrism (data not shown).

#### 3) Checkpoint comparison

ICB targets inhibitory checkpoints, thus the differential expression of these targets can be utilized to assess ICB sensitivity. A curated panel of T/NK checkpoints (KIR3DL2, KIR3DL1, KIR2DL3, KLRG1, LAG3, CD300A, TIGIT, KIR2DL1, SIGLEC7, SIGLEC9, LILRB1, LAIR1, KLRC1, PDCD1, CTLA4, HAVCR2, BTLA, ADORA2A, ADORA2B, VSIR, CD276, and CD47) was evaluated to compare NK1 and Tex within the lung discovery scRNA-seq cohort. Additionally, PD-1 protein expression was compared between the NK1 and Tex via spectral cytometry, utilizing a paired Wilcoxon signed-rank test.

### Prioritization of targets

#### 1) Ligand-receptor (LR) interaction analysis

We aimed to identify dominant LR interactions mediating NK1 immune evasion specifically rather than mere byproducts of NK differentiation. To mitigate the risk of false negatives driven by dropout, a candidate receptor was required to be expressed in at least 5% of the NK1 cells. Concurrently, any candidate interacting ligand was required to meet this 5% expression threshold within its respective putative source cell lineage. We evaluated LR pairs through a three-tier pseudobulk-based differential enrichment pipeline:

- **Receptor upregulation:** The receptor must exhibit significantly higher expression within the NK1 cells of Ns tumors compared to Ts tumors.
- **Ligand availability:** The corresponding ligand must demonstrate significantly higher expression across the global, unsorted pseudobulk profile in Ns tumors.
- **Source explainability:** To confirm that the global ligand upregulation is not a bulk artifact, the specific cellular origin of the ligand must be identified. The ligand must be significantly upregulated within the isolated pseudobulk profile of at least one specific cell type. Ideally, these source cellular subsets should also align with the Ns direction along the PC1 axis.

To obtain a comprehensive LR interaction database, four databases (LIANA [81], CellChatDB [82], CellTalkDB [83], and CellPhoneDB [84]) were integrated and deduplicated. We also manually added LR pairs derived from recent publications that were omitted from these established databases. The final compiled human interaction database contains 7,369 unique interactions.

#### 2) Cytokine analysis

Cytokine activity was estimated from the pseudobulk transcriptomic profiles utilizing the CytoSig algorithm (50). We explored whether intracellular checkpoints might attenuate cytokine signaling based on a panel of IL-15 signaling negative regulators identified via genome-wide CRISPR screening by Nikolic et al. (47). The expression of these genes were evaluated within NK1 cells between Ns and Ts tumors.

#### 3) EvoTarget

EvoTarget was designed as an scRNA-seq-based, unbiased screening strategy to prioritize tumor-cell genes associated with the NK1-skewed immune-evasion axis. A gene was considered expressed in a tumor cell if its raw count was greater than zero, and NK1-directional candidate genes were retained for subsequent scoring only if their mean sample-level expression proportion in Ns was at least 5% in the lung discovery cohort, thereby removing lowly expressed genes while minimizing false-negative exclusion caused by scRNA-seq dropout. Retained genes were then evaluated using five independent evidence layers, with all analyses performed at the sample or cancer-type level to avoid cell-level pseudoreplication. First, tumor-cell differential-expression evidence prioritized genes with stronger expression enrichment in Ns over Ts in the lung discovery cohort. Second, sample-level correlation analyses assessed whether tumor-cell expression was positively associated with the NK1 score and negatively associated with the Tex score across independent cohorts (lung discovery, pan-cancer, and in-house scRNA-seq cohorts) and available clinical or technical strata with more than 15 samples. Third, cancer-level correlation analysis summarized tumor-cell gene expression, NK1 score, and Tex score by cancer-type medians and prioritized genes showing concordant positive correlations with NK1 and negative correlations with Tex across tumor types with more than 15 samples. Fourth, pretreatment malignant tumor-cell pseudobulk profiles from the ICB scRNA-seq cohort were used to prioritize genes with higher expression in non-responders than in responders. Finally, scores from these layers were summed into an NK1-directional EvoTarget score.

### Statistical analysis

For multiple comparisons, P values were adjusted to control the FDR utilizing the Benjamini-Hochberg procedure. All statistical analyses were executed utilizing R software (version 4.4.0) and Python (version 3.11).

## Data Availability

The newly generated human sequence data, including pan-cancer single-cell RNA-seq data and bulk RNA-seq data from the three in-house clinical trials will be available at publication date. All other public or controlled-access data used in this study are listed in Table S1.

## Acknowledgments

This work was supported by Noncommunicable Chronic Diseases-National Science and Technology Major Project (SDH: 2024ZD0520200 and 2024ZD0520205), Guangdong Basic and Applied Basic Research Foundation (SDH: 2023B1515020008), National Natural Science Foundation of China (SDH: 82573424; LZ: 82241232, 82272789, U25C2027), and National Science and Technology Major Project (LZ: 2024ZD0519700). We are grateful to the patients enrolled in all datasets utilized in this study. We thank Prof. Dawei Yang, Prof. Christopher J. Hanley, Prof. Peng Zhang, Prof. Fengying Wu, Dr. Yulan Deng, Dr. Siyang Li, Dr. Bolun Zhou, Prof. Hirokazu Matsushita, Prof. Weimin Li, Dr. Jiayi Deng, Prof. Wenzhao Zhong, Prof. Hongbin Ji, Dr. Jiatao Zhang, Prof. Satoshi Yamasaki, Prof. Andrew Chow, Prof. Takanari Okamoto, Dr. Zhoufeng Wang, Prof. Jun Wang, Dr. Shijie Tang, Dr. Di Chen, Prof. Hailong Piao, and Dr. Wenxin Luo for sharing controlled datasets and providing detailed clinical information. We also thank Dr. Yongqiang Zheng and Prof. Qi Zhao for their efforts in maintaining the bioinformatics computing cluster at our institution.

## Authors’ Contributions

Anlin Li: Conceptualization, methodology, investigation, visualization, project administration, supervision, writing–original draft, writing–review and editing. Zhixin Yu: Methodology, investigation, visualization, writing–original draft, writing–review and editing. Wenda Zhang: Investigation, visualization. Hanming Chang: Investigation, visualization. Sirui Feng: Investigation, visualization. Xuan Yang: Investigation, visualization. Kangqiao Xiong: Investigation, visualization. Lina He: Investigation, visualization. Zerui Zhao: Investigation. Lujun Shen: Investigation. Zihui Tan: Investigation. Wei Du: Investigation, visualization. Xu Zhang: Investigation. Yi Hu: Investigation. Xiaodong Su: Investigation. Sha Fu: Investigation, funding acquisition, project administration, supervision, writing–review and editing. Li Zhang: Funding acquisition, project administration, supervision, writing–review and editing. Shaodong Hong: Conceptualization, methodology, funding acquisition, project administration, supervision, writing–review and editing.

## Authors’ Disclosures

Authors declare that they have no competing interests.

## Notes

**Conflicts of interest:** Authors declare that they have no competing interests.

### Competing Interest Statement

The authors have declared no competing interest.

