## Supplementary material for "Killer-cell dominance dichotomy governs tumor immune networks and stratifies inflamed cancers": Figure S1-S39

### Supplementary Figures

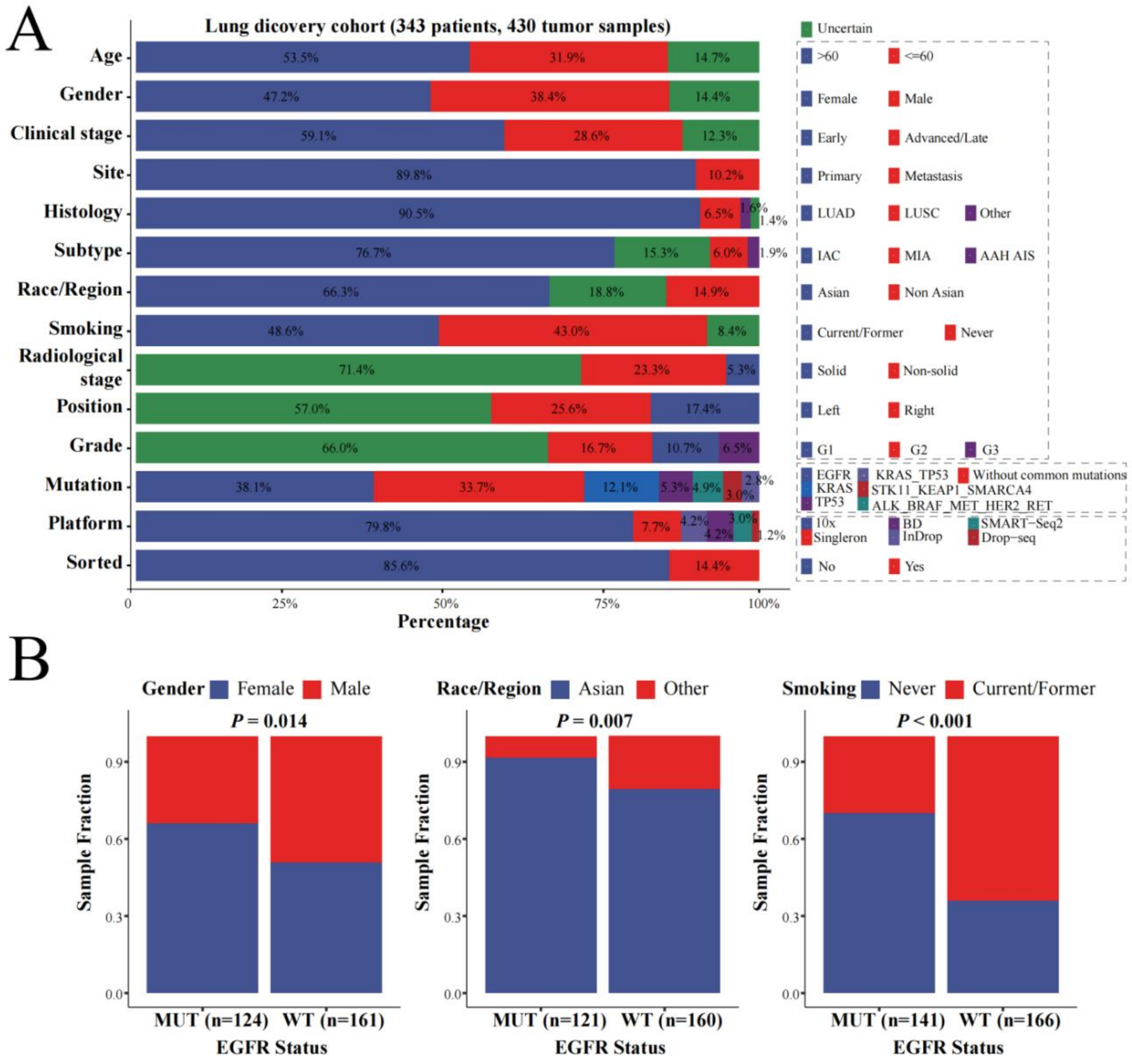


**Fig. S1. Clinicopathologic and molecular information of the lung discovery cohort.** (**A**) Overview of the baseline clinical, pathological, demographic, genetic, and technical characteristics. (**B**) The relative sample fractions of gender, race/region, and smoking history stratified by EGFR mutation status.

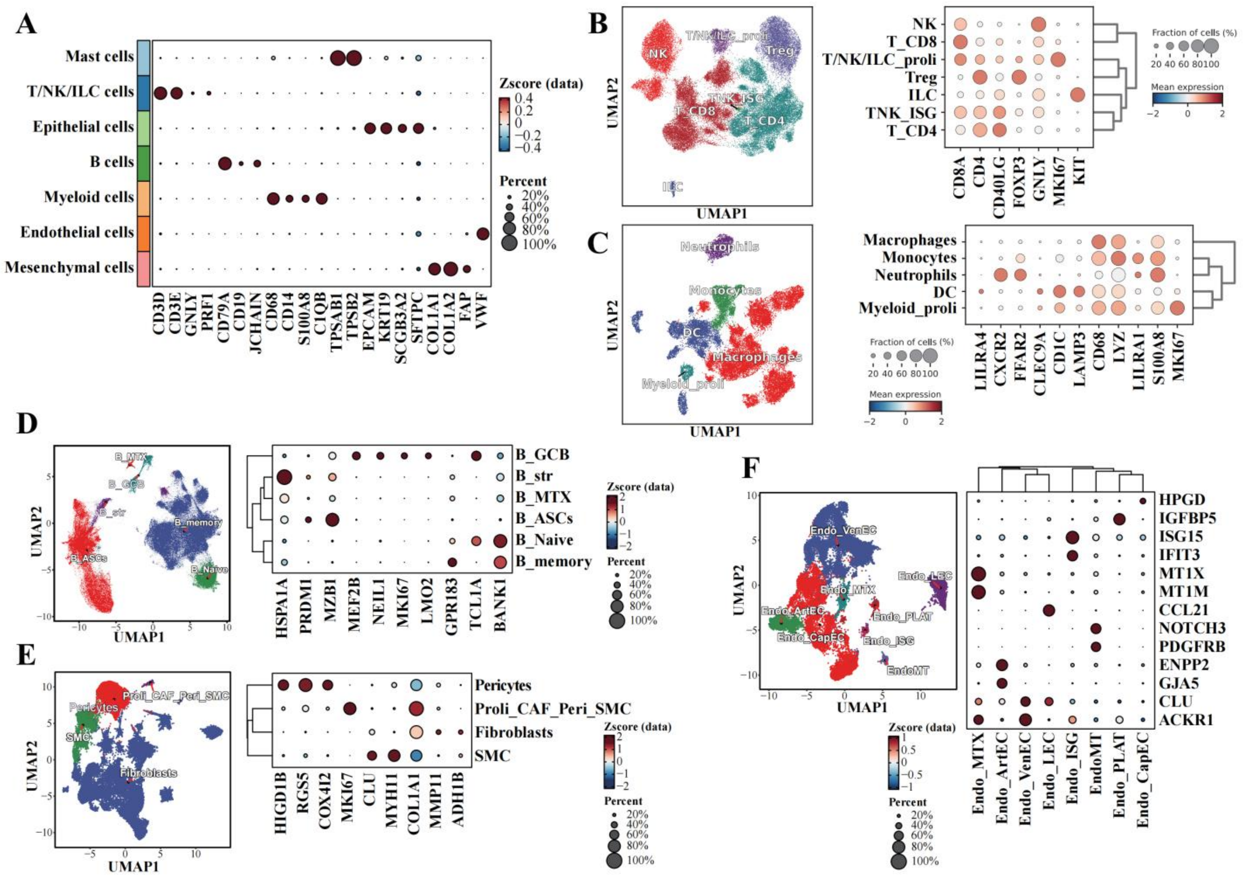


**Fig. S2. Single-cell hierarchical clustering in lung discovery cohort.** (**A**) Dot plot illustrating the expression of canonical marker genes for the annotation of major lineage compartments. (**B to F**) Uniform manifold approximation and projection (UMAP) visualizations (left) and corresponding marker gene dot plots (right) displaying the sub-clustering of specific lineages. To optimize visualization of massive subsets, UMAP embeddings for the T/NK/ILC (**B**) and myeloid (**C**) compartments represent aggregated metacells by SEACells, whereas all other panels represent original single cells. Throughout all sub-clustering dot plots, point sizes indicate the fraction of cells expressing the specific marker, color intensities reflect scaled mean expression levels, and accompanying dendrograms define the hierarchical transcriptional relatedness among the subsets.

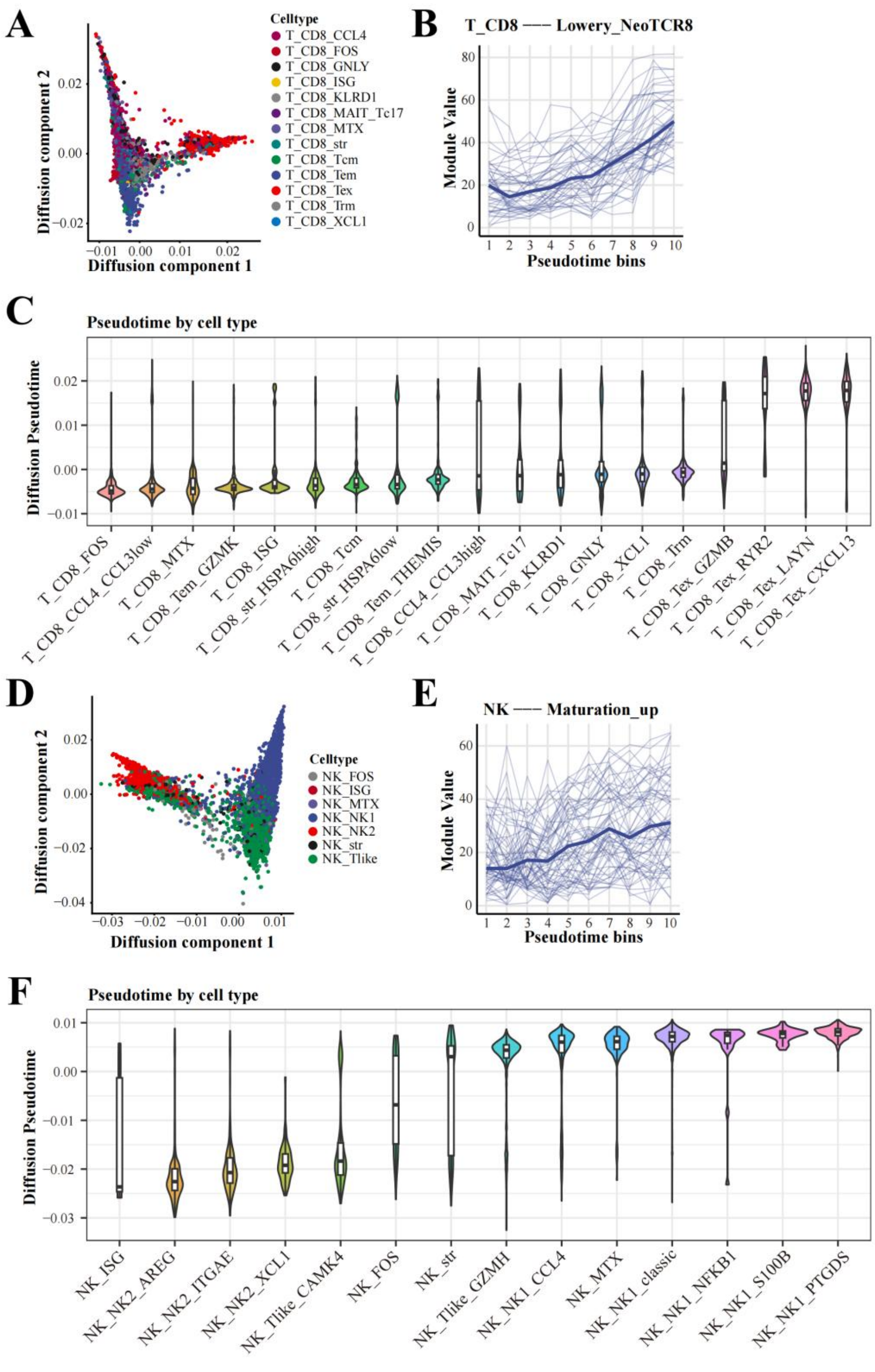


**Fig. S3. Diffusion component analysis of CD8$^+$ T cells and NK cells.** (**A**) Diffusion map embedding of the CD8$^+$ T cells along the first two components (DC1 and DC2). (**B**) Trace-plot of a tumor-reactive T cell score (Lowery_NeoTCR8) along DC1. Cells were stratified into 10 sequential bins based on their DC1 coordinates. Thin faded lines represent the median score values for individual samples, while the bold blue line represents the fitted global median trend across the entire cohort. (**C**) The distribution of DC1 across distinct CD8$^+$ T cell subsets. Subsets are rank-ordered from left to right by their median DC1 . Internal box plots display the median and interquartile ranges. (**D**) Similar as (A) but describing NK cells. (**E**) Similar as (B) but describing the trace-plot of a NK maturation signature along DC1. (**F**) Similar as (C) but describing NK subsets.

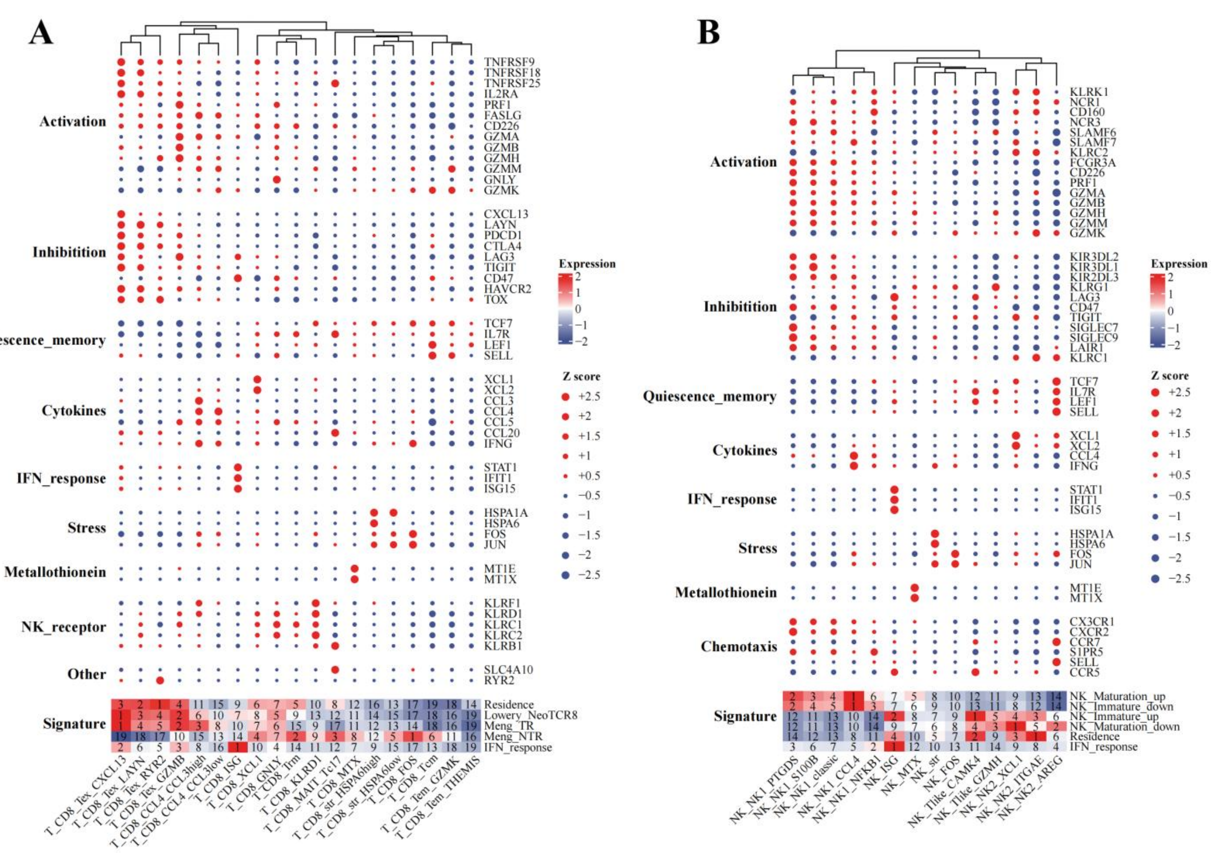


**Fig. S4. Unsupervised hierarchical clustering of markers and gene signatures of CD8⁺ T and NK cells.** Hybrid dot plot and heatmap visualizations displaying the expression profiles of curated functional marker genes and module signatures across (**A**) CD8⁺ T cell subsets and (**B**) NK cell subsets. To mitigate individual sample variance and technical noise, scRNA-seq data were aggregated into sample-level pseudobulk matrices, and the median expression or score values across all samples were utilized to establish subset-specific profiles. The upper sections of each panel present individual marker genes grouped by predefined categories. For individual genes, dot color designates the directionality of the row-scaled Z-score (red indicates positive relative expression; dark blue indicates negative or baseline expression), while dot size is proportional to the absolute magnitude of the Z-score (capped at a maximum Z-score of 2.5 utilizing a non-linear power transformation for visual scaling). The bottom sections display signature scores (color-scaled by Z-score), with the numerical text overlay denoting the intra-signature rank of each subset (where 1 represents the highest scoring subset).

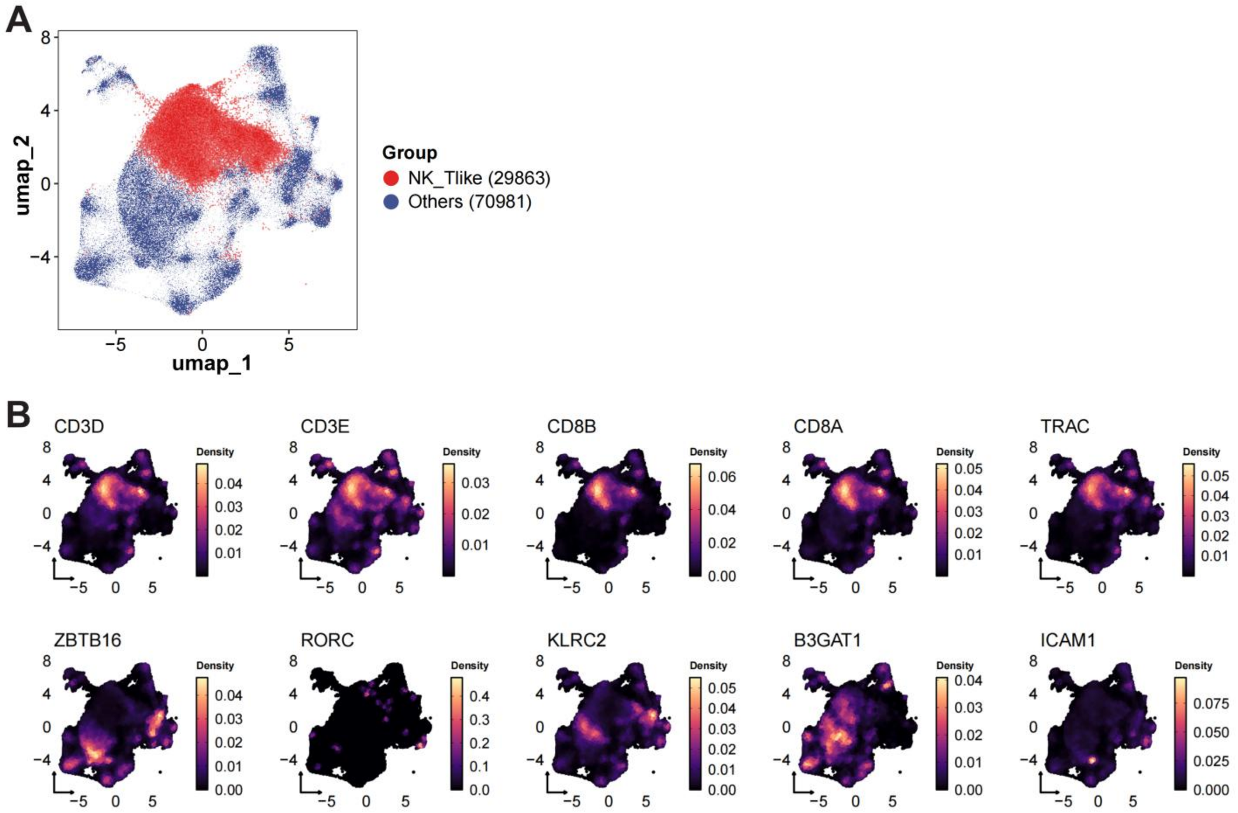


**Fig. S5. Features of the NK_Tlike subset .** (**A**) Uniform manifold approximation and projection (UMAP) embedding of the NK cells, highlighting the position of the NK_Tlike cluster. (**B**) UMAP-based two-dimensional kernel density estimation plots illustrating the expression distribution of selected markers including canonical T cell–defining transcripts (*CD3D*, *CD3E*, *CD8B*, *CD8A*, *TRAC*), natural killer T (NKT) cell transcription factors (*ZBTB16* [PLZF], *RORC*) and memory/adaptive-like NK cell markers (*KLRC2* [NKG2C], *B3GAT1* [CD57]).

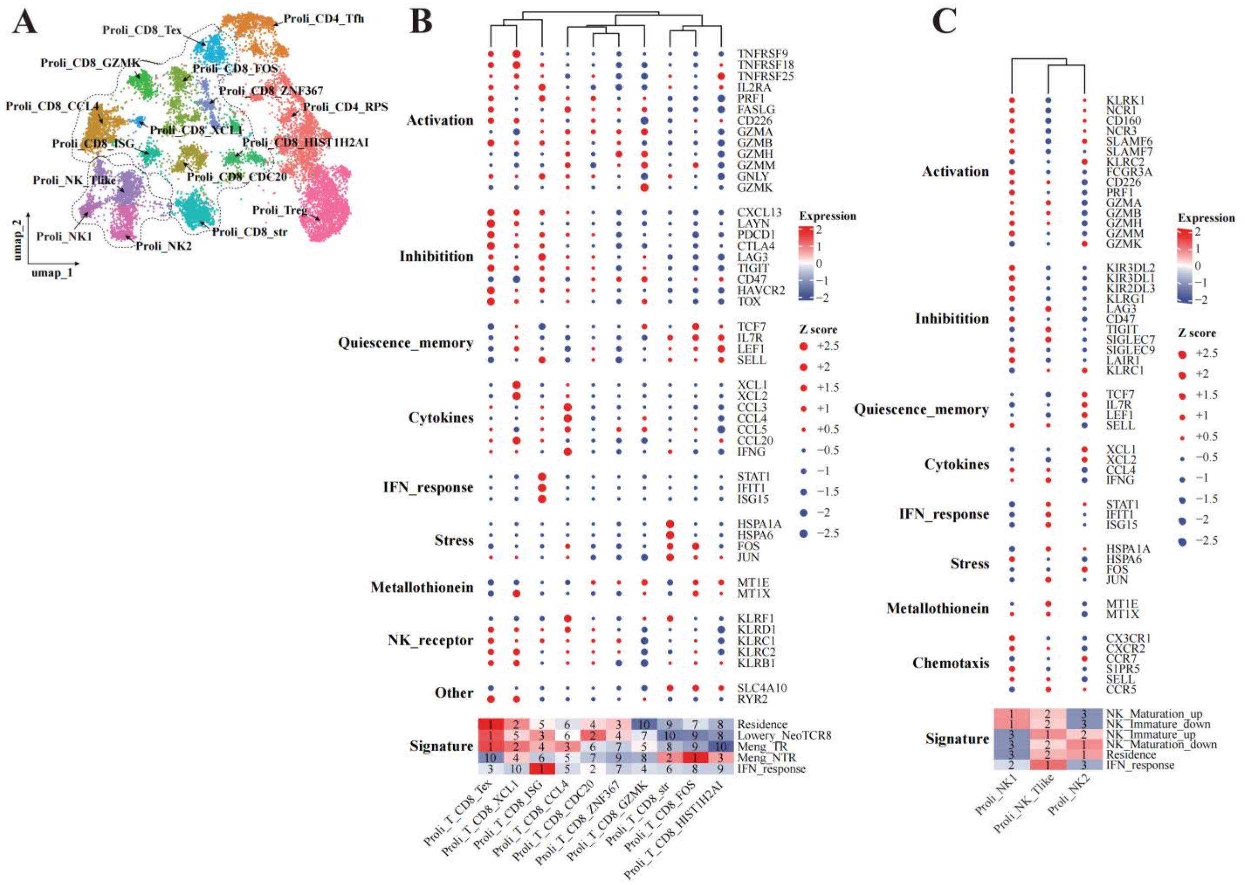


**Fig. S6. Unsupervised hierarchical clustering of markers and gene signatures of proliferating CD8$^{+}$ T and NK cells.** Hybrid dot plot and heatmap visualizations displaying the expression profiles of curated functional marker genes and module signatures across proliferating subsets of (**A**) CD8$^{+}$ T cells and (**B**) NK cells. To mitigate individual sample variance and technical noise, scRNA-seq data were aggregated into sample-level pseudobulk matrices, and the median expression or score values across all samples were utilized to establish subset-specific profiles. The upper sections of each panel present individual marker genes grouped by predefined categories. For individual genes, dot color designates the directionality of the row-scaled Z-score (red indicates positive relative expression; dark blue indicates negative or baseline expression), while dot size is proportional to the absolute magnitude of the Z-score (capped at a maximum Z-score of 2.5 utilizing a non-linear power transformation for visual scaling). The bottom sections display signature scores (color-scaled by Z-score), with the numerical text overlay denoting the intra-signature rank of each subset (where 1 represents the highest scoring subset).

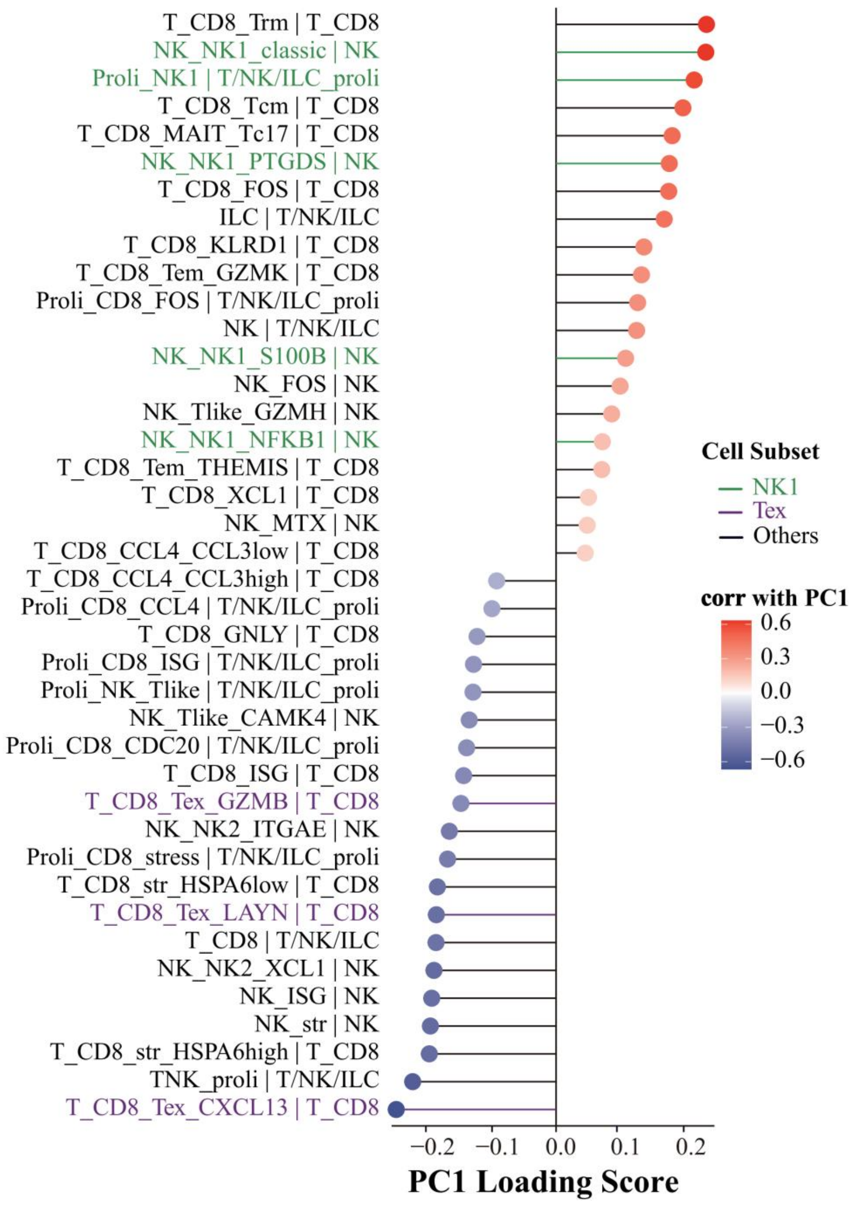


**Fig. S7. PC1 loading scores based on a non-imputation strategy.** PC1 loading scores were derived from a sample-level principal component analysis (PCA) utilizing the proportions of $CD8^+$ T and NK cell subsets. Input data for the PCA underwent a non-imputation, complete-case analysis strategy, wherein any samples lacking quantification for any specific subset were entirely excluded from the matrix.

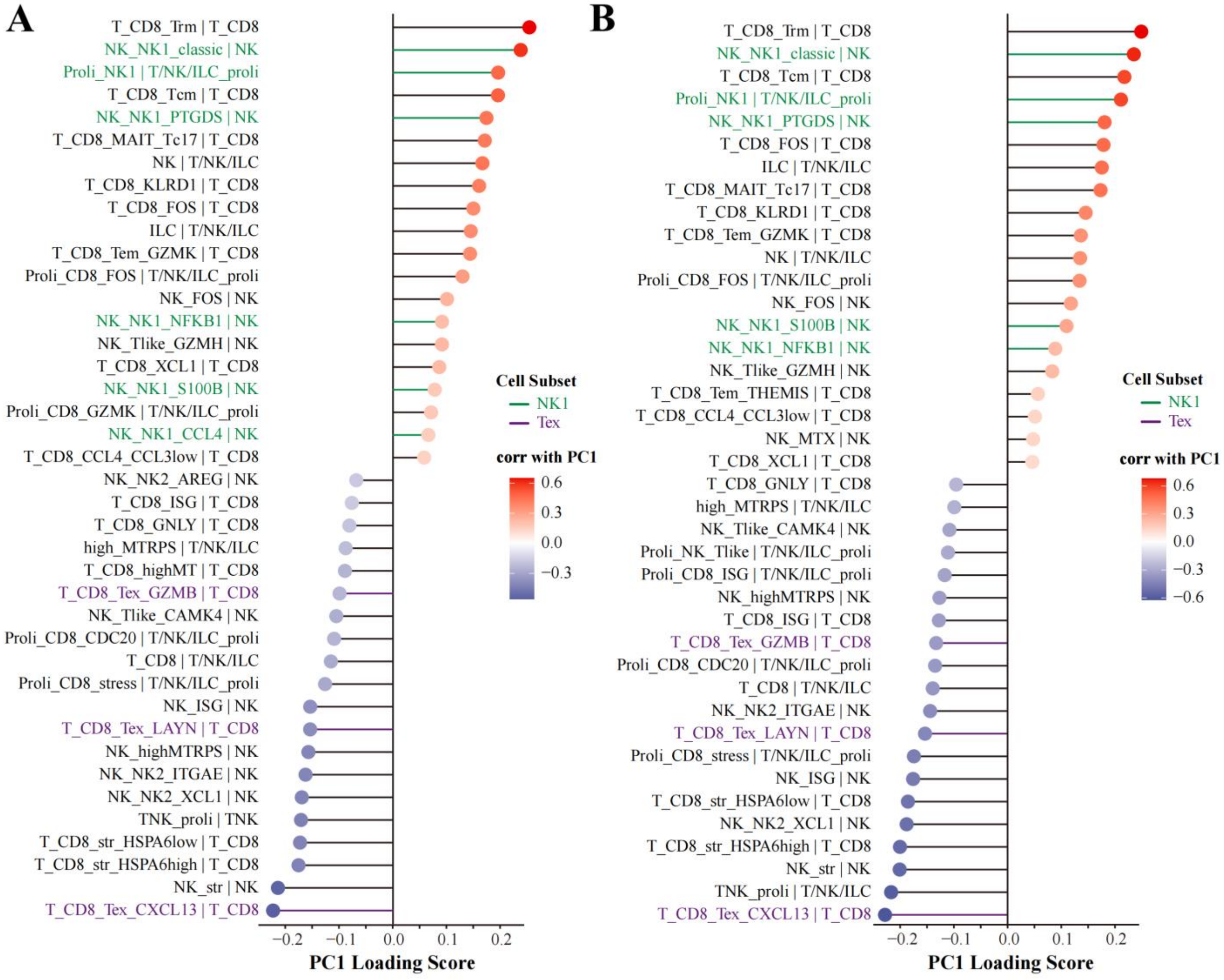


**Fig. S8. PC1 loading scores when retaining clusters with high mitochondrial/ribosomal genes.** PC1 loading scores were derived from a sample-level principal component analysis (PCA) utilizing the proportions of $CD8^+$ T and NK cell subsets. PCA was repeated inclusive of cell clusters exhibiting high mitochondrial or ribosomal gene expression, utilizing two distinct pre-processing strategies for handling missing subset fractions. (**A**) Loading scores following mean imputation, wherein missing values were imputed using the respective column mean. (**B**) Loading scores following non-imputation (complete-case analysis), wherein any samples lacking quantification for any specific subset were entirely excluded from the matrix.

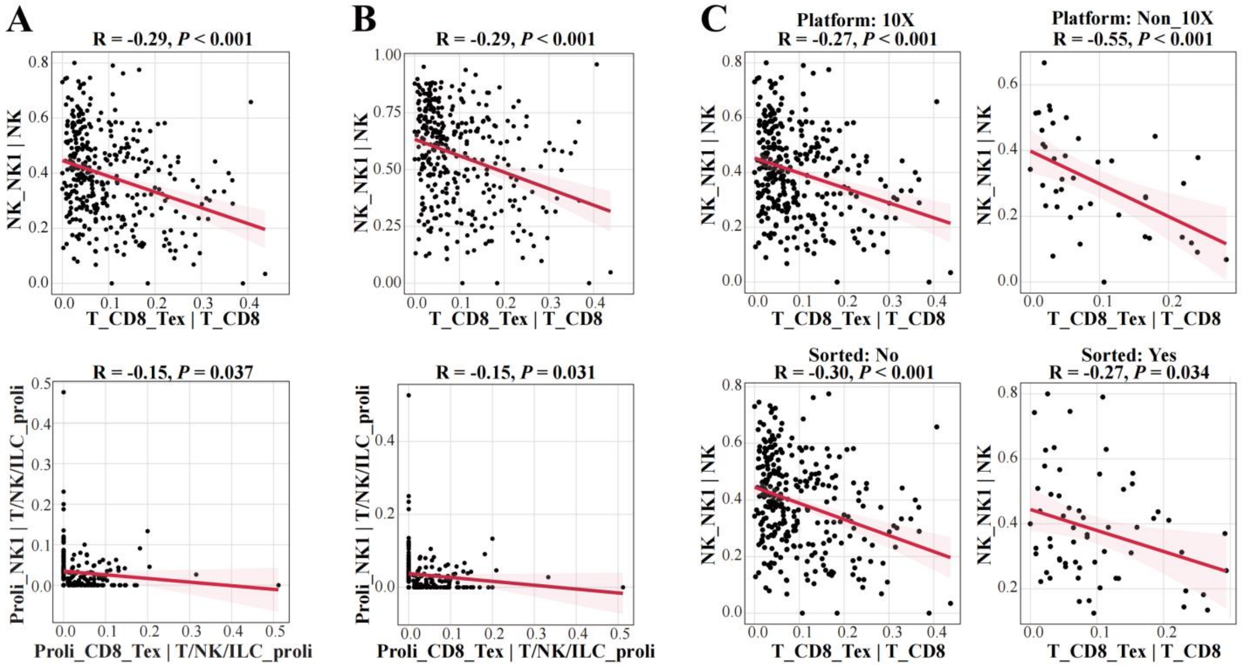


**Fig. S9. Subgroup sensitivity analysis of killer divergence.** (**A**) Inverse correlation between the relative proportions of Tex and NK1, evaluated independently within the global non-proliferating (top) and actively proliferating (bottom) cytotoxic lymphocyte compartments. (**B**) Negative correlation between Tex and NK1 fractions following the exclusion of the ambiguous NK_Tlike subset from the denominator. (**C**) Scatter plots stratified by sequencing platform (10X Chromium versus non-10X, top row) and sample processing protocol (unsorted versus flow-sorted, bottom row). Across all panels, individual dots represent independent patient tumor samples, with the solid red line denoting the linear regression fit and the surrounding light pink shading indicating the 95% confidence interval.

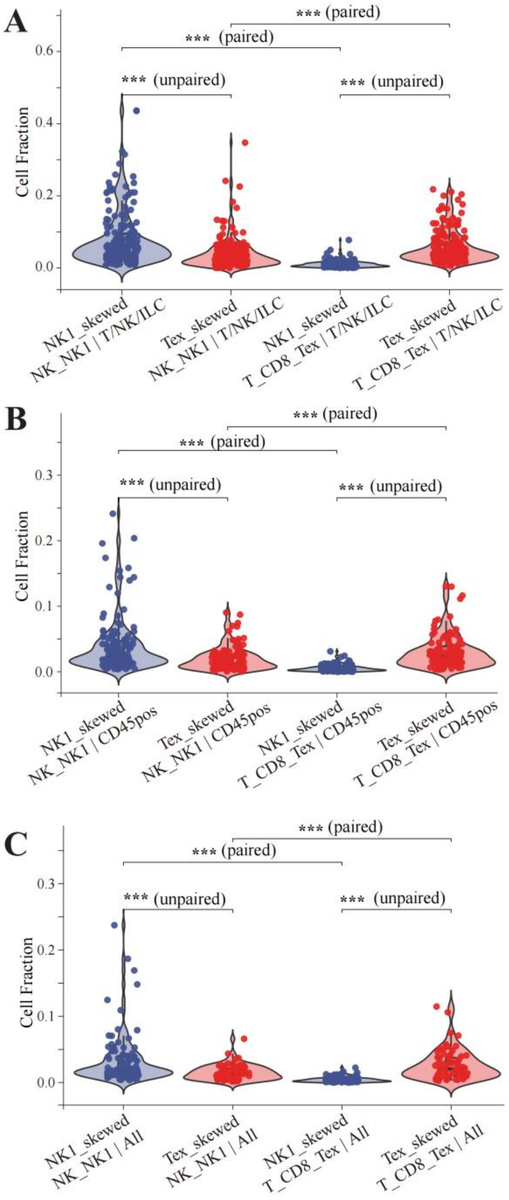


**Fig. S10. Killer divergence persists independent of compositional denominators.** Cellular proportions were systematically recalculated using progressively broader, higher-order denominators: **(A)** the total T, NK, and innate lymphoid cell (T/NK/ILC) compartment, **(B)** the total $CD45^+$ leukocyte (immune) compartment, and **(C)** all captured cells representing the global tumor microenvironment. For analyses utilizing the total $CD45^+$ or global denominators (B and C), samples exhibiting systematic cell-type biases, such as those processed via targeted flow cytometry sorting, were excluded. Statistical significance was evaluated using unpaired Wilcoxon rank-sum tests for cross-group comparisons (comparing the identical cell subset fraction between NK1-skewed and Tex-skewed tumors) and paired Wilcoxon signed-rank tests for within-group comparisons (comparing NK1 versus Tex fractions within the same tumor sample) (****P* < 0.001). Internal box plots delineate the median and interquartile ranges.

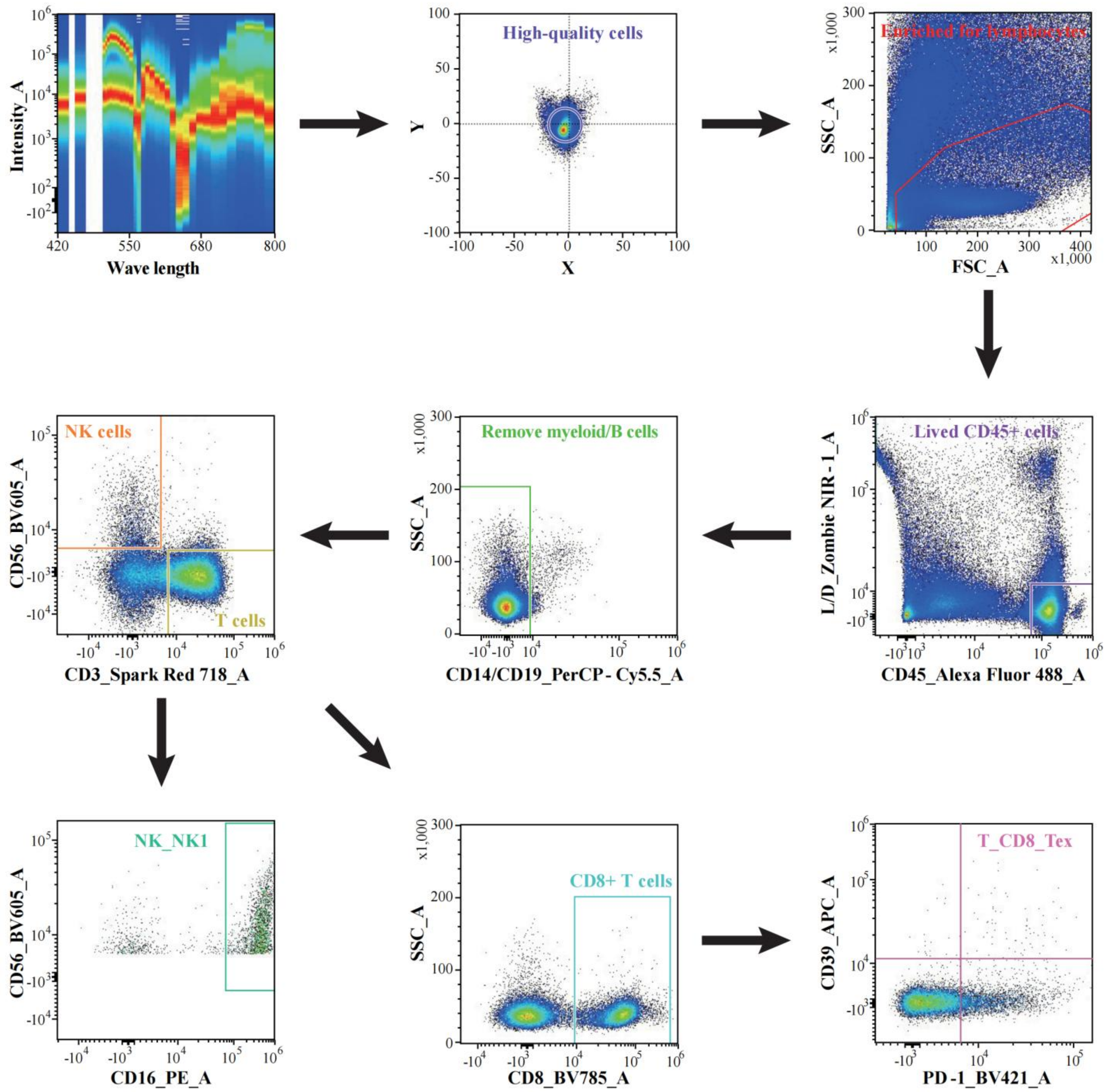


**Fig. S11. The gating strategy in spectral cytometry analysis.**

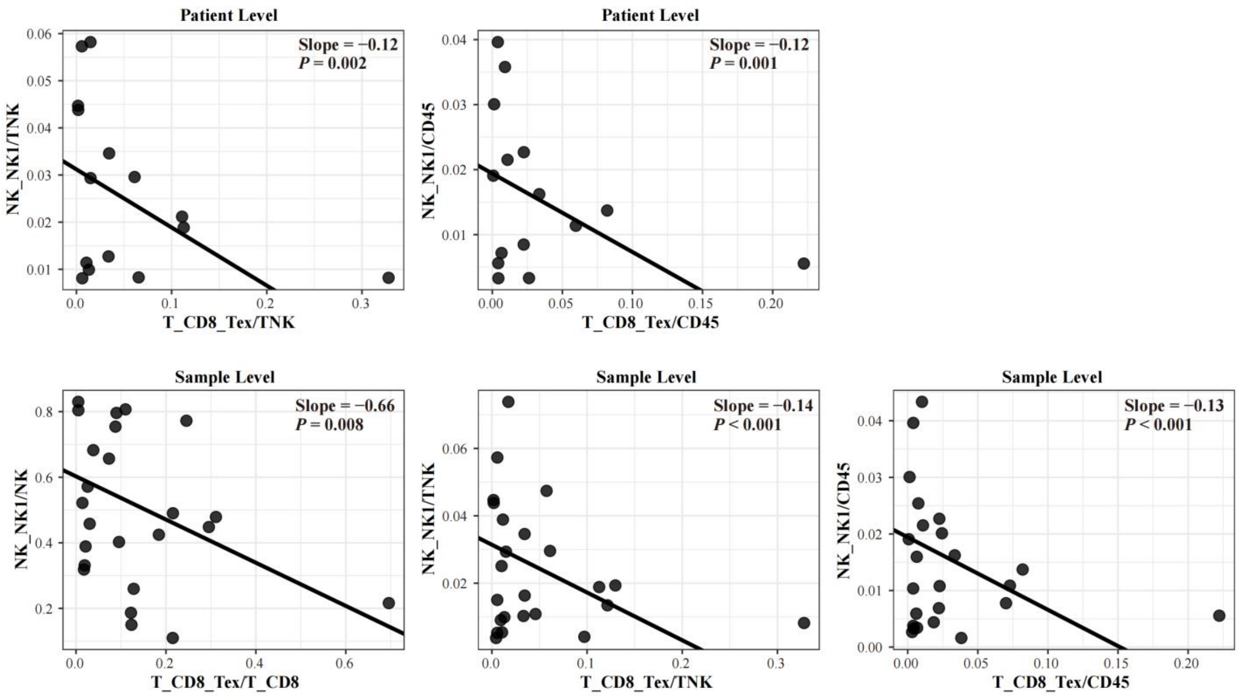


**Fig. S12. Spectral cytometry validation of the killer divergence.** Scatter plot demonstrating a robust inverse correlation between the proportion of NK1 and Tex cells across varied compositional denominators at patient-level or sample-level. The solid black line represents the linear fit derived from a linear mixed-effects model. When evaluating relationships at the sample level, both patient identity and acquisition date were modeled as random intercepts. For patient-level aggregated data, the acquisition date was retained as a random intercept.

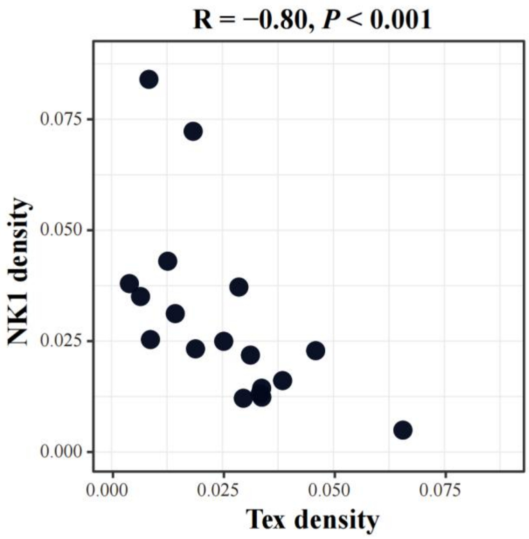


**Fig. S13. Mutually exclusive relationship between Tex and NK1 regarding infiltration density.**

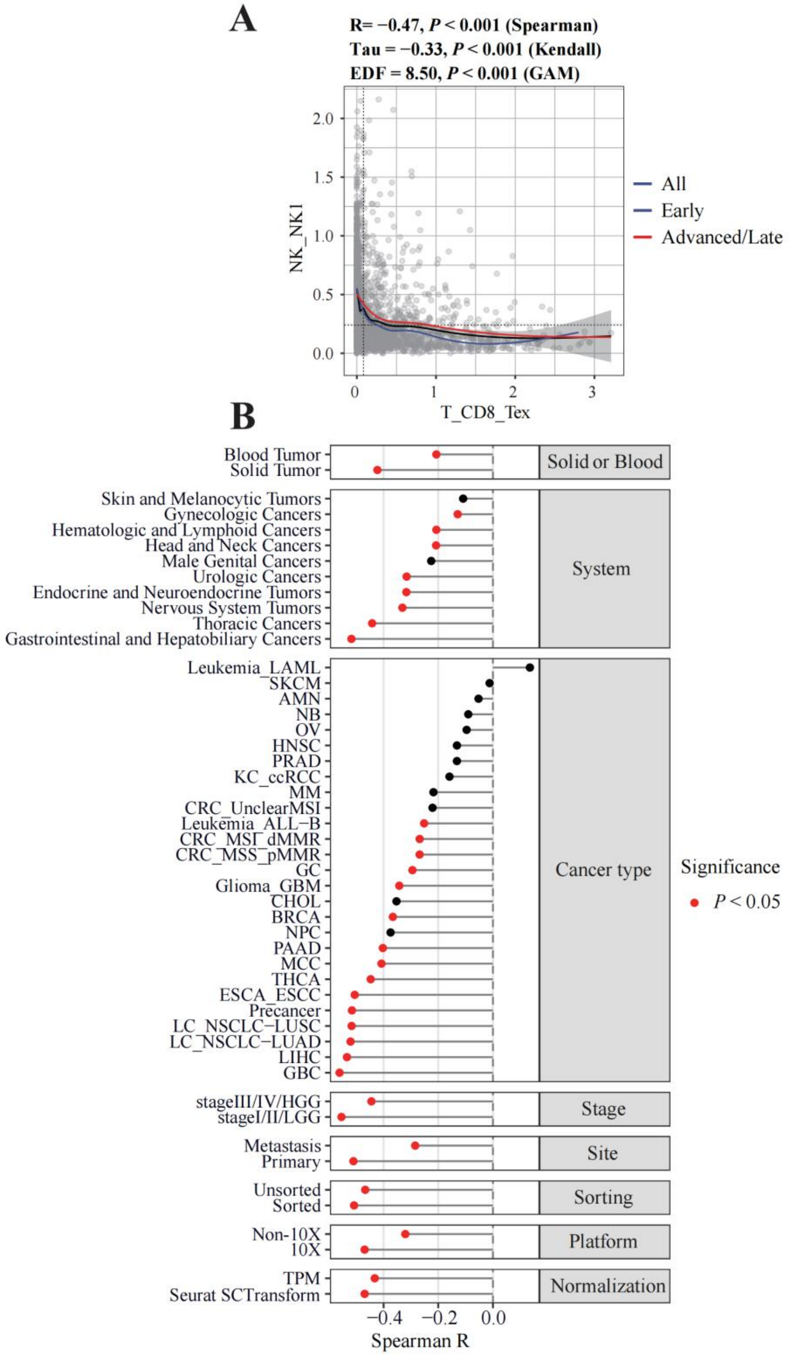


**Fig. S14. Tex-NK1 divergence is a conserved framework in pan-cancer. (A)** Mutually exclusive relationship between the Tex and NK1. The global inverse correlation (blue line) and the correlation stratified by early (dark blue line) versus advanced/late (red line) stage are modeled using generalized additive models. **(B)** Spearman correlation coefficient between Tex and NK1 across subgroups. For both panel A and B, samples from the lung discovery cohort were excluded. Subgroups comprising fewer than 15 samples were excluded from this analysis.Node color indicates statistical significance (red, $P < 0.05$; black, $P \geq 0.05$).

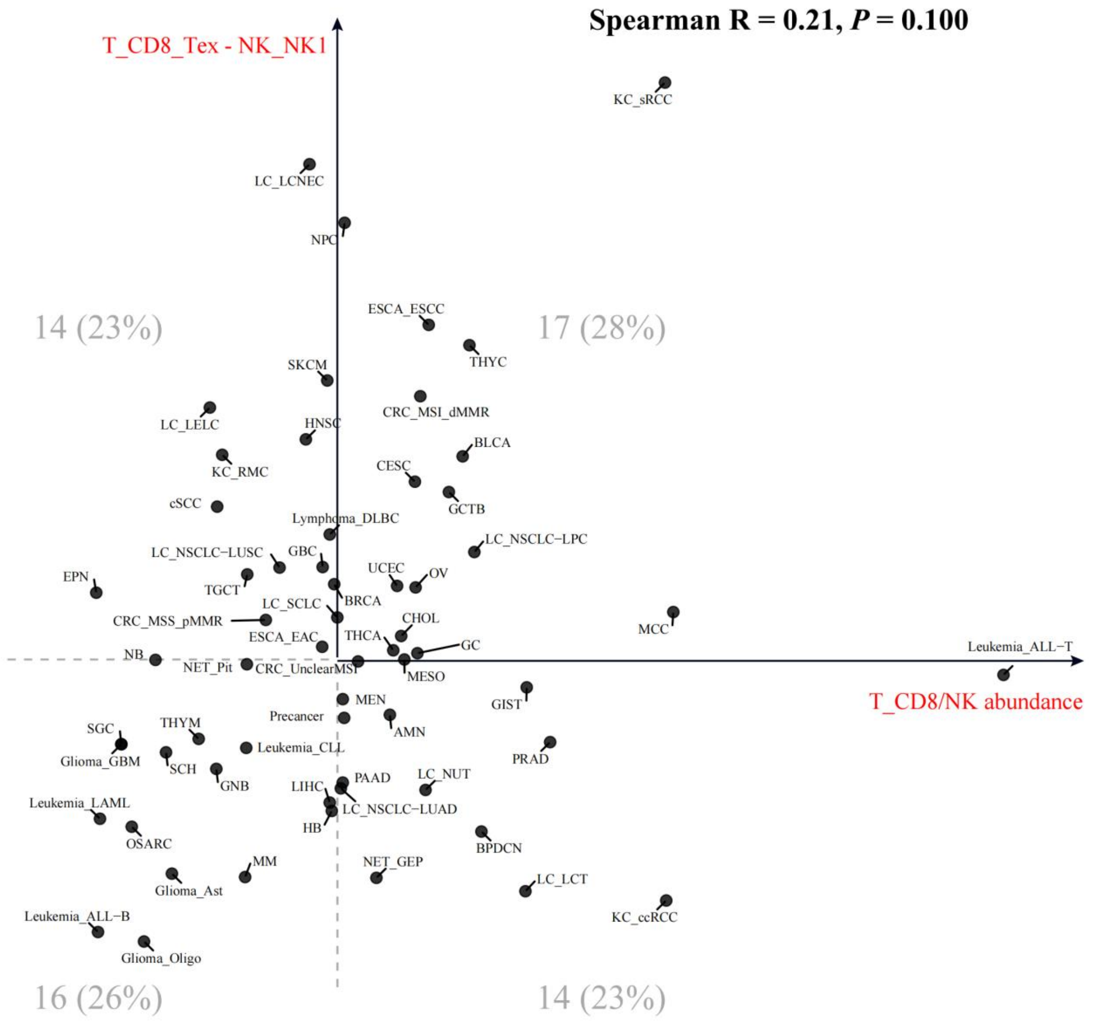


**Fig. S15. Independent relationship between the Tex-NK1 skewness and conventional "hot-cold" system.** The y-axis represents the relative dominance of the Tex versus NK1, computed as the difference in their median score. The x-axis defines the classic "hot-cold" status, quantified by the abundance of the $CD8^{+}$ T and NK cells relative to total $CD45^{+}$ cells. Each data point signifies the median value of a malignancy, with labels corresponding to the abbreviations listed in Table. S1. To center the visualization, both axes were scaled relative to the global pan-cancer medians, dividing the landscape into four combinatorial quadrants.

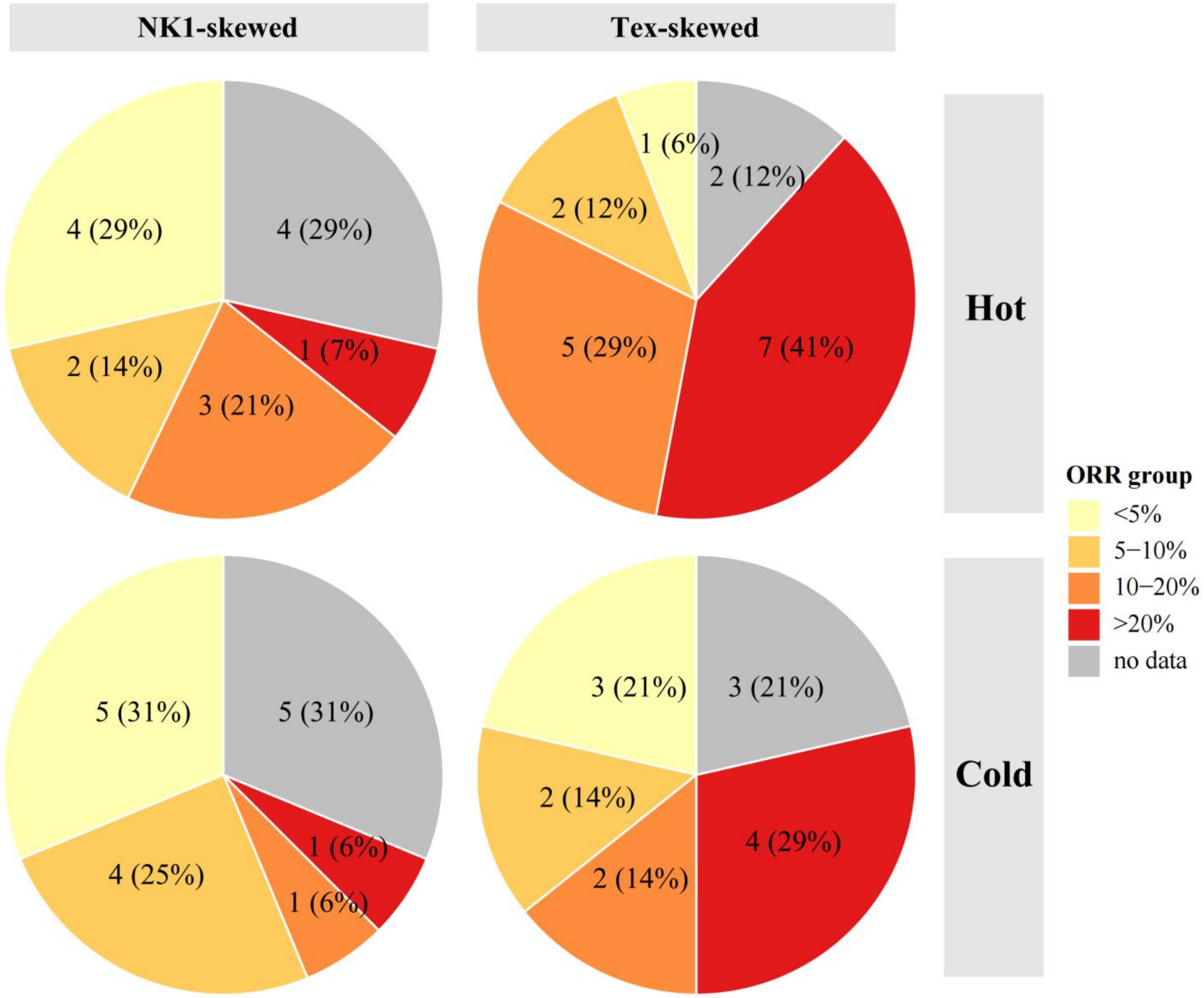


**Fig. S16. Joint relationship of Tex-NK1 skewness and the conventional "hot-cold" system with immunotherapy response.** Distribution of objective response rate groups was stratified into four quadrants defined by the killer divergence axis (columns: NK1-skewed versus Tex-skewed) and the conventional immune infiltration axis (rows: Hot versus Cold). Values within the pie charts represent the number and proportion of cancer types.

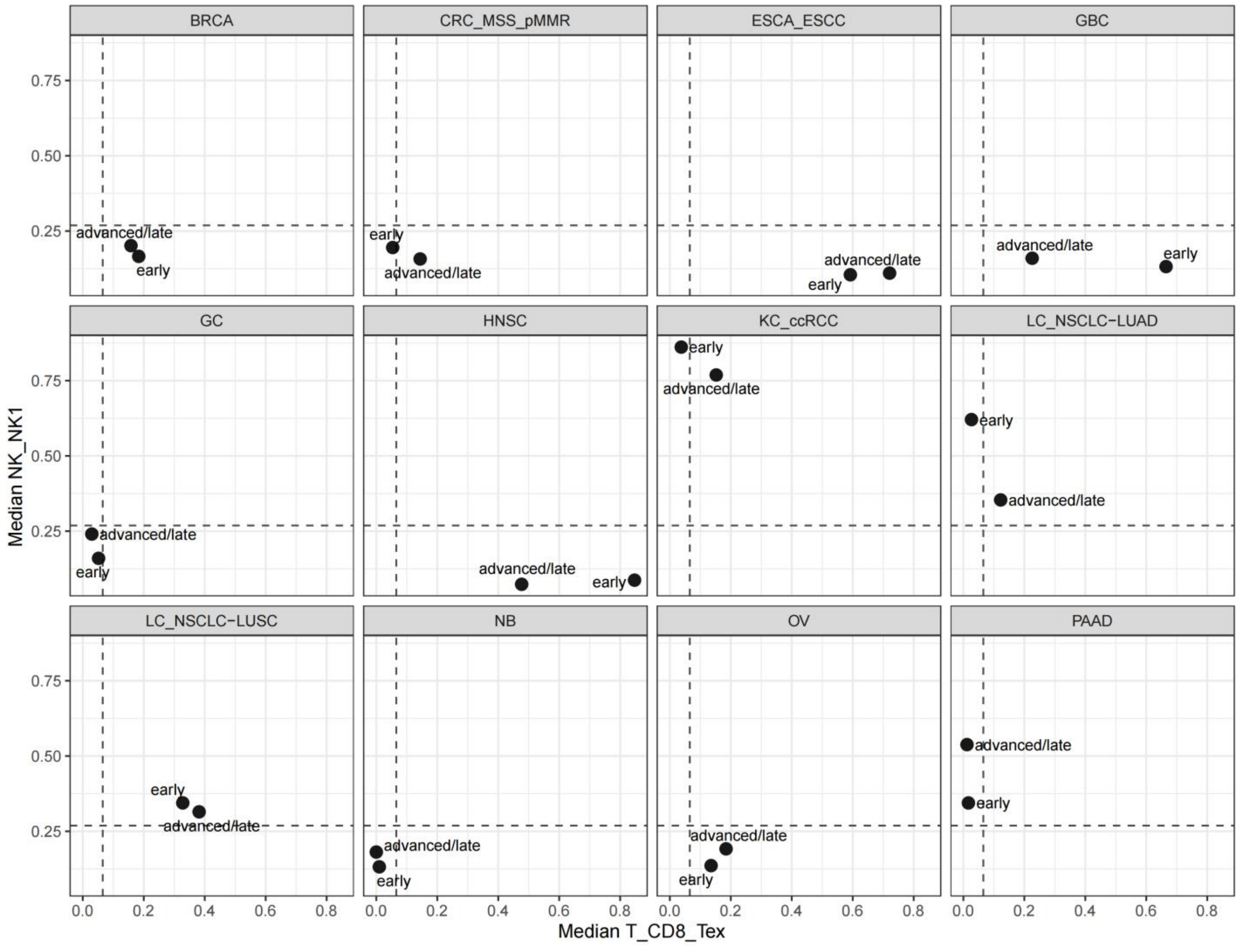


**Fig. S17. Killer divergence stratified by clinical stage.** Each black dot represents the median value for a specific subgroup within that respective cancer. Only cancer types possessing sufficient statistical power (defined as $n \geq 10$ samples for both early and advanced/late stages) were retained. Dashed vertical and horizontal reference lines denote global subset medians. Abbreviation for each malignancy can be found in Table. S1.

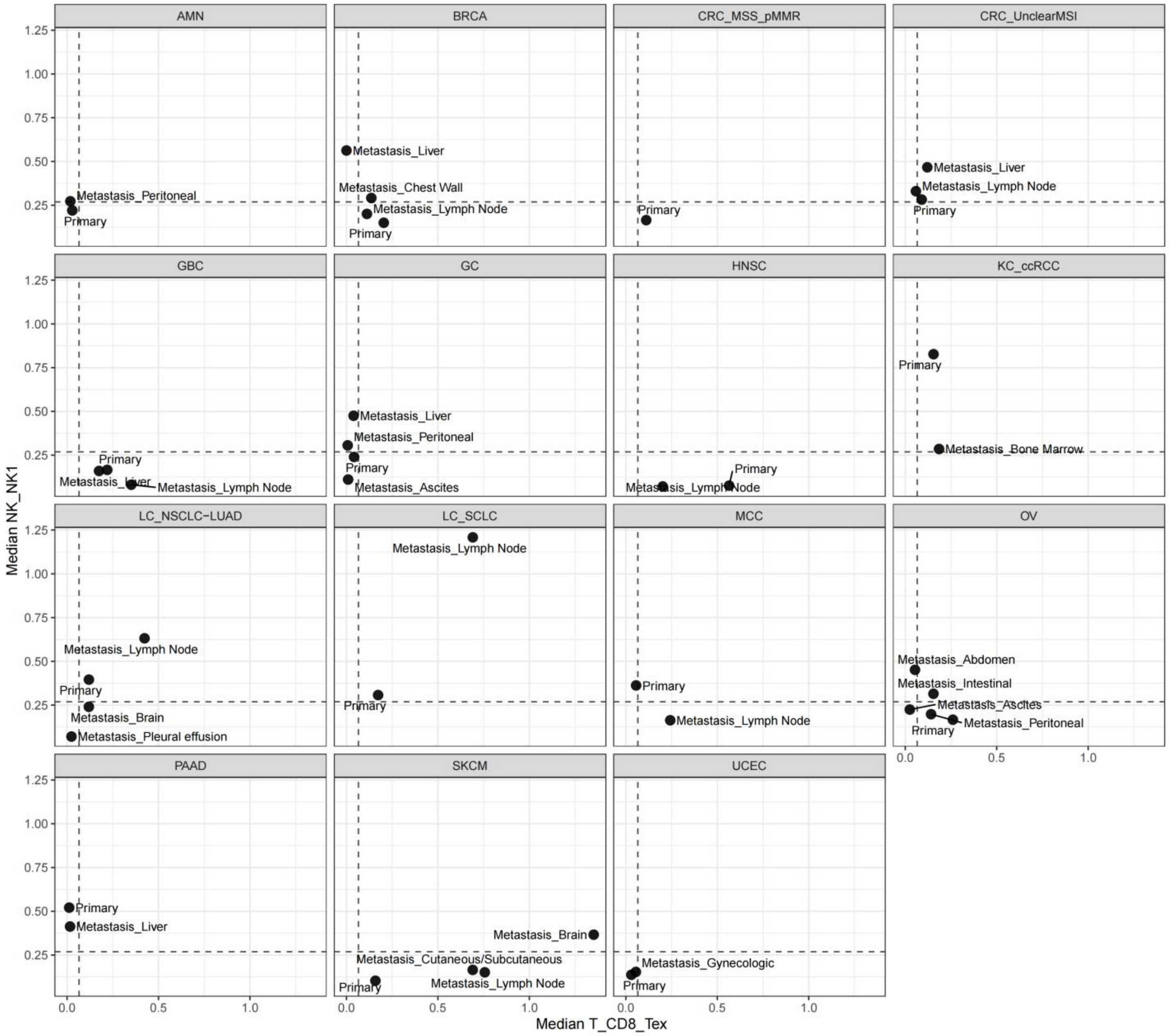


**Fig. S18. Killer divergence stratified by site.** Each black dot represents the median value for a specific subgroup within that respective cancer. To eliminate potential confounding effects driven by clinical stage disparities, specific malignancies (including LUAD, PRAD, LIHC, GC, and BRCA) were restricted exclusively to advanced/late-stage samples. Dashed lines represent global subset medians. Abbreviation for each malignancy can be found in Table. S1.

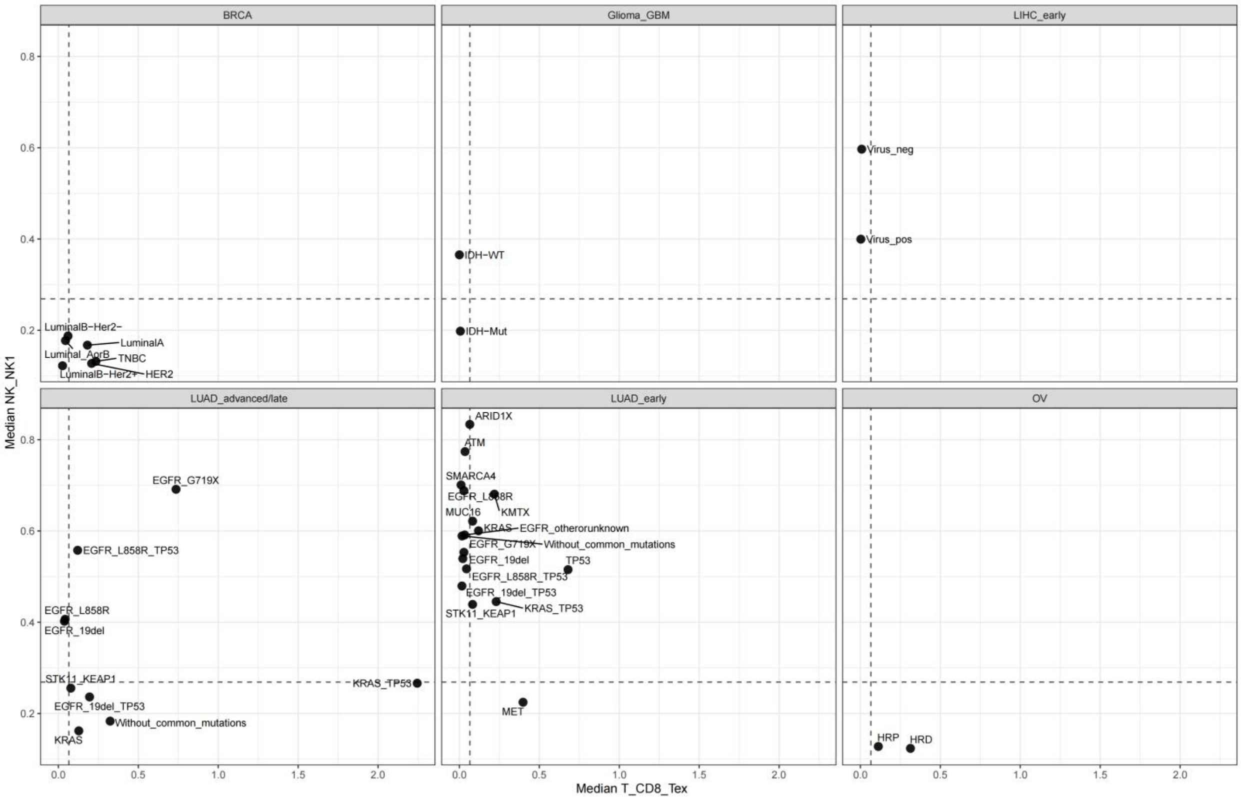


**Fig. S19. Killer divergence stratified by molecular feature.** Each black dot represents the median value for a specific subgroup within that respective cancer. Dashed lines indicate global subset medians. Abbreviation for each malignancy can be found in Table. S1.

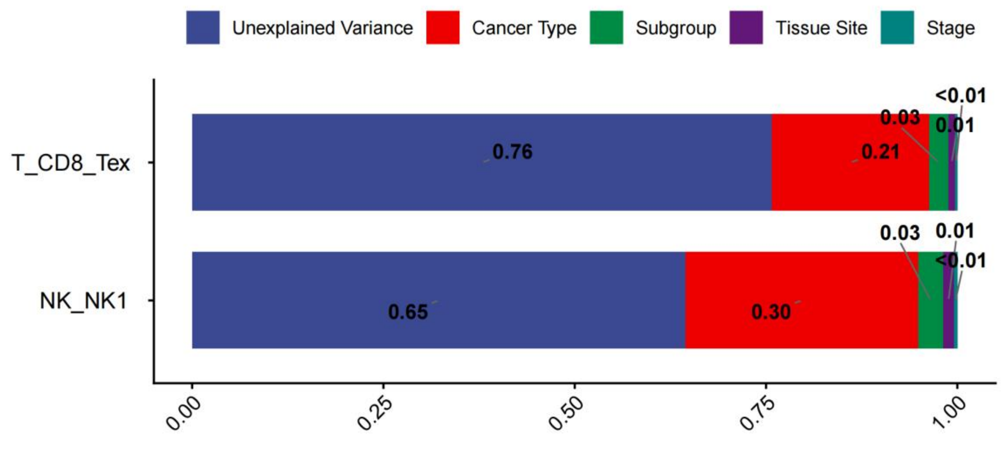


**Fig. S20. Prediction of Tex or NK1 based on clinical variables using multivariate linear model.**

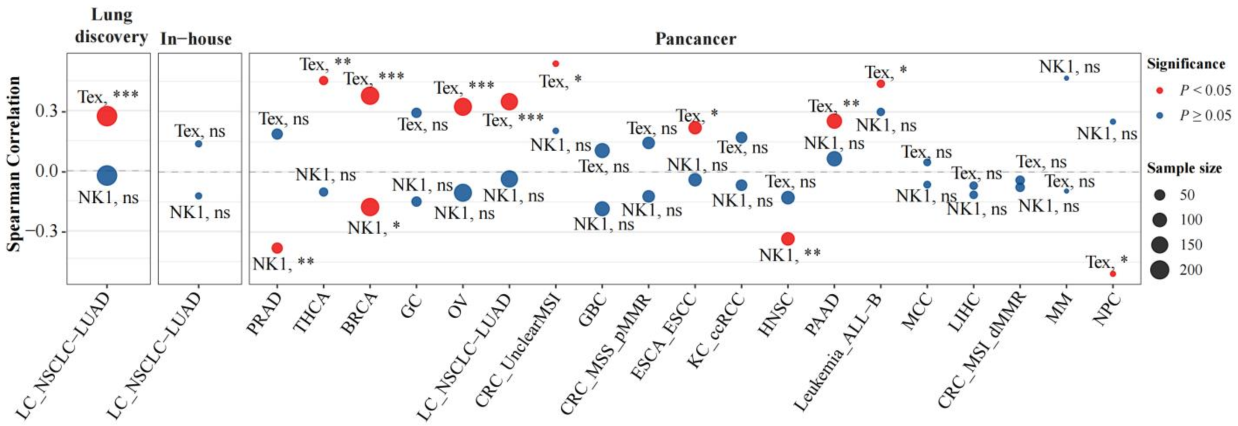


**Fig. S21. Associated of MHC-I score with Tex or NK1.** Cancer type with <15 samples was excluded in this analysis. Node size is proportional to the specific cohort sample size. Node color designates statistical significance (red, $P < 0.05$; blue, $P \geq 0.05$). Adjacent text labels identify the evaluated killer cell subset (Tex or NK1) alongside standard significance thresholds (* $P < 0.05$; ** $P < 0.01$; *** $P < 0.001$; ns, not significant).

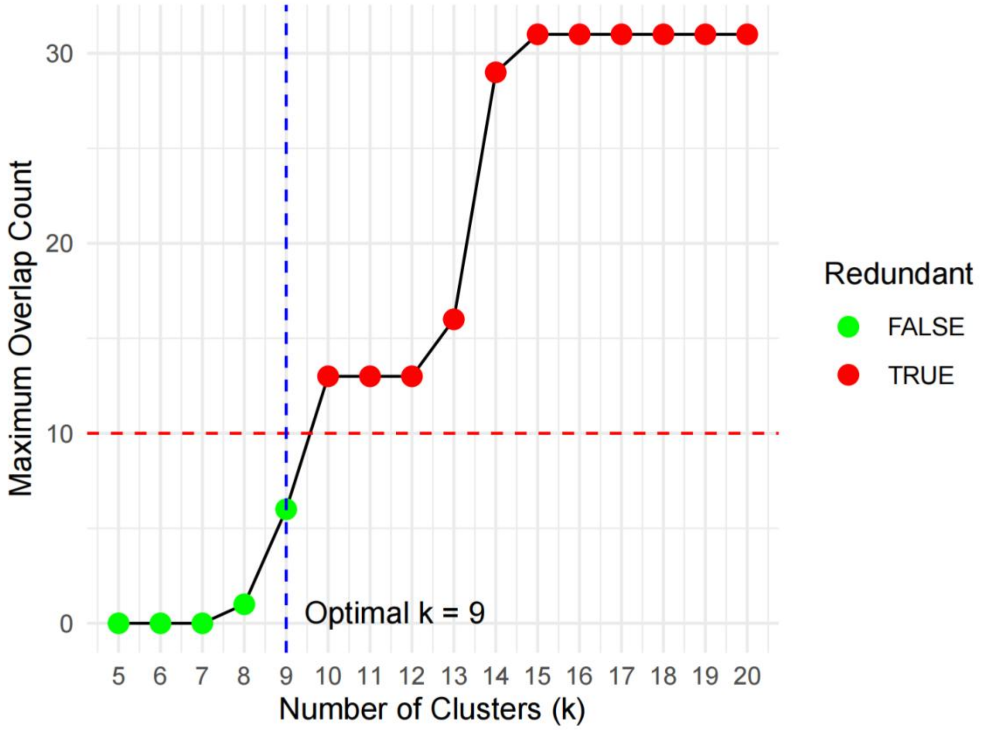


**Fig. S22. Identification of optimal clustering number for malignant meta-programs.** To objectively determine the optimal number of distinct MPs, the hierarchical dendrogram was iteratively partitioned across a testing range of 5 to 20 clusters (*k*). For each *k*, the maximum gene overlap count between any two resultant MP clusters was evaluated. A redundancy threshold was established at a maximum of 10 overlapping top genetic features (denoted by the horizontal dashed red line). Clustering resolutions that satisfy this independence criterion are marked by green nodes (Redundant = FALSE), whereas sub-optimal resolutions generating redundant MPs are marked by red nodes (Redundant = TRUE). The vertical dashed blue line designates $k = 9$ as the optimal clustering resolution, representing the highest number of distinct MPs achievable without violating the genetic overlap threshold.

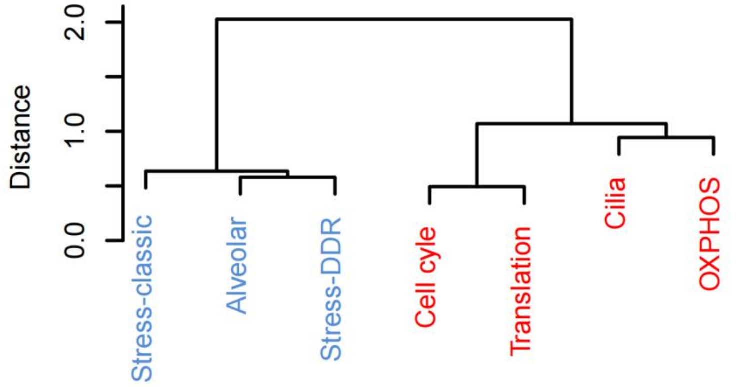


**Fig. S23. Clustering of meta-programs (MP) into MP families.** Dendrogram illustrating the transcriptional relatedness among the tumor meta-programs (MPs). Sample-level MP activity was quantified utilizing a background-corrected control scoring methodology applied to pseudobulk expression of malignant cells. Pairwise distances between individual MPs were calculated using Spearman's rank correlation, followed by unsupervised hierarchical clustering utilizing the Ward.D2 method.

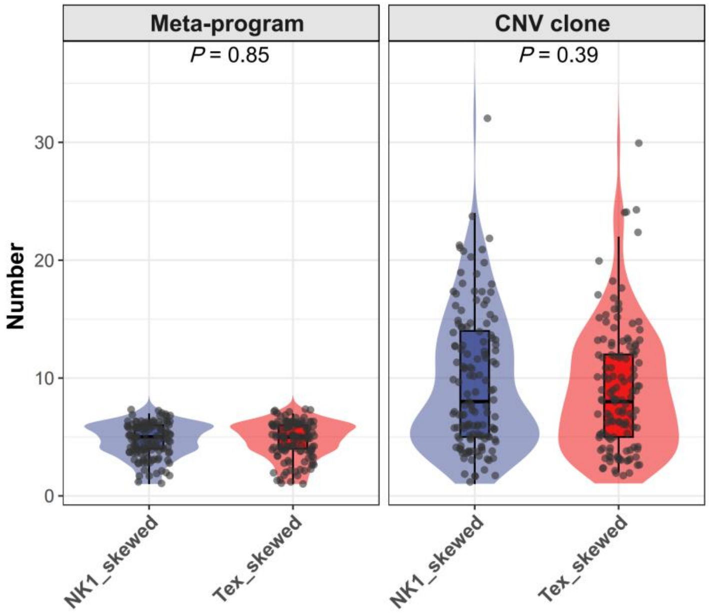


**Fig. S24. Comparison of intratumoral heterogeneity (ITH) levels between killer-skewed groups.** ITH levels were calculated using two strategies: (left) calculating the number of detected meta-programs, and (right) calculating the number of copy number variant clones inferred from inferCNV.

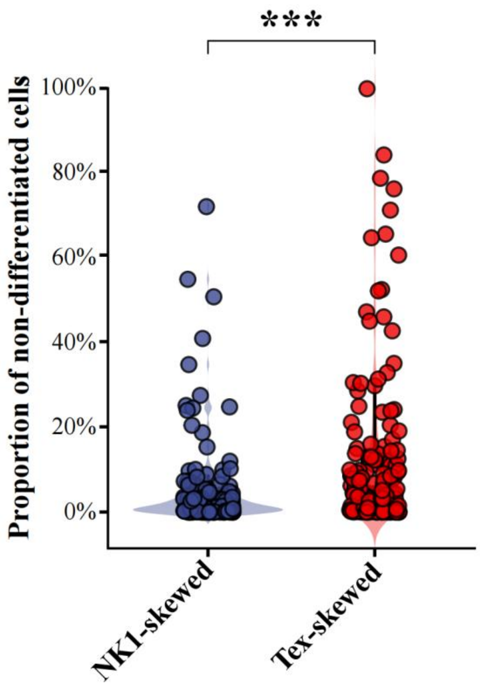


**Fig. S25. Comparison of developmental potential based on CytoTRACE 2.** The proportion of non-differentiated cells was calculated utilizing the CytoTRACE2_Potency classifications.

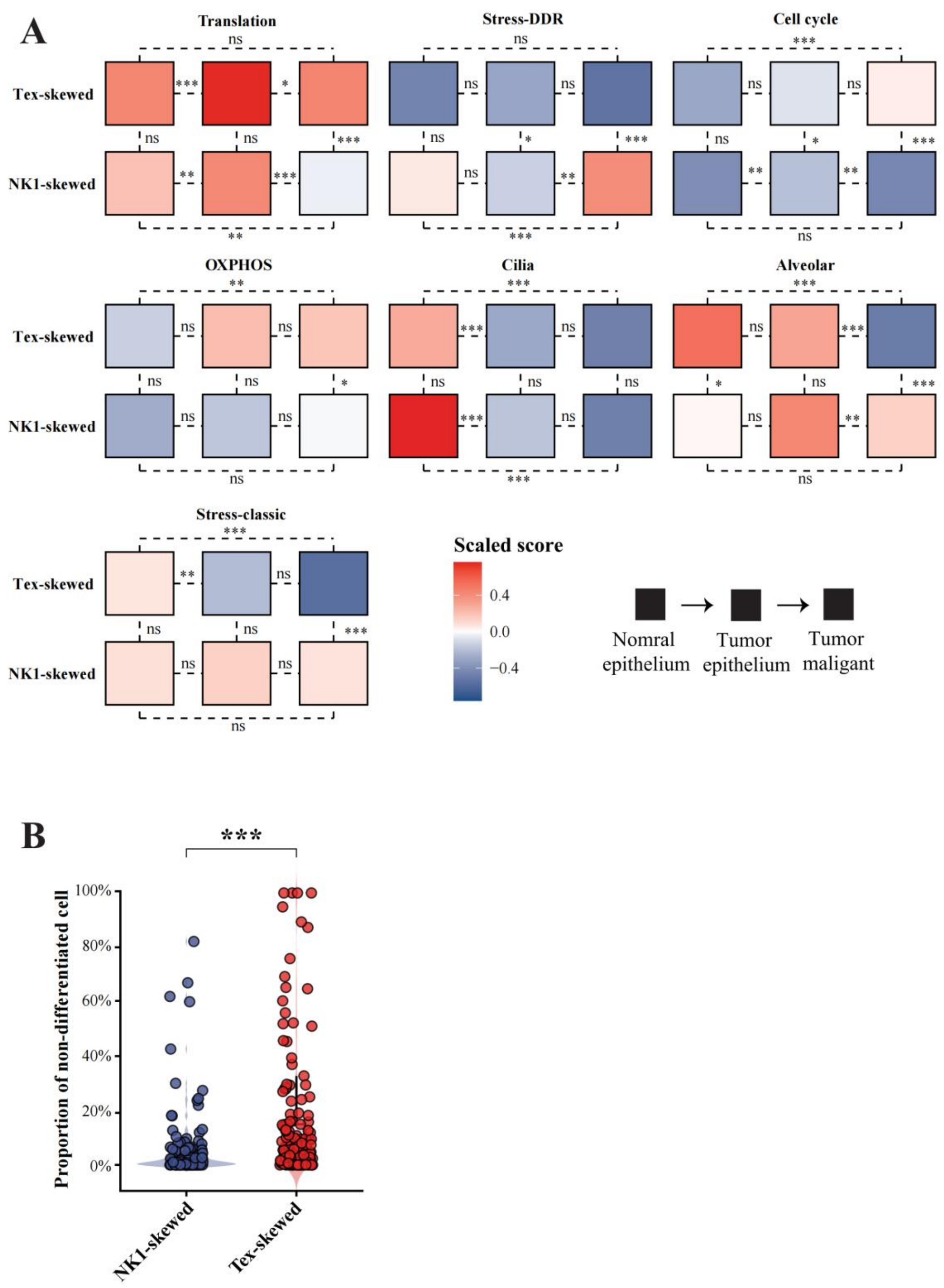


**Fig. S26. Validation of meta-program results when restricting the analysis to tumor cells defined by the overlap of inferCNV and CopyKAT.** This figure is analogous to Fig. 4B and Fig. S26, but it additionally incorporates CopyKAT predictions to establish a more stringent malignant consensus.

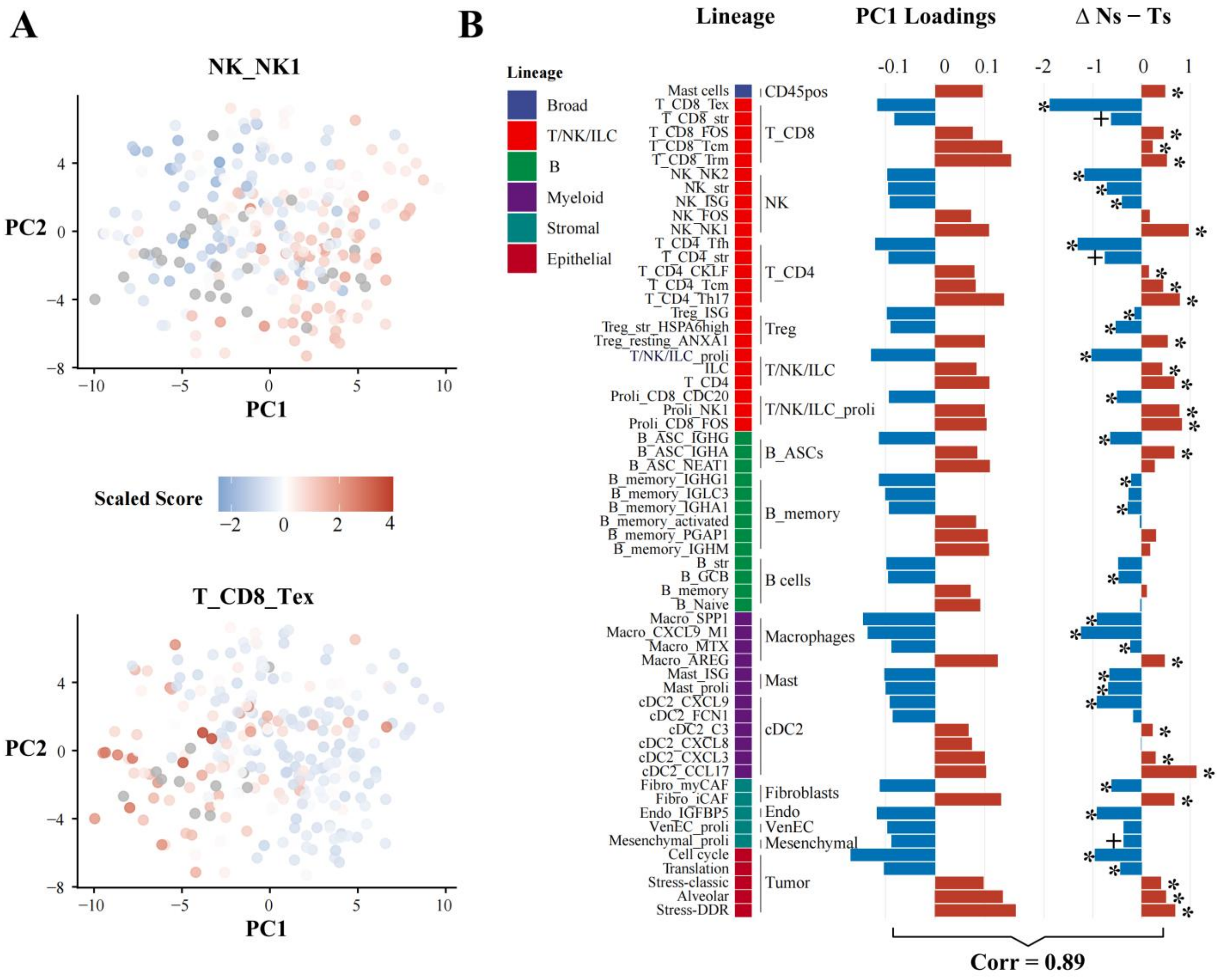


**Fig. S27. Validation of the dominance pattern using lower-granularity cell clustering.** This figure is analogous to Fig. 4C-D, but it utilizes a different PCA input based on 228 lower-granularity cell subsets (see Table. S6). *FDR < 0.05, $^{+}P$<0.05.

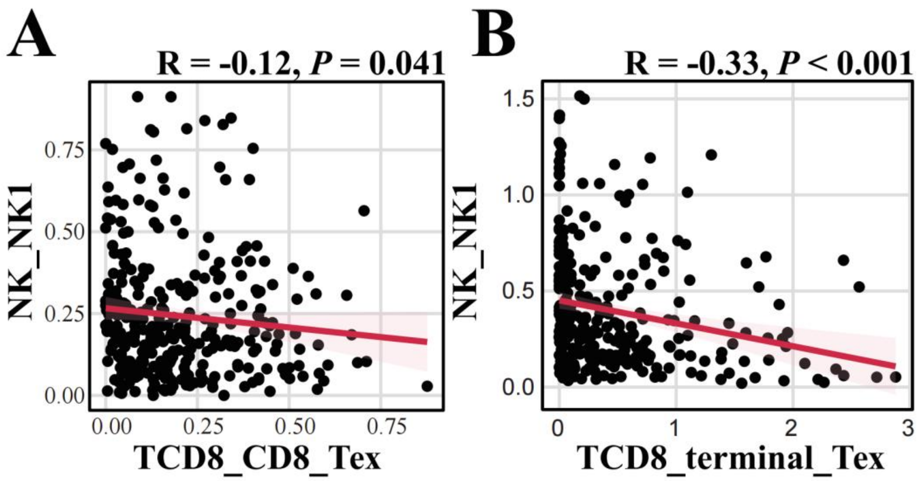


**Fig. S28. Recapitulation of the Tex-NK1 divergence in the ICB scRNA-seq cohort.** (**A**) Negative correlation at the proportional level. (**B**) Further validation utilizing Tex and NK1 signature scores.

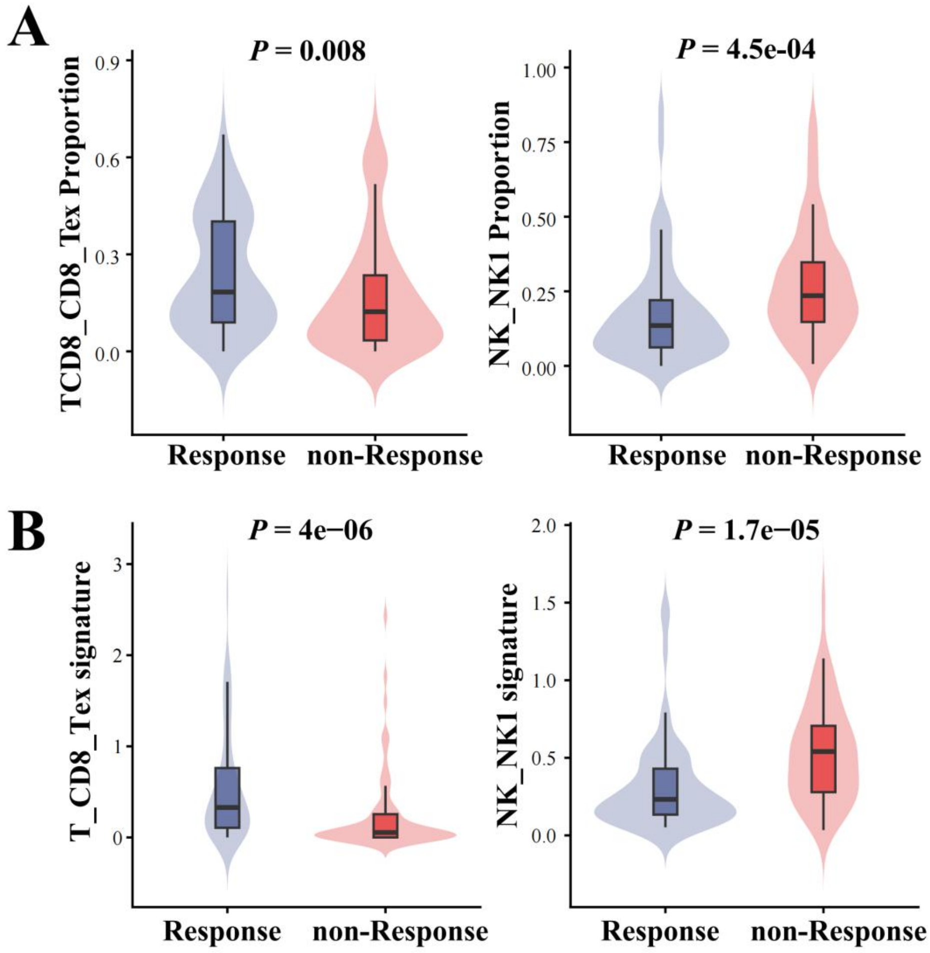


**Fig. S29. Association of killer subsets with ICB response in patients treated with monotherapy.** (**A**) ICB responders exhibited significantly higher Tex and lower NK1 cell proportions compared to non-responders. (**B**) Further validation utilizing Tex and NK1 signature scores.

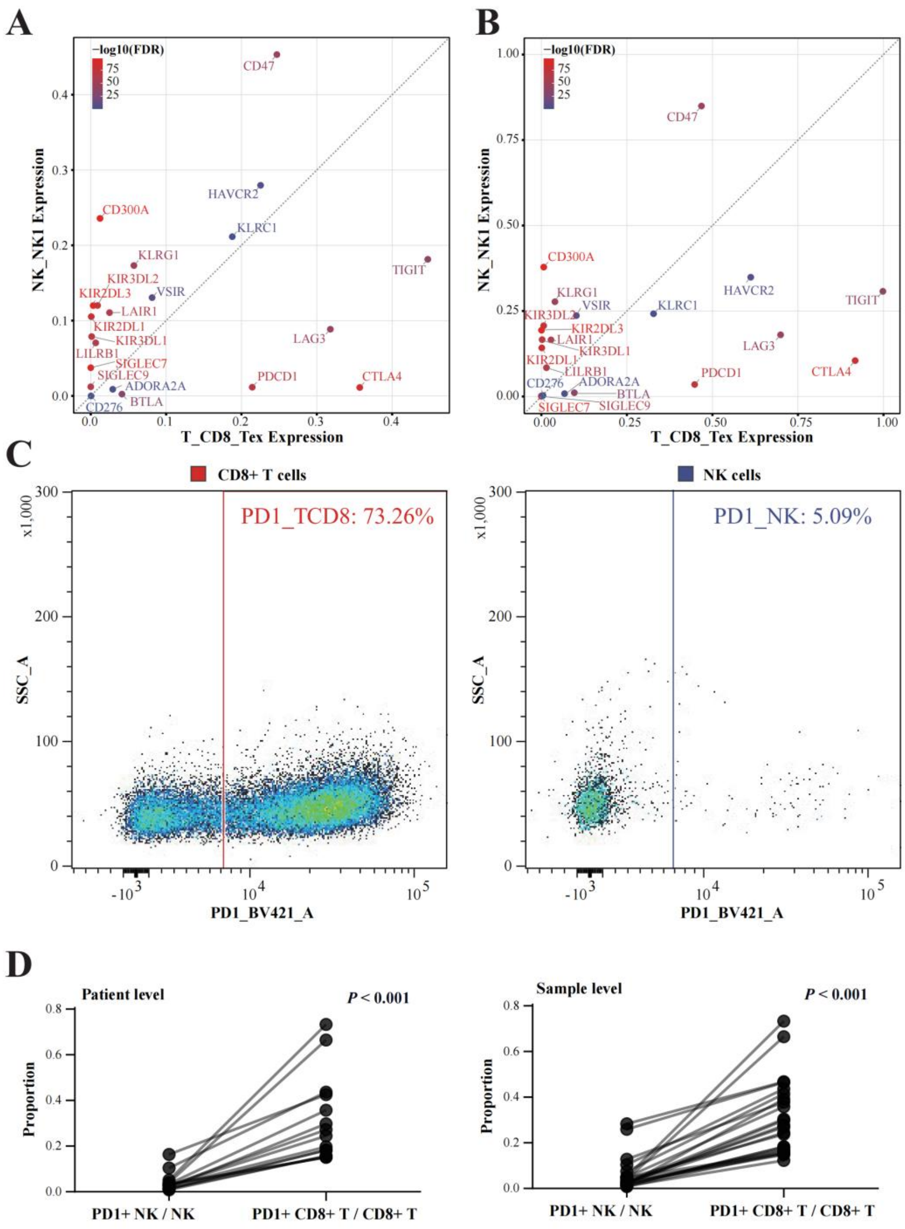


**Fig. S30. Comparison of checkpoint profiles between Tex and NK1 cells.** (**A**) A curated panel of T/NK checkpoints was evaluated to compare NK1 and Tex populations within the lung discovery scRNA-seq cohort. (**B**) Representative spectral flow cytometry plot showing PD-1 expression in CD8$^{+}$ T cells vs NK cells from Sample_8 (Table. S1). (**C**) PD-1 protein expression was compared between the NK1 and Tex subsets via spectral flow cytometry, utilizing a paired Wilcoxon signed-rank test.

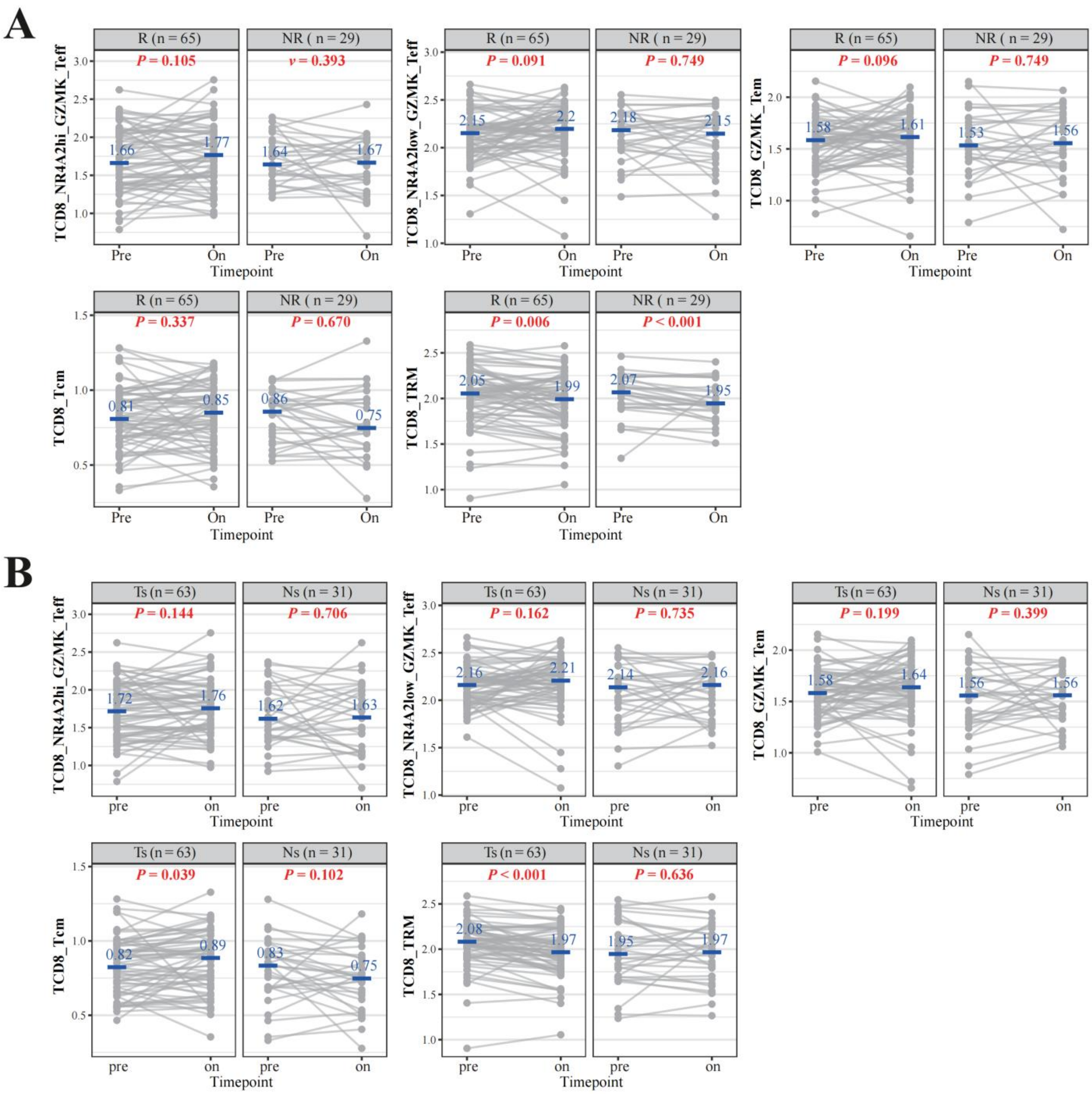


**Fig. S31. Overall CD8⁺ T cell dynamic shifts following ICB.** This figure is related to Fig.5 E-F. Comparisons between matched pre- and on-treatment tumor samples were utilized to evaluate the dynamic shifts of key progenitor memory CD8⁺ T cells. (**A**) Association of these dynamics with clinical response. (**B**) Association of these dynamics with baseline killer profiles.

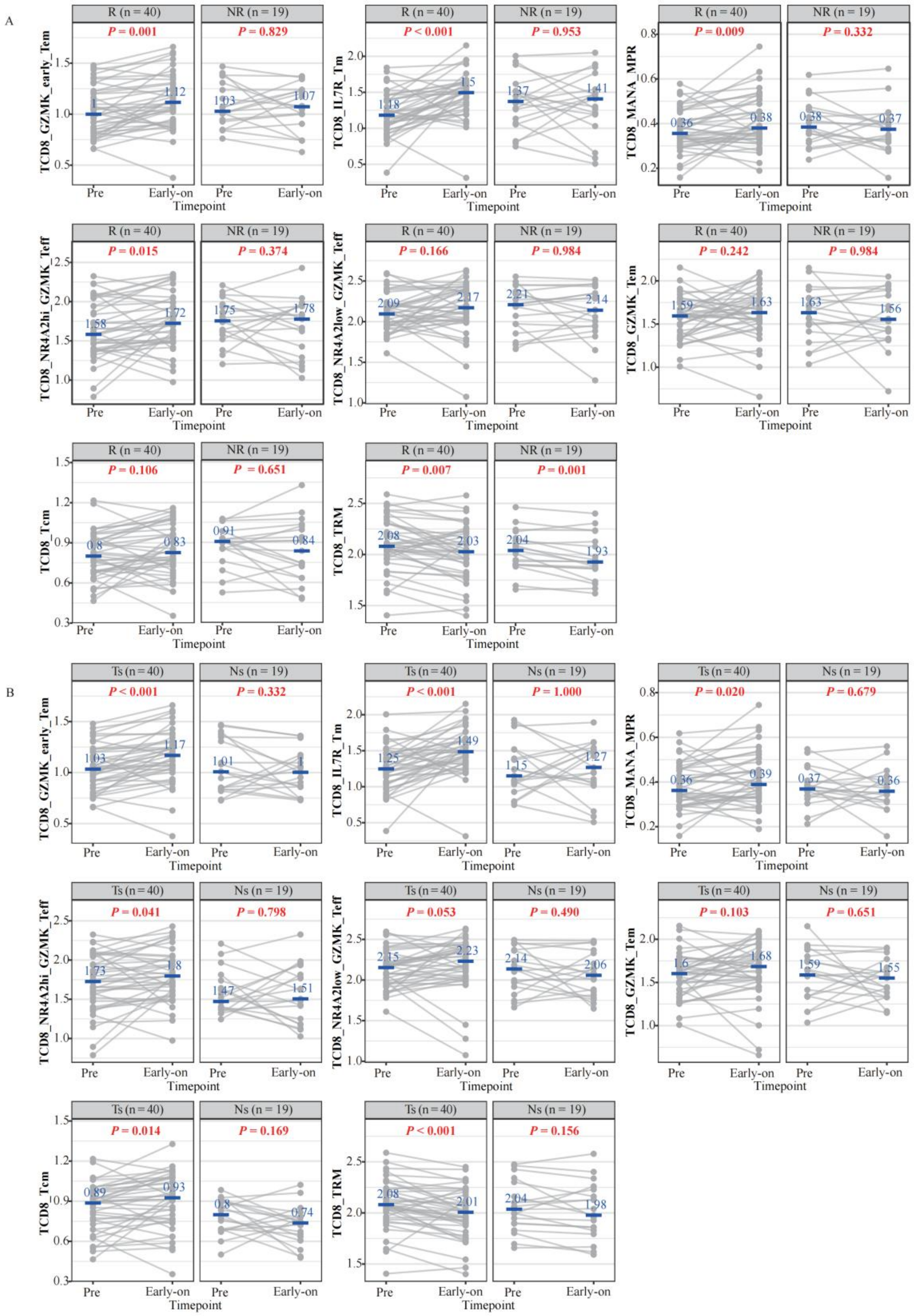


**Fig. S32. Early CD8⁺ T cell dynamic shifts following ICB.** On-treatment samples were stratified into early and late on-treatment groups utilizing a four-week cutoff. Comparisons were performed between matched pre-treatment and early on-treatment tumor samples. (**A**) Association of these dynamics with clinical response. (**B**) Association of these dynamics with baseline killer profiles.

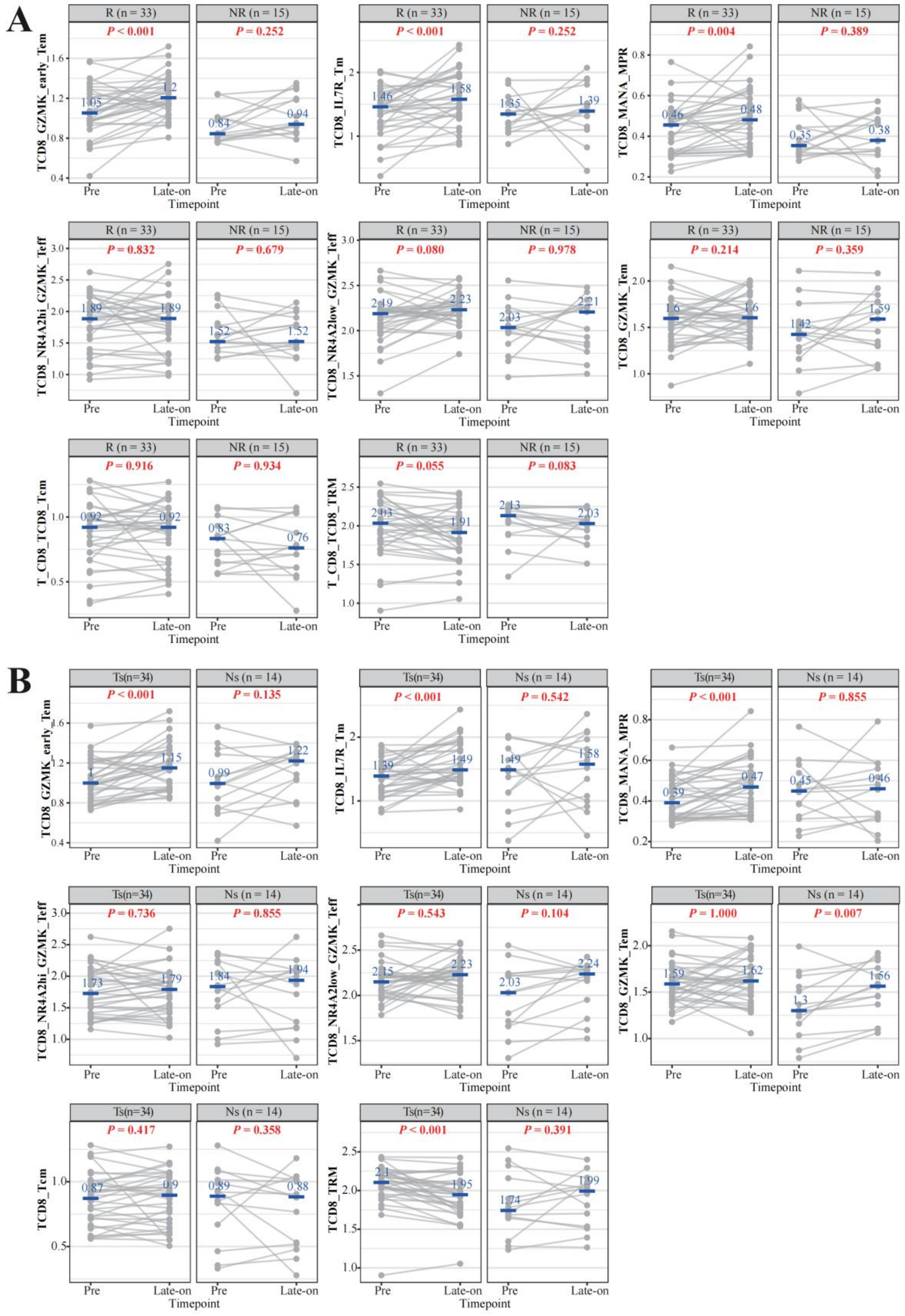

**Fig. S33. Late CD8⁺ T cell dynamic shifts following ICB.** On-treatment samples were stratified into early and late on-treatment groups utilizing a four-week cutoff. Comparisons were performed between matched pre-treatment and late on-treatment tumor samples. (**A**) Association of these dynamics with clinical response. (**B**) Association of these dynamics with baseline killer profiles..

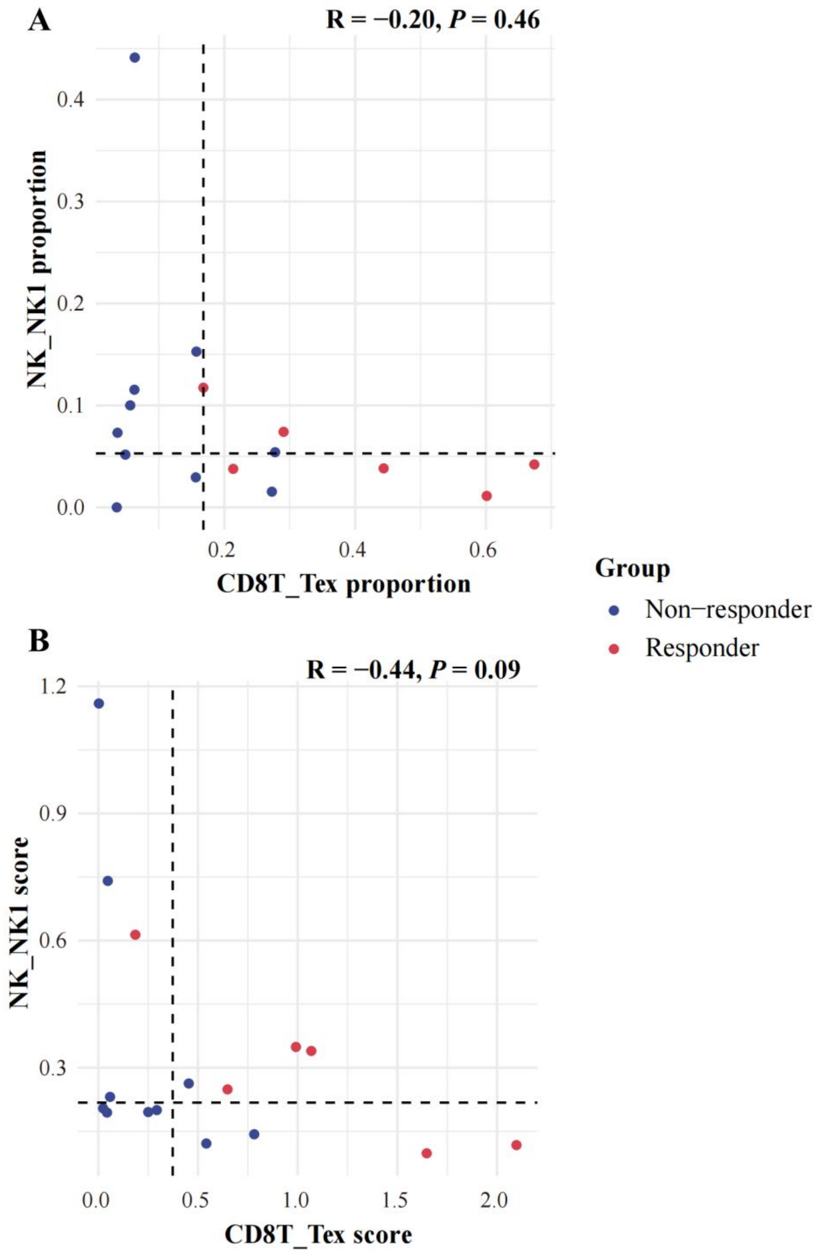


**Fig. S34. Association of killer divergence with adoptive cellular therapy response.** This cohort included 18 pre-treatment samples from melanoma and GBM patients. (**A**) A negative correlation was observed between Tex and NK1, with clinical responders enriched within the Tex-skewed population; however, statistical significance was not reached, likely due to the limited sample size. (**B**) Further validation utilizing Tex and NK1 signature scores.

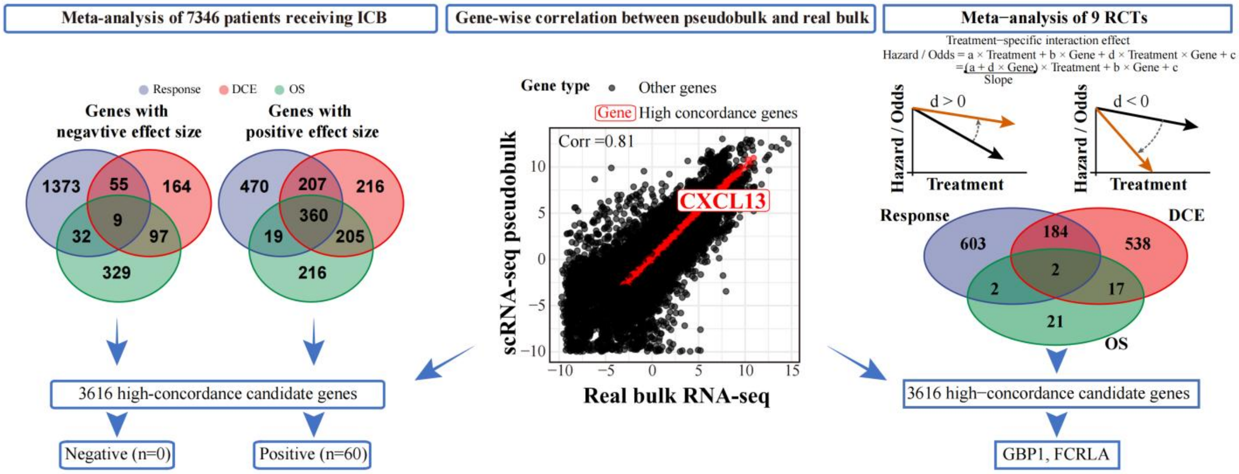


**Fig. S35. Meta-analysis pipeline for identifying a robust predictive signature in bulk RNA-seq ICB cohort.** (**Left**) Venn diagrams summarizing a random-effects meta-analysis across 7,346 patients receiving ICB, displaying the intersection of genes significantly associated with objective response rate, disease control endpoints (DCE), and overall survival (OS). Filtering this intersection against 3,616 cross-modality high-concordance candidate genes yielded a final 60-gene robust ICB efficacy signature; no negative predictors passed these strict criteria. (**Middle**) Scatter plot evaluating the cross-modality transcriptomic concordance utilized for candidate gene filtering. Median gene expression profiles from the aggregated scRNA-seq pseudobulk lung discovery cohort (y-axis) are plotted against actual bulk RNA-seq expression levels from the TCGA-LUAD cohort (x-axis), demonstrating high global concordance (Spearman's corr = 0.81). Genes exhibiting the highest cross-modality conservation—defined as falling within the bottom quartile of the absolute quantitative difference between modalities after excluding the lowest-expressing decile—are highlighted in red (e.g., *CXCL13*), establishing the 3,616 high-concordance candidate pool. (**Right**) Analytical framework and Venn diagram for identifying genes that predict ICB-specific therapeutic benefit over conventional therapies. (Top) Schematic of the treatment-by-gene interaction models (Cox proportional hazards for time-to-event endpoints; logistic regression for binary response). If the interaction coefficient $d$ is positive, higher gene expression will flatten the slope between treatment and hazard/odds, indicating a reduced advantage of ICB over conventional therapy. If $d$ is negative, higher gene expression will steepen the slope between treatment and hazard/odds, indicating an increased advantage of ICB over conventional therapy. (Bottom) Venn diagram intersecting genes that demonstrate significant, consistent ICB-specific benefit across nine randomized controlled trials evaluated across response, DCE, and OS endpoints. Subsequent intersection with the high-concordance candidate pool identified two $d < 0$ genes, *GBP1* and *FCRLA*, as predictors of ICB-over-control benefit.

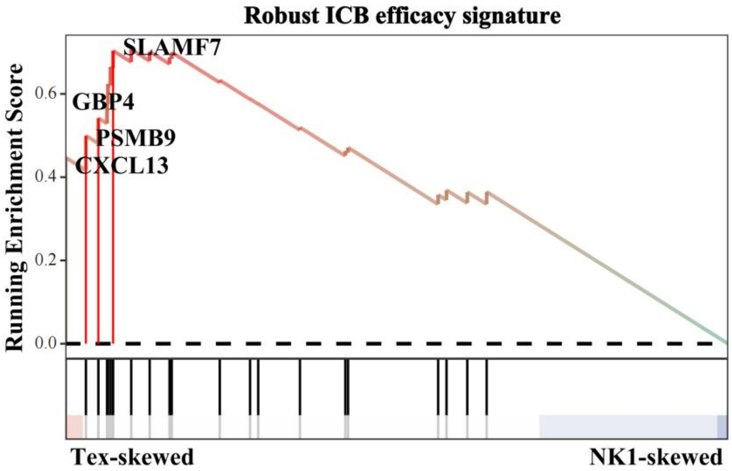


**Fig. S36. Association of pseudobulk profiles with a robust ICB efficacy signature derived from bulk RNA-seq monotherapy cohorts.** This plot is analogous to gene set enrichment analysis in Fig. 5H, but it is restricted exclusively to patients receiving single-agent ICB therapy.

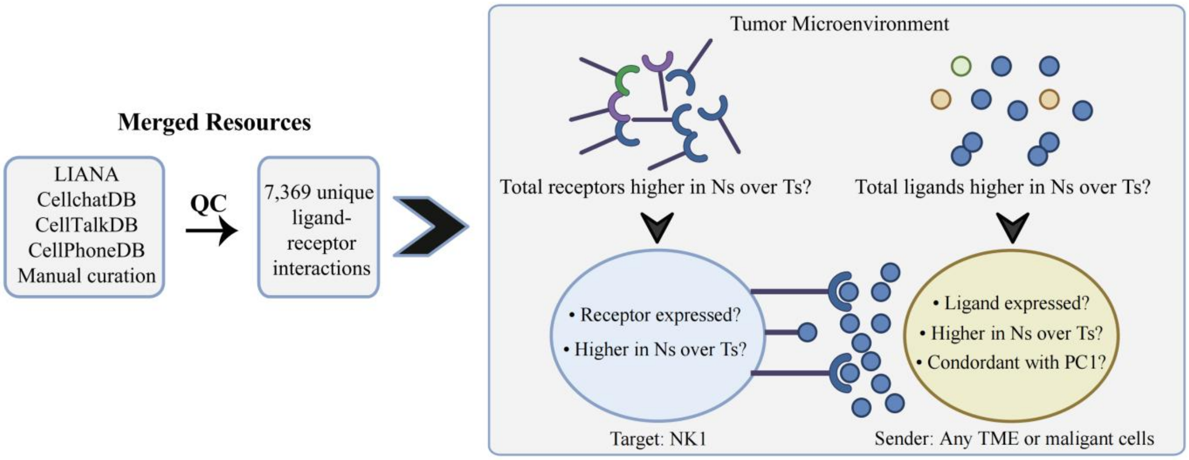


**Fig. S37. Identification of dominant ligand-receptor interactions driving NK1 evasion.** A candidate receptor was required to be expressed in at least 5% of the NK1 cells. The receptor must exhibit significantly higher expression within the NK1 cells of NK1-skewed (Ns) tumors compared to Tex-skewed (Ts) tumors. The corresponding ligand must demonstrate significantly higher expression across the global, unsorted pseudobulk profile in Ns tumors. To confirm that the global ligand upregulation is not a bulk artifact, the specific cellular origin of the ligand must be identified. Any candidate interacting ligand was required to meet this 5% expression threshold within its respective source cell lineage. The ligand must be significantly upregulated within the isolated pseudobulk profile of at least one specific cell type. Ideally, these source cellular subsets should also align with the Ns direction along the PC1 axis.

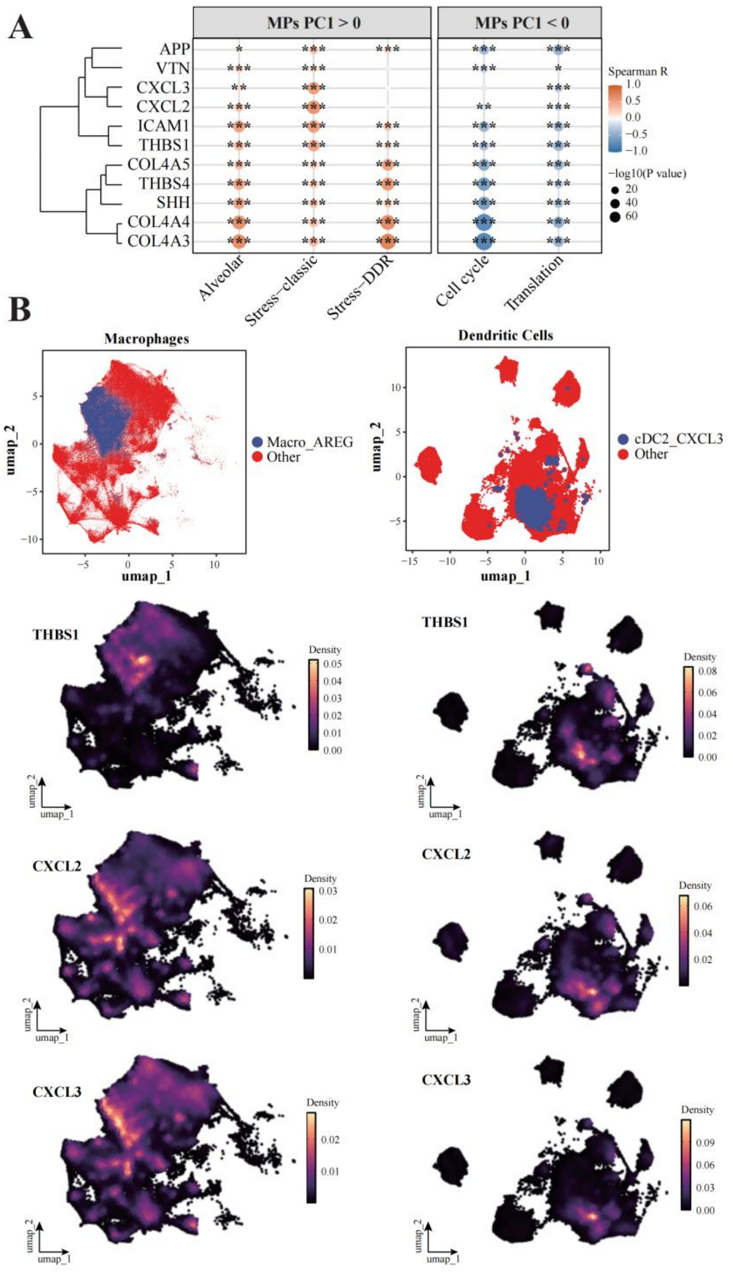


**Fig. S38. Malignant meta-programs and myeloid subsets express NK1-suppressive ligands.** (**A**) Spearman rank correlation between the expression of top candidate NK1-interacting ligands and the activity scores of distinct tumor-intrinsic meta-programs (MPs) in tumor cells. The MPs are segregated into those aligned with the NK1-skewed pole (MPs PC1 > 0: Alveolar, Stress-classic, and Stress-DDR) versus the Tex-skewed pole (MPs PC1 < 0: Cell cycle and Translation). Node color designates the Spearman correlation coefficient. Node size is proportional to the statistical significance. Asterisks explicitly denote statistical significance thresholds (* $P < 0.05$; ** $P < 0.01$; *** $P < 0.001$). (**B**) Cellular origins of the ligands within the myeloid compartment. (Top) Uniform manifold approximation and projection (UMAP) plots of the macrophage (left) and dendritic cell (right) lineages, specifically highlighting the spatial localization of the Macro_AREG and cDC2_CXCL3 subsets relative to all other lineage constituents. (Bottom) Kernel density estimations mapped onto corresponding UMAP coordinates.

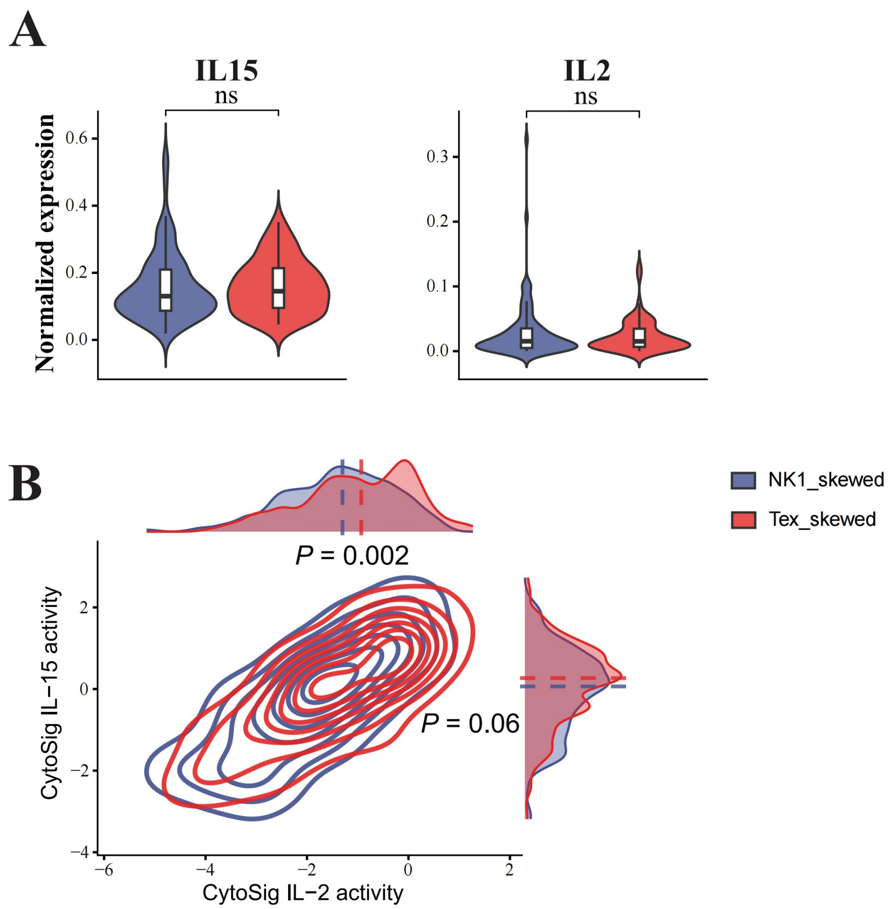


**Fig. S39. Lower cytokine activity in NK1 cells in NK1-skewed versus Tex-skewed groups.** (**A**) Expression of the IL-15 and IL-2. (**B**) Contour plot coupled with density histograms illustrating the predicted cytokine signaling pathway activities by CytoSig for IL-2 and IL-15. The internal horizontal lines represent the median, and the box edges denote the interquartile ranges. * $P < 0.05$; ** $P < 0.01$; *** $P < 0.001$; ns, not significant.
